# Polyploid cardiomyocytes define disease-specific transcriptional states in the mammalian heart

**DOI:** 10.64898/2026.01.31.701472

**Authors:** Paul Kiessling, Mehdi Joodaki, Daryna Pikulska, Emilia Scheidereit, Giulia Cesaro, Mayra Ruiz, Damin Kühn, Kai Peng, Osman Goni, Sebastian Foecking, Samaneh Samiei, Xian Liao, Kai Li, Zihao Feng, Delin Wang, Lampros Mavrommatis, Anna-Maria Vllaho, Merwan Rombach, Giulia Cucinella, Mingbo Cheng, Till Lautenschläger, David Rodriguez Morales, Rogier J. A. Veltrop, Leon J. Schurgers, Moritz von Scheidt, Barbara Mara Klinkhammer, Florian Kahles, Jennifer Kranz, Aitor Aguirre, Anne Loft, Niklas Klümper, Markus Eckstein, Thomas Seidel, Giancarlo Forte, Rik Westland, Man Zhang, Heng Zhao, Feng Ren, Sikander Hayat, Junedh Amrute, Benjamin Kopecky, Rebekka K. Schneider, Hind Medyouf, Pierre-Louis Tharaux, Peter Boor, Rafael Kramann, Linda W. van Laake, Annelotte Vos, Bernadette S. de Bakker, Jermo Hanemaaijer-van der Veer, Kory Lavine, Nikolaus Marx, Karin Klingel, Michael T. Schaub, Stefanie Dimmeler, Alex Zhavoronkov, Hendrik Milting, Ivan G. Costa, Christoph Kuppe

**Affiliations:** Division of Nephrology and Clinical Immunology, RWTH Aachen University, Medical Faculty, Aachen, Germany; Institute of Computational Genomics, RWTH Aachen University, Medical Faculty, Aachen, Germany; Center for Computational Life Sciences, RWTH Aachen University, Aachen, Germany; Department of Information Engineering, University of Padova, Padova, Italy; Department of Computer Science, RWTH Aachen University, Aachen, Germany; Institute of Cardiovascular Regeneration, Goethe University Frankfurt, Frankfurt am Main, Germany; German Center for Cardiovascular Research (DZHK), Partner Site Rhine-Main, Germany; Cardio-Pulmonary Institute (CPI), Frankfurt, Germany; Department of Biochemistry, Cardiovascular Research Institute Maastricht (CARIM), Maastricht University, Maastricht, The Netherlands; Department of Cardiology, Division of Heart and Lungs, University Medical Center Utrecht, Utrecht University, Heidelberglaan 100, 3584 CX Utrecht, The Netherlands; Regenerative Medicine Center Utrecht and Circulatory Health Research Center, University Medical Center Utrecht, Utrecht University, Heidelberglaan 8, 3584 CS Utrecht, The Netherlands; LMNAcardiac.org, LMNA Patient Organization, Vredehofstraat 13761, HA Soest, The Netherlands; Netherlands Heart Institute, Holland Heart House, Moreelsepark 1, 3511 EP Utrecht, The Netherlands; Department of Cardiology, TUM Klinikum Deutsches Herzzentrum, Technical University Munich, Munich, Germany; Deutsches Zentrum für Herz- und Kreislaufforschung (DZHK), Partner Site Munich Heart Alliance, Munich, Germany; Institute of Pathology, RWTH Aachen University Hospital, Aachen, Germany; Department of Internal Medicine I-Cardiology, University Hospital Aachen, RWTH Aachen University, Aachen, Germany; Department of Urology and Pediatric Urology, University Hospital RWTH Aachen, Aachen, Germany; Department of Urology and Kidney Transplantation, Martin-Luther-University, Halle, Germany; Institute for Quantitative Health Science and Engineering, Michigan State University, East Lansing, MI, USA; Department of Biomedical Engineering, Michigan State University, East Lansing, MI, USA; Center for Functional Genomics and Tissue Plasticity (ATLAS), Department of Biochemistry and Molecular Biology, University of Southern Denmark (SDU), Odense, Denmark; The Novo Nordisk Foundation Center for Genomic Mechanisms of Disease, Broad Institute of MIT and Harvard, Cambridge, MA, USA; Department of Urology and Pediatric Urology, University Hospital Bonn, Bonn, Germany; Institute of Experimental Oncology, University Hospital Bonn, Bonn, Germany; Institute of Pathology, Universitätsklinikum Erlangen, Friedrich-Alexander-Universität Erlangen-Nürnberg (FAU), Erlangen, Germany; Comprehensive Cancer Center Erlangen-EMN (CCC ER-EMN), Erlangen, Germany; Comprehensive Cancer Center Alliance WERA (CCC WERA), Erlangen, Germany; Bavarian Cancer Research Center (BZKF), Erlangen, Germany; Institute of Cellular and Molecular Physiology, Friedrich-Alexander-Universität Erlangen-Nürnberg, Erlangen, Germany; International Clinical Research Center (ICRC), St Anne’s University Hospital, Brno, Czech Republic; Faculty of Medicine, Masaryk University, Brno, Czech Republic; School of Cardiovascular and Metabolic Medicine and Sciences, King’s College London, London, UK; Department of Pediatric Nephrology, Amsterdam UMC-Emma Children’s Hospital, Location University of Amsterdam, Meibergdreef 9, 1105 AZ, Amsterdam, The Netherlands; Insilico Medicine Shanghai Ltd., Shanghai, China; Insilico Medicine Hong Kong Ltd., Hong Kong SAR, China; Cardiovascular Research Institute and Department of Medicine, Icahn School of Medicine at Mount Sinai, New York, NY, USA; Windreich Department of Artificial Intelligence and Human Health, Icahn School of Medicine at Mount Sinai, New York, NY, USA; Center for Cardiovascular Research, Division of Cardiology, Department of Medicine, Washington University in St. Louis School of Medicine, St. Louis, MO, USA; Department of Medicine, Division of Cardiology, University of Colorado Anschutz Medical Campus, Aurora, CO, USA; Department of Developmental Biology Erasmus Medical Center Rotterdam, The Netherlands; Oncode Institute Erasmus Medical Center Cancer Institute Rotterdam, The Netherlands; Department of Cell and Tumor Biology, Faculty of Medicine University Hospital RWTH Aachen, Aachen, Germany; Department of Hematology, Oncology, Hemostaseology, Stem Cell Transplantation, Medical Faculty, Center for Integrated Oncology (CIO), RWTH Aachen University, Aachen, Germany; Université Paris Cité, INSERM, Paris Cardiovascular research Center (PARCC), Paris, France; Department of Internal Medicine, Nephrology and Transplantation, Erasmus Medical Center, Rotterdam, The Netherlands; Department of Cardiology, Division of Heart & Lungs, University Medical Centre Utrecht and Utrecht University, Utrecht, The Netherlands; Department of Pathology, University Medical Center Utrecht, Utrecht, The Netherlands; Amsterdam UMC location University of Amsterdam, Dept. of Obstetrics and Gynecology, Amsterdam, The Netherlands; Amsterdam Reproduction and Development Research Institute, Amsterdam, The Netherlands; Cardiopathology, Institute for Pathology and Neuropathology, University Hospital Tübingen, Tübingen, Germany; Department of Biology, RWTH Aachen University, Aachen, Germany; Insilico Medicine US Inc., Cambridge, MA, USA; Heart- and Diabetescenter NRW, Erich and Hanna Klessmann-Institute for Cardiovascular Research and Development, University Hospital of the Ruhr University Bochum, Bad Oeynhausen, Germany

## Abstract

The adult mammalian heart has a limited regenerative capacity. Following injury, cardiomyocytes undergo a hypertrophic response accompanied by polyploidization, which has been described as a barrier to proliferation and regeneration of the heart^1,2^. However, the unique molecular programs of polyploidy, or genome multiplied cardiomyocytes, and their influence on the disease-related myocardial remodelling process remains unclear. Here, we integrate single-nuclei and high-resolution spatial multi-omics across human, rat, and mouse hearts to define novel cardiac cell states and their tissue niches in ischemic and non-ischemic heart disease. Computational analysis across scales allowed us to generate detailed networks of the cardiac tissue remodelling process as well as tissue and sub-cellular environments uniquely enriched in polyploid cardiomyocytes or their diploid origins. We identify a conserved, dichotomous transcriptional program distinguishing diploid from polyploid cardiomyocytes. Polyploid cardiomyocytes demonstrated rewired metabolic and chromatin-remodeling transcriptional programs and recapitulate the gene signature of immature human fetal cardiomyocytes. Notably, we observe that polyploid cardiomyocytes—rather than the general myocyte population—are the primary sites of enrichment for major heart-failure drug targets, including the mineralocorticoid, β1-adrenergic, and glucagon-like peptide-1 receptors. Based on our cross-species dataset we further identified TNIK, a Wnt-pathway regulator expressed in polyploid cardiomyocytes across species, as a potential therapeutic target and demonstrate that pharmacological TNIK inhibition improves cardiac function after myocardial infarction in rats. Together, this species-spanning, disease-resolved study redefines cardiomyocyte heterogeneity in heart disease and suggests a therapeutic path to heart failure treatment by targeting polyploid cardiomyocytes.

## Results

Heart failure represents a major global health burden, affecting nearly half a billion individuals worldwide and causing more deaths than most forms of cancer^3^. Despite significant treatment advancements, current therapies typically address the symptoms and systemic consequences of heart failure, but do not directly target the pathways driving maladaptive remodeling or those that restrict cardiac regeneration^4^. The human heart possesses a minimal intrinsic regenerative capacity^5^ compared to that of other species such as zebrafish^6^ and neonatal mice^7^. Human cardiomyocytes (CMs) originate from proliferative mesodermal precursors during embryogenesis but rapidly lose their regenerative capacity shortly after birth, coinciding with a metabolic shift from glycolysis to fatty acid oxidation and the initiation of polyploidization^1,8,9^. The adult human heart predominantly responds to injury with maladaptive remodeling, stress-associated gene activation, concomitant fibrosis, and progressive functional loss^10,11^. This stress response is typically associated with upregulated expression of the genes NPPB and NPPA, which are co-regulated by an evolutionarily conserved enhancer cluster and induced by mechanical stress^12,13^. Recent studies suggest that metabolic perturbations, specifically inhibition of fatty acid oxidation, induce a fetal cell state shift in CMs and can induce cardiomyocyte proliferation and regeneration highlighting the influence of metabolic states on cardiomyocyte cell state plasticity^14^. It remains unclear whether additional CM cell states, beyond the canonical NPPB/NPPA stress-response, drive cardiac tissue remodelling.

Polyploidization, characterized by increased genome copy numbers or multinucleation in CMs, emerges postnatally as the heart matures^15^. It represents a fundamental cell growth principle in biology, utilized by different organisms and is common in plants and salivary glands of insects^16–18^. Polyploid cell types can be found in various tissues, including heart^19^, muscle^20^, liver^21^ and pancreas^22^ and have been suggested to play a role in tissue repair and regeneration^21^. While several mechanisms like cell fusion, mitotic slippage, and endocycling have been identified as polyploid mechanisms, endocycling, i.e. rounds of DNA replication without cell division, has been identified as the major mechanism in the heart, which is particularly linked to mitochondrial function^23^. Polyploidization has been hypothesized to serve as a barrier to CM proliferation, thereby limiting cardiac regeneration^2^. This appears paradoxical given that polyploidy is beneficial in other organs, such as the liver^24^, where it supports tissue regeneration^24^ and protects against genotoxic stress^25,26^. Despite extensive research^27^, the functional and molecular distinctions between diploid and polyploid CMs in the mammalian heart, especially regarding their disease-modifying relevance in the human heart, have remained elusive.

Here, we present a single-cell and spatial multi-omics analysis of over 4.5 million cardiac nuclei from healthy and diseased hearts across 105 human, 16 rat, and 16 mouse donors. This includes a temporal map of mouse myocardial infarction (MI) and the evaluation of a novel therapeutic approach targeting polyploid CMs in a rat model. Selection of samples was performed based on pathologist assessment and refined by UNI, a foundation model for pathology image analysis^28^. Samples were measured using probe-based single-nucleus RNA sequencing (snRNA, Chromium Fixed RNA Profiling) and spatial transcriptomics (MERFISH, Xenium) techniques using fixed-frozen or formalin-fixed paraffin-embedded (FFPE) cardiac tissue. An advanced computational framework exploring histology and sample level spatial information identified several novel disease-associated cell states and profibrotic tissue niches in ischemic and non-ischemic heart disease. To take full advantage of the subcellular resolution of our spatial transcriptomic measurements, we studied patterns in the position of individual transcripts across CMs. We identify multiple cell state-specific subcellular patterns, among others a specific signature enriched in the intercalated disk (ICD) region, the cardiomyocyte-cardiomyocyte interface. In summary, our study provides a high-resolution, cross-species dataset that distinguishes polyploid from diploid CM states, resolves their interactions with other cardiac cell types, and delineates the regulatory networks underlying myocardial remodeling in ischemic and non-ischemic failing human myocardium. Specifically, we identify polyploid CMs as key regulators of maladaptive cardiac remodelling and TNIK (TRAF2 and NCK interacting kinase), selectively expressed in polyploid CMs, as a novel therapeutic target limiting cardiac maladaptive remodelling.

### Spatial multi-omics analysis of human heart disease

To characterize cellular heterogeneity and tissue remodelling in ischemic and non-ischemic human heart disease, we analysed heart tissue of 105 adult and fetal human donors and 16 rat and 16 mouse samples (**Fig. 1a-c**), using probe-based snRNA-seq (Flex) and high-resolution spatial transcriptomics (ST; MERFISH/Xenium) (**Fig. 1d**, **Extended Data Fig. 1a-b; Supplementary Table 1**). In total, the snRNA-seq dataset includes 997,362 nuclei and the ST dataset 3,593,941 nuclei (**Fig. 1d**). Human samples were obtained from left ventricular and septal walls, encompassing controls from non-failing donor hearts and patients with ischemic cardiomyopathy (ICM), non-ischemic dilated cardiomyopathy (DCM) of idiopathic etiology, and acute myocardial infarction (ICM_AMI) to capture a wide range of tissue remodelling dynamics (**Supplementary Table 1**). The tissue specimens were collected at heart transplantation or during implantation of a ventricular assist device. The five fetal heart tissue samples were obtained from human fetuses from terminated pregnancies between 12 and 21 post-conception weeks (p.c.w.) (**Fig. 1a**) from the Dutch Fetal Biobank. In addition, we analyzed heart tissues following drug treatment during the cardiac tissue remodelling phase (**Fig. 1b)** in rats (total n=52, n=16 for snRNA-seq) and following myocardial infarction (MI) at five different time points (**Fig. 1c, Extended Data Fig. 1a; Supplementary Table 1**) in mice (n=16, snRNA-seq + Xenium). For each cardiac sample, we obtained a 25 µm tissue section from either fixed-frozen or FFPE tissue specimens for nuclei isolation. We optimized our probe-based snRNA-seq isolation protocol to achieve high data quality and the lowest possible background from ambient RNA. 5 µm tissue sections were collected for high-resolution spatial transcriptomics (**Fig. 1d**, **Methods**). From each sample, an H&E staining was performed and evaluated by an expert cardiac pathologist.

**Figure 1.**
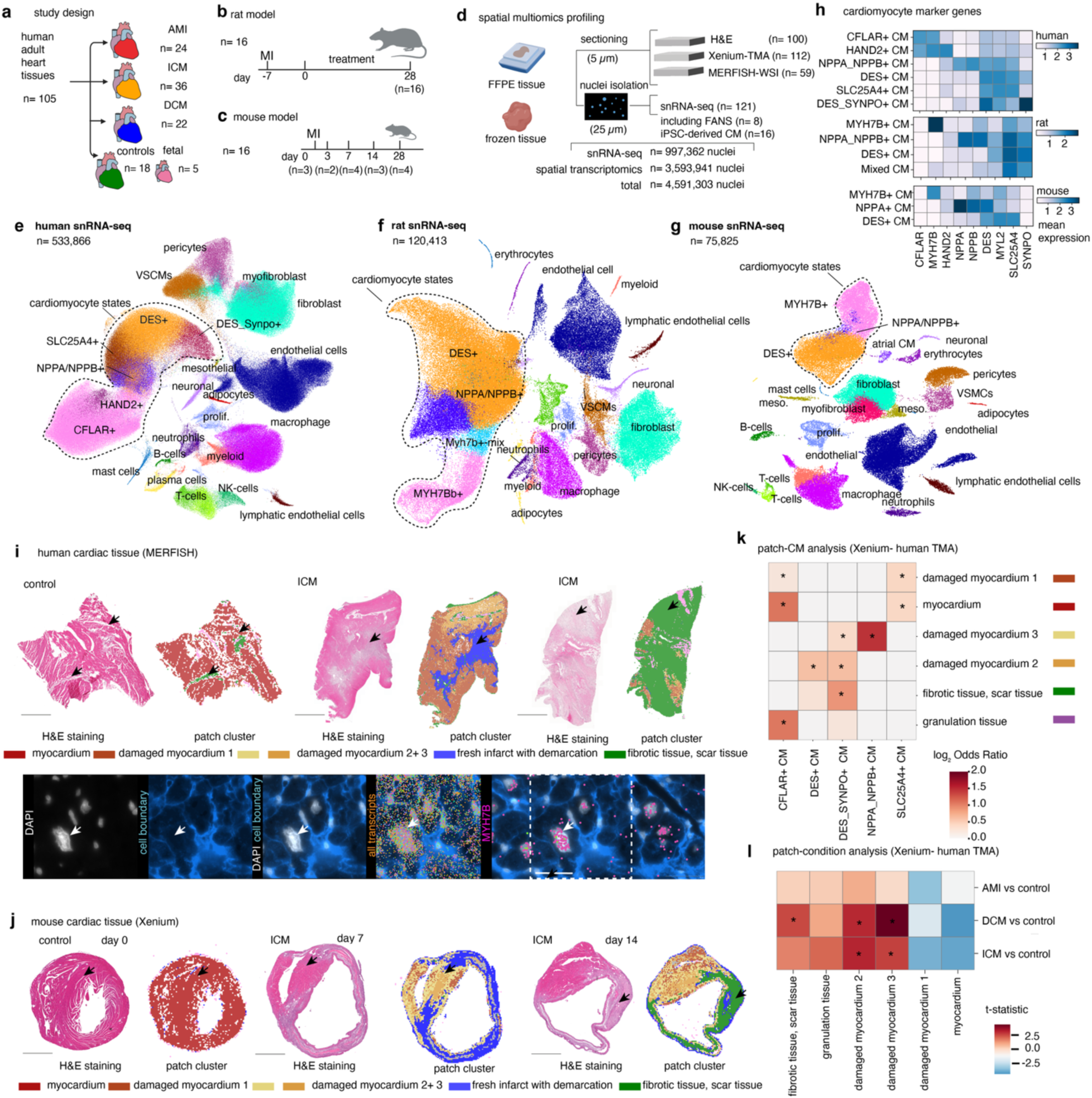
Cross-species single-nucleus and spatial multimodal atlas identifies conserved and human-specific cardiomyocyte states in heart disease. **a,** Study design and human cardiac tissue cohort. Adult human heart samples (n = 100) were obtained from non-failing controls, ischemic cardiomyopathy (ICM), dilated cardiomyopathy (DCM), and acute myocardial infarction (AMI), complemented by fetal heart samples (n = 5). **b,** Rat MI model. Hearts were harvested at day 7 post-MI and after treatment at day 28. **c,** Mouse myocardial infarction (MI) model. Hearts were collected at baseline and at multiple time points following MI (days 1, 3, 7, 14, and 28). **d,** Overview of spatial multimodal profiling. FFPE and frozen cardiac tissues were processed for histology (H&E), spatial transcriptomics (Xenium and MERFISH), and single-nucleus RNA sequencing (snRNA-seq). In total, 4,591,303 nuclei were profiled across snRNA-seq and spatial transcriptomics. **e,** UMAP embedding of human snRNA-seq data (n = 533,866 nuclei), colored by major cardiac cell types. Cardiomyocyte states are highlighted, including DES⁺, NPPA/NPPB⁺, SLC25A4⁺, DES_SYNPO⁺, HAND2⁺, and CFLAR⁺ cardiomyocytes. **f,** UMAP embedding of rat snRNA-seq data (n = 120,413 nuclei), revealing conserved cardiomyocyte states, including DES⁺, NPPA/NPPB⁺, MYH7B⁺, and mixed cardiomyocyte populations. **g,** UMAP embedding of mouse snRNA-seq data (n = 75,825 nuclei), identifying DES⁺, NPPA/NPPB⁺, and MYH7B⁺ cardiomyocyte states alongside non-myocyte populations. **h,** Heatmaps showing expression of selected cardiomyocyte marker genes across cardiomyocyte states in human (top), rat (middle), and mouse (bottom), highlighting conserved and species-specific transcriptional programs. Expression in logarithmic counts. **i,** Spatial mapping of cardiomyocyte patch clusters in human cardiac tissue using MERFISH. Representative H&E-stained sections and corresponding patch cluster annotations are shown for control and ICM samples. Bottom panels show high-resolution MERFISH images with DAPI and selected marker genes (MYH7B). Scale bar indicates 1 mm. **j,** Spatial mapping of myocardial patch clusters in mouse hearts using Xenium across control, MI day 0, day 7, and day 14. H&E staining, patch cluster annotations, and spatial transcriptomic overlays are shown. Marker expression highlights cardiomyocyte and fibroblast-rich regions. Scale bar indicates 1 mm. **k,** Patch-level cardiomyocyte (path-CM) composition analysis in human Xenium tissue microarrays (TMAs), showing enrichment of cardiomyocyte states across myocardial, damaged myocardium, fibrotic/scar, and granulation tissue regions. Asterisks indicate statistically significant enrichments. Significance was determined by Fisher’s exact test with adjusted p-values < 0.05. **l,** Patch-condition analysis comparing AMI, DCM, and ICM to controls in human Xenium TMAs, revealing condition-specific enrichment of myocardial damage and fibrotic tissue states. Significance was determined by t-test with adjusted p-values < 0.05. Asterisks denote statistically significant differences.

For snRNA dataset annotation we followed a two-step approach. An initial graph-based clustering^29^ and annotation based on marker genes resulted in the annotation of 20 major cell types across the three species (**Supplementary Table 2**). Each major cell type was then subclustered into detailed dataset-specific cell states with a focus on cross-species cardiomyocyte diversity (**Fig. 1e-g**, **Supplementary Fig. 1-3)**. Our snRNA-seq data contained a median of 2,358 UMIs and 1,403 detected genes per nucleus for the adult human, 1,190 UMIs and 612 detected genes per nucleus for rat, and 1,512 UMIs and 1,051 detected genes per nucleus for mouse (**Supplementary Table 2**).

Altogether, our cross-species snRNA data identified several novel CM cell states, as well as cell types not captured in previous studies, such as neutrophils (**Fig. 1e-g**). The enhanced recovery originates from a nuclei-isolation protocol optimized for fixed heart tissue that reduces ambient-RNA bias and minimizes loss of nuclei with atypical morphology (e.g. neutrophils)^30,31^. While the transcriptomic complexity (genes detected per UMI) was comparable to our previous polyA-3’-enriched snRNA dataset^11^, our new data achieves substantially higher gene detection rate (up to 14,452 genes vs 6,202 genes) and deeper sequencing coverage (2,534 vs 2,117 median UMIs per cell), indicating improved capture efficiency and transcriptome sampling (**Extended Data Fig. 1c)**.

Marker gene analysis identified two primary CM clusters conserved across three mammalian species, each containing distinct subclusters (**Fig. 1e-h**). The first major group was characterized by *DES* expression and included two specific CM subpopulations: one defined by *NPPA* and *NPPB* expression, and another human-specific subcluster defined by *DES⁺/SYNPO⁺* expression. These *DES⁺/SYNPO⁺* CMs displayed a transcriptional program enriched for cytoskeletal remodeling genes—including *SYNPO*, *SYNPO2L*, and *SYNPO2*—alongside increased *TAZ* (*WWTR1*) expression, consistent with a mechano-adaptive, disease-associated state (**Extended Data Fig. 1d**). The second major CM group was defined by the expression of *CFLAR* and *MYH7B* (or *Myh7b* in rodents). *MYH7B* represents an ancient member of the myosin heavy chain motor protein family, and its expression has not previously been described in a unique adult human CM subpopulation^32^. Unlike the alpha and beta myosin heavy chains (encoded by *MYH6* and *MYH7*), *MYH7B* does not produce a full-length protein due to post-transcriptional exon-skipping^33^; instead, it functions as a long non-coding RNA that regulates the ratio of alpha to beta myosin heavy chains and modulates the expression of other sarcomeric genes^32^. Within this second group, we also identified a small, human-specific subfraction of CMs expressing *HAND2*, a cardiac-enriched transcription factor essential for lineage specification during development whose reactivation in adult cells is linked to stress- and remodeling-associated gene programs. *CFLAR* (also known as c-FLIP), an apoptotic regulator with a reported role in beneficial cardiac remodeling^34^, serves as an additional key marker for this novel CM group in humans (**Fig. 1h**). Notably, these novel CM states were not detectable in two previously published datasets^11,35^ utilizing polyA-3’ snRNA technology (**Extended Data Fig. 1e-f**), highlighting the technical sensitivity required to resolve these populations.

In human hearts, CMs were approximately evenly distributed between the CFLAR⁺ and DES⁺ groups at the cohort level, however, substantial inter-individual variability was observed. Across patients, the percentage of CFLAR⁺ CMs ranged from 20% to 80%. (**Extended Data Fig. 1g**). The proportion of CFLAR⁺ CM was higher in ICM and DCM than in ICM_AMI (**Extended Data Fig. 1g**). Notably, variation in the CFLAR⁺/DES⁺ distribution was not associated with clinical measures of cardiac function, including left ventricular ejection fraction (Spearman’s ρ = 0.23, p = 0.17, n = 36) or left-ventricular end-diastolic diameter (LVEDD) (Spearman’s ρ = −0.27, p = 0.10, n = 36), in samples with available metadata.

To further resolve disease-associated compositional changes, we analyzed cell abundance shifts in our human snRNA data (**Extended Data Fig. 1h**). ICM_AMI samples exhibited a pronounced increase in immune cell populations, including SPP1⁺ macrophages, monocytes, plasma cells, and mast cells, alongside expansion of myofibroblasts and CTHRC1⁺ fibroblasts (**Extended Data Fig. 1h)**. CM sub-states also showed disease-specific remodeling patterns. CFLAR⁺ CMs were significantly expanded in ICM and DCM, whereas ICM_AMI was characterized by a reduction of multiple CM states, including DES⁺, SLC25A4⁺, and specific CFLAR⁺ CM substates (0, 3, and 4) (**Extended Data Fig. 1h**). Additionally, perivascular and ELN⁺ fibroblasts increased significantly only in ICM and DCM, while in ICM_AMI myofibroblast and CTHCR1⁺ fibroblasts were significantly increased, highlighting divergent stromal remodeling programs between chronic and acute cardiac injuries (**Extended Data Fig. 1h**).

We used consecutive FFPE sections to perform spatial transcriptomic experiments with probe sets covering 500 (MERFISH) or up to 5,100 (Xenium) genes (**Extended Data Fig.1b**). Panel design was based on multiple publicly available datasets (**Methods**). In larger cells like CMs, this allowed us to recover up to 10,000 transcripts per cell, while across all cell-types 175 to 306 median genes per segmented cell/nuclei were detected (**Fig. 1i-j**, **Extended Data Fig. 2a**). Out of 150 biosamples (human, mouse and rat), about 50% were measured in both snRNA and spatial modalities (**Extended Data Fig. 1a**).

Segmentation of CMs is an unsolved challenge, impeded by their elongated and irregular shape. We trained “heartbreaker”, a Cellpose^36^ model finetuned on expert annotated sections of our MERFISH data. This approach demonstrated superior performance relative to other state-of-the-art algorithms (**Extended Data Fig. 2b-c; Methods)**. Due to the lower signal-to-noise ratio of cell membrane staining in Xenium, we opted for nuclei segmentation using CellPose for the processing of those samples (**Extended Data Fig. 2d)**. For our subcellular analysis of one tissue microarray (TMA) with 26 samples, we performed a post-Xenium IF staining with wheatgerm agglutinin (WGA)^37^ and DAPI, which allowed us to confidently segment cells and nuclei (**Extended Data Fig. 2e**).

We annotated each spatial dataset by transferring cell state labels from the corresponding species-matched snRNA-seq dataset. To ensure robust annotations, we used five state-of-the-art label transfer tools^38–42^ and assigned each cell the label predicted by the majority of tools. The agreement between tools served as a measure of annotation confidence, indicating which cell states could be reliably distinguished in the spatial data despite capturing only a fraction of the genes measured in the whole-transcriptome snRNA-seq datasets. When tools disagreed, cell types were merged into broader categories until at least three out of five tools agreed. This resulted in comparable cell type proportions between the spatial and snRNA-seq datasets from the same donor (Spearman’s ρ > 0.8).

The inclusion of histology (e.g., H&E staining) provided an additional modality that enabled an integrative, cross-species analysis of the spatial data (**Fig. 1i+j**). To analyze the H&E stainings, we employed a deep learning model specialized in heart tissue^28^. We divided the images into small patches and used the model to classify them based on their histological features (**Extended Data Fig. 3a**). This method successfully distinguished several specific tissue compartments, effectively bridging the analysis between human and mouse samples (**Extended Data Fig. 3b**). The identified tissue compartments effectively captured the transition from the healthy heart (myocardium, adipose tissue) through stress (damaged myocardium 1, 2 and 3), major damage (fresh infarct with demarcation, granulation tissue, necrotic tissue) and chronic damage (fibrotic tissue, scar tissue) (**Fig. 1i, Extended Data Fig. 3b**).

While the majority of tissue compartments were consistent with the pathologist’s annotation, we observed patch sub-clusters (damaged myocardium 1, 2 and 3), that were not readily identifiable by the experts (**Extended Data Fig. 3c**). To uncover compositional differences we related the cell types identified in our spatial transcriptomic measurement (Xenium measurement of human TMAs) to the H&E-derived annotation. We detected differences in the CM composition of different areas (**Fig. 1k**). While CFLAR⁺ CMs were enriched in myocardium, damaged myocardium 1 and granulation tissue, DES⁺ CMs enriched in damaged myocardium 2 together with DES-SYNPO⁺ CMs, which were also found in fibrotic and scar tissue regions. NPPA-NPPB⁺ CMs (border-zone stressed cardiomyocyte signature) were found in high levels in the patches annotated as myocardium damaged 3 in both human and mouse. We could identify an enrichment of immune cells (macrophages, monocytes) in the fibrotic and granulation tissue (**Extended Data Fig. 3d**). Myofibroblasts were localized especially in damaged myocardium 2 and 3 and fibrotic scar tissue regions. Comparing the different disease groups (ICM_AMI, ICM, DCM) we were able to identify an enrichment of myocardium and damaged myocardium 1 in controls whereas damaged myocardium 3 was enriched in DCM as well as fibrotic/scar tissue (Fig. 1l). Quantitative analysis of nearest-neighbor counts revealed that DES+ CMs form more cohesive spatial clusters (based on nearest-neighbor distances), while CFLAR+ CMs are more spread out (**Extended Data Fig. 3e**), supporting the model of distinct spatial patterning of CM states in the diseased human myocardium.

Together, these analyses establish a spatially resolved, cross-species dataset integrating single-nucleus transcriptomics, spatial gene expression, and histology-based tissue annotation. We identify novel CM (CFLAR⁺, DES⁺, DES-SYNPO⁺, SLC25A4⁺) beyond the established stressed NPPA-NPPB⁺ CM state. By anchoring transcriptomic states to specific morphological zones—ranging from healthy myocardium to chronic scar—we demonstrate that novel CM states (CFLAR⁺, DES⁺, etc.) are not transient signatures but are related to specific stages of pathological remodeling.

### Spatial niches of human heart disease

While the histology-based compartments provided a macro-scale framework of tissue injury, they represent broad morphological zones rather than precise functional units. Cellular colocalization serves as a proxy for the complex communication networks that operate at different length scales to maintain homeostasis or drive pathology. To bridge the gap between these large-scale regions and cellular behavior, we moved from tissue compartments to spatial niches—repeating tissue structures with co-localized cells—and characterized how they differed across patients and disease conditions (ICM_AMI, ICM, or DCM).

We performed a sample level cell–cell colocalization analysis using a framework combining NicheSphere and PILOT^43^ (**Extended Data Fig. 4a, Methods**). Using NicheSphere, we identified six distinct niches in the cell interaction networks in the contrast control vs. ICM samples (**Fig. 2a**). Similar to the H&E spatial data analysis, our spatial transcriptomic analysis revealed that CM states were localized in separate niches e.g. CFLAR⁺ and SLC25A4⁺ CM together with macrophages, pericytes and lymphatic endothelial cells in niche 4. Cells comprising this niche, characteristic of healthy myocardium, showed diminished spatial colocalization in ICM. Regarding niches with increased colocalization in ICM, niche 3 includes CM cell states (DES⁺, NPPA_NPPB⁺ and DES_Synpo⁺). Niche 1 includes myofibroblasts clustered together with proinflammatory macrophages, monocytes and fibroblasts forming a unique profibrotic and proinflammatory tissue niche (**Fig. 2a**). Focusing on colocalization statistics associated with the different novel CM subtypes, only myofibroblasts increase their colocalization with all CM subtypes (**Fig. 2b**). However we also noted distinct differences between CFLAR⁺ and SLC25A4⁺ CMs compared to the other CMs. These CM states lost their colocalization to macrophages, pericytes and mast cells, whereas DES_SYNPO⁺ CMs uniquely increased their localization to tissue resident macrophages (**Fig. 2b**). We next compared the spatial niches (based on spatial transcriptomics) with our identified patch clusters (based on H&E staining) and observed strong concordance between the two approaches. (**Extended Data Fig. 4b**). The niche 4 (CFLAR+, SLC25A4+ CMs) was related to healthy myocardium, whereas niche 3 (DES, NPPA-NPPB-CMs) was significantly enriched in damaged myocardium 1-3 (**Extended Data Fig. 4b**). By resolving the cellular neighborhoods of each CM state, we demonstrate that cardiac remodeling involves the formation and dissolution of specific multicellular niches: loss of homeostatic CFLAR⁺/SLC25A4⁺-macrophage-pericyte networks in ICM parallels emergence of profibrotic DES⁺/myofibroblast assemblies, revealing niche-level reorganization as a hallmark of disease progression.

**Figure 2.**
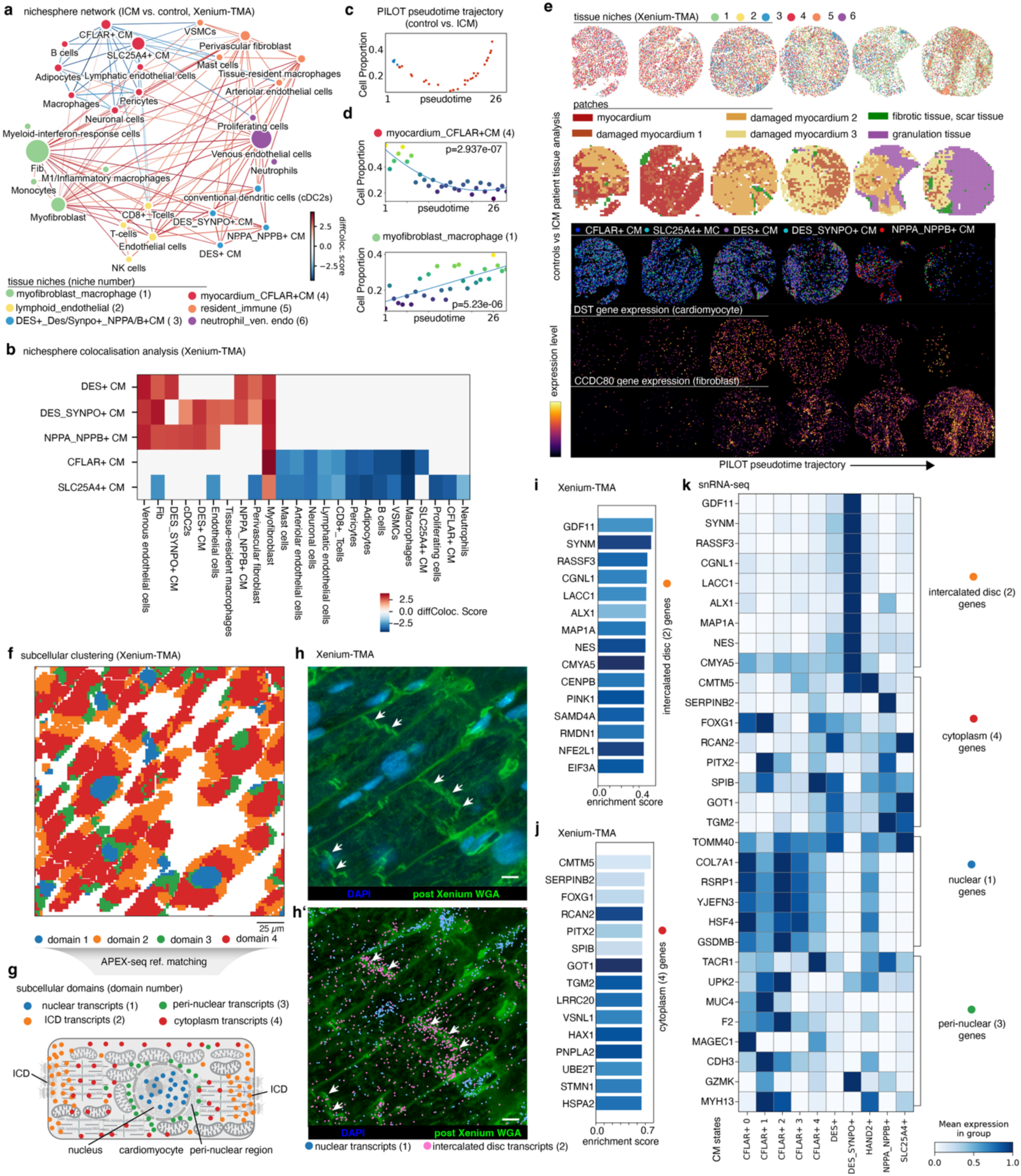
Spatial tissue niches and subcellular transcript organization reveal cardiomyocyte state–specific remodeling in cardiomyopathy. **a,** NicheSphere interaction network control vs. ICM. Network representation of spatial tissue niches inferred from Xenium-TMA sections comparing ischemic cardiomyopathy (ICM) with controls. Nodes represent cell-type–specific niches and edges indicate significantly altered spatial interactions, with red and blue edges denoting increased or decreased interactions in ICM, respectively. **b,** Spatial colocalization analysis. Heatmap of differential colocalization (diffColoc score) between cardiomyocyte (CM) states and other cardiac cell types across Xenium-TMA samples. For **a** and **b**, only statistically significant interactions are shown (adjusted Wilcoxon’s rank sums test; p-value < 0.05). **c,** PILOT pseudotime trajectory (control as blue dots vs. ICM as red dots). Trajectory of TMA samples illustrating progressive niche remodeling from control to ICM tissue states. **d,** Niche proportion changes along pseudotime. Changes in relative abundance (y-axis) of myocardium-associated CFLAR⁺ CMs (top) and myofibroblast–macrophage niches (bottom) along pseudotime (x-axis), indicating reciprocal dynamics during disease progression (p-value - goodness of fit test on the regression of the cell proportion and pseudo-time). **e,** Tissue niche composition across six selected Xenium-TMA samples along the Trajectory from early to middle and late stages. Spatial maps showing niche assignments (top), histopathology-based tissue annotation (middle; myocardium, damaged myocardium, fibrotic and granulation tissue), and CM state distributions (bottom) across selected control and ICM samples. Representative gene expression patterns of genes with trajectory expressific changes are shown: DST (cardiomyocytes) and CCDC80 (fibroblasts). **f,** Subcellular clustering. Spatial clustering of transcripts within cardiomyocytes identifies four recurrent subcellular domains across Xenium-TMA sections. Scale bar, 25 µm. **g,** Reference-based domain annotation. Mapping of Xenium-derived subcellular domains to APEX-seq reference signatures, assigning nuclear, intercalated disc (ICD), peri-nuclear, and cytoplasmic transcript compartments. **h,** Imaging validation of subcellular domains. Post-Xenium wheat germ agglutinin (WGA) staining and DAPI highlighting cardiomyocyte structure and nuclei, with arrows indicating enrichment of domain-specific transcripts. Bottom panel shows spatial localization of nuclear and ICD-associated transcripts. **i,** ICD-associated gene enrichment. Bar plot of genes enriched in intercalated disc–associated transcripts identified in Xenium-TMA data. **j,** Cytoplasmic gene enrichment. Bar plot of genes enriched in cytoplasmic transcript domains in Xenium-TMA data. **k,** snRNA-seq validation of subcellular gene programs. Heatmap showing mean expression of domain-specific genes across CM states in snRNA-seq data, confirming nuclear, peri-nuclear, cytoplasmic, and ICD-associated transcriptional programs. Particular enrichment is found of ICD domain 2 associated transcripts in DES_SYNPO+ CMs.

To study the dynamical changes of spatial niches, we performed a sample level trajectory analysis using PILOT (**Fig. 2c**). A statistical analysis on niche changes over the trajectory indicated a decline of niche 4 (CFLAR myocardium) and expansion of niches 1, 2 and 5 (macrophage, myofibroblast, lymphoid cell, and resident immune niches), as well as a transient increase of niche 3 with DES-CMs (**Fig. 2d; Extended Data Fig. 4c**). Selected cores display changes in niche, patches, cells and genes among the disease trajectory (**Fig. 2e)**. For example, CCDC80 was identified as a gene with a significant increase in expression in PILOT’s trajectory for fibroblasts and myofibroblast cells (**Fig. 2e**). We further found this gene was highly expressed in fibroblasts (not shown) particularly in the CTHRC1⁺ fibroblast subpopulation in the snRNA-seq data **(Extended Data Fig. 4d)**. Other trajectory-related genes for CFLAR-CMs include CM contractile genes (*TNNT2*, *MYBPC3*) showing a reduction, whereas *TTN*, *PDE4DIP* and *DST* have an increased expression during disease progression (**Fig. 2e; Extended Data Fig. 4e**). Thus, PILOT trajectory analysis suggests that CFLAR⁺ CMs in ICM undergo a transition from a sarcomere-rich contractile state toward a stress-adaptive, cytoskeletal, and ECM-anchored (extracellular matrix) phenotype (based on the DST and PDE4DIP gene changes)^44,45^ which is accompanied by increasing activation and presence of ECM-secreting fibroblast subsets.

Comparing controls vs. DCM we identified five cardiac tissue niches (**Extended Data Fig. 4f-h**). Two niches had similar cell types and changes to ICM niches, i.e. decrease in colocalization of cells in the CM-rich CFLAR myocardium (niche 4) and increase of colocalization of cells in myofibroblast and macrophages (niche 2) (**Extended Data Fig. 4f**). However, our analysis revealed that the DES_Synpo⁺ CM state constituted their own cardiac niche in DCM with endothelial and dendritic cells (niche 3) (**Extended Data Fig. 4f-g**). Similar to ICM, all CMs increased their spatial colocalization with myofibroblasts (**Extended Data Fig. 4g**). Interestingly, the DES_Synpo⁺ CMs demonstrated more spatial interactions with other cell-types compared to ICM, indicating a potential unique role in DCM. Moreover, in DCM samples, niche 3 with DES-SYNPO⁺ CM was highly enriched in fibrotic tissue patches (**Extended Data Fig. 4i**). A comparison of niche distribution in ICM vs. DCM samples (**Fig. 2e** vs. **Extended Data Fig. 4j**) revealed distinct spatial architectures. ICM samples exhibited significantly higher spatial clustering compared to the diffuse, fine-grained distribution observed in DCM (Wilcoxon rank-sum statistic = 4.09, p < 0.0001) (**Extended Data Fig. 4k**). This focal organization was especially pronounced for inflammatory ICM niches 5 and 6, which formed contiguous domains.

In CFLAR⁺ CMs of contrast control vs. DCM samples, contractile genes (*TNNT2*, *MYBPC3*) decline early as in ICM (**Extended Data Fig. 4l**), while *TTN*, *OBSCN* and the NMD kinase *SMG1* increase, indicating a CM-intrinsic program of sarcomere scaffold realignment. Comparative niche analysis (ICM vs. DCM) demonstrates conserved localization of CFLAR⁺ and NPPA_NPPB⁺ CMs in distinct cardiac tissue niches across cardiomyopathy etiologies, contrasted by disease-specific redistribution of DES-SYNPO⁺ CM states (**Extended Data Fig. 5a**). For the controls vs. ICM_AMI contrast, we detected six tissue niches using NicheSphere (**Extended Data Fig. 5b**). Niche 1 contained all CM substates and represented the niche losing colocalization interactions across other cell types (**Extended Data Fig. 5b**). Interestingly, an increased colocalization is observed between myofibroblasts and CFLAR⁺ CM cells as in other contrasts (**Extended Data Fig. 5b**). Among the niches with increased colocalization in ICM_AMI condition, we also observe a fibrotic niche (niche 4) and a tissue resident (niche 6) with increase in ICM_AMI (**Extended Data Fig. 5d**). The latter niche included SPP1⁺ tissue-resident macrophages, which represent the cell type with the highest amount of ICM_AMI specific interactions, recapitulating our previous findings^11^.

### Subcellular features of human cardiomyocytes

Next we analyzed the subcellular localization patterns of RNA molecules across individual human CMs using BENTO^46^, a toolkit for the analysis of subcellular RNA distribution. Unsupervised clustering revealed four distinct subcellular domains characterized by recurring micrometer-scale expression patterns **(Fig. 2f-j).** Enrichment analysis of these domains based on known subcellular organelle mRNA signatures from APEXseq^47^ confirmed a nuclear (domain 1), perinuclear (domain 3), and two cytoplasmic domains (domain 2, 4) **(Fig. 2f-h, Extended Data Fig. 5e).** The two cytoplasmic domains were enriched for different mRNAs (**Fig. 2i-j, Extended Data Fig. 5e**). We hypothesized that the cytoplasmic domain 2 could correspond to CM-CM cell contacts, the area where the ICD supports the synchronized contraction of the heart also based on previous reports^48^. This was confirmed by co-staining with WGA, where the ICD is distinguishable by a higher signal intensity compared to lateral membranes. (**Fig. 2h-h’, Extended Data Fig. 5e-h**). The ICD domain was enriched with genes related to membrane-cytoskeleton linkage like *CMYA5*, an important regulator of the cardiac dyad architecture^49^ (**Extended Data Fig. 5h**). Notably, the RNA-level presence of the RNA-binding protein *SAMD4A*^50^ and of the translation initiation factor *EIF3A*^51^ is consistent with localized post-transcriptional regulation and translational activity, raising the possibility of spatially organized protein synthesis in proximity to the ICD **(Fig. 2i)**. After identifying genes with specific subcellular expression patterns, we explored their expression levels in our human snRNA data (**Fig. 2k**). Our analysis shows an enrichment of ICD-related (domain 2) transcripts in DES_SYNPO⁺ CMs and of nuclear and perinuclear genes (domain 1 and 3) in CFLAR⁺ CM clusters **(Fig. 2k)**. In summary, we define distinct subcellular mRNA domains in human CMs and demonstrate that their utilization differs systematically across our novel CM cell states, with ICD-associated transcripts preferentially enriched in DES_SYNPO⁺ CMs and nuclear transcripts in CFLAR⁺ CMs, consistent with divergent adaptive subcellular RNA localization programs in human heart disease.

### Novel human cardiomyocyte cell states

Next, we analyzed the molecular differences of the CM cell-states (**Fig. 3a-b**). We based our annotation of the CMs clusters on unique marker gene expression (**Fig. 3b**). Desmin (*DES* gene), as intermediate filament in CMs, has important structural roles; it connects several CMs specific subcellular structures like the ICD complexes and with organelles like mitochondria and the nucleus^52,53^. HAND2⁺ CMs represented the least represented CM cell state with 1.77% (**Supplementary table 2**). The re-expression of *HAND2*, a fetal CM transcription factor^54^, combined with *MYH7B* points towards a fetal gene expression shift. *CFLAR*, or *c-FLIP*, a direct Casp8 inhibitor and master anti-apoptotic regulator^55^, is uniquely and highly expressed in human CMs when compared to mouse and rat snRNA-seq (**Fig. 3b**, **Fig. 1h**). It has been demonstrated that the gene *CFLAR* protects CMs from cell death in MI^56^. Based on the apparent specificity of CFLAR⁺ marker genes to enlarged nuclei in our spatial transcriptomics data we hypothesized that the novel HAND2⁺ and CFLAR⁺ CMs states could be related to polyploid, or genome duplicated CM cell states. Nuclear DAPI stainings have been used to identify polyploid nuclei in tissues previously^57^. To confirm our hypothesis we extracted morphometric properties (using CellProfiler^58^) of CM nuclei from our spatial transcriptomics Xenium dataset and checked for association with our transcriptomically defined CM cell state identities. This analysis showed a large degree of heterogeneity in nuclear appearance for the different CM states (**Fig. 3c-d, Extended Data Fig. 6a-d**), and especially between CFLAR⁺ and DES⁺ CMs. Differences in DAPI texture and nuclei size were also obviously distinct upon visual inspection (**Fig. 3d, Extended Data Fig. 6a**). Among the most distinct nuclear features in CFLAR⁺ CMs were the area of the nuclei followed by features related to DAPI intensity, edge intensity, entropy and granularity, consistent with differences in polyploidy between the analyzed CM states and potentially linked to chromatin changes **(Extended Data Fig. 6b-c).** A machine learning model trained on these nuclei images was able to differentiate CFLAR⁺ and DES⁺ CM based on both CellProfiler morphometrics as well as vision model (Dinov3) features (**Extended Data Fig. 6d, Methods**).

**Figure 3.**
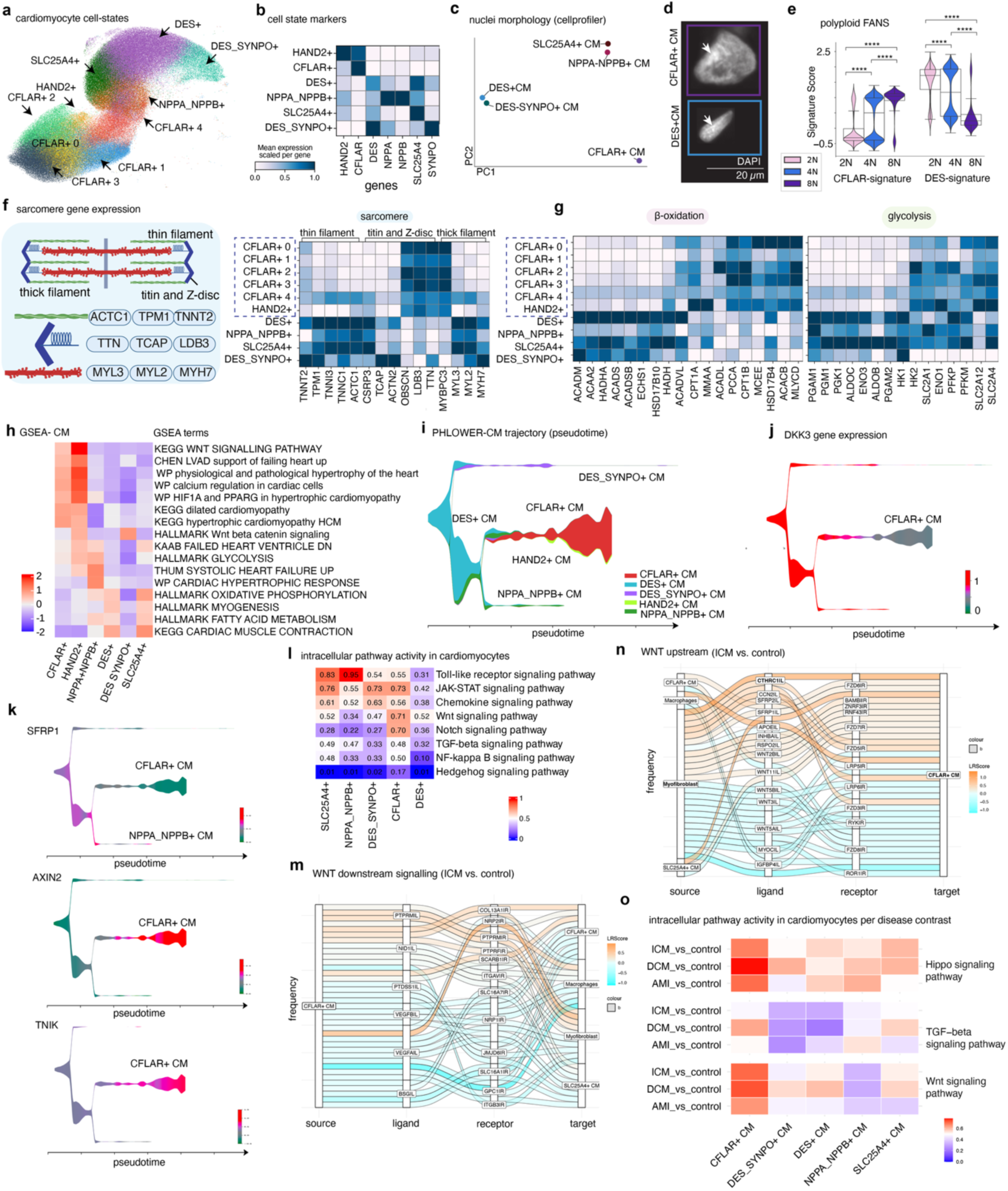
Molecular, structural, metabolic, and signaling features of novel human cardiomyocyte cell states. **a,** UMAP embedding of snRNA-seq highlighting distinct CM populations, including CFLAR⁺, DES⁺, DES_SYNPO⁺, HAND2⁺, NPPA_NPPB⁺, and SLC25A4⁺ CMs. Arrows indicate cluster identities. **b,** Cell state marker genes. Heatmap showing scaled mean expression of representative marker genes across CM states, confirming transcriptional distinctions between CMs. **c,** Nuclei morphology analysis. PCA of CellProfiler-derived nuclear morphology features from DAPI staining separates CM states, indicating systematic differences in nuclear shape and size across transcriptional identities. **d,** Representative nuclei images. Immunofluorescence images of DES⁺ and CFLAR⁺ cardiomyocyte nuclei illustrating distinct nuclear morphologies (arrows). Scale bar, 20 µm. **e,** Polyploidy enrichment using FANS. Violin plots of cell-state signature scores across nuclei with different ploidy levels (2N, 4N, 8N) obtained by FANS. CFLAR and DES signature enrichment differs by ploidy state (p-value < 0.0001). **f,** Sarcomere gene expression. Schematic of sarcomeric organization (thin filaments, thick filaments, titin and Z-disc) and heatmap of corresponding gene expression across CM states, highlighting differential regulation of structural components. **g,** Metabolic gene programs. Heatmaps showing expression of genes involved in β-oxidation (left) and glycolysis (right) across CM states, indicating metabolic specialization. **h,** Gene set enrichment analysis (GSEA) of hallmark, KEGG, and WikiPathways terms across CM states, revealing differences in hypertrophic signaling, metabolism, WNT signaling, and cardiac contraction programs. **i,** Differentiation trajectory of CM states. PHLOWER-based trajectory inference depicting transitions between CM states along pseudotime, with relative contributions of CFLAR⁺, DES⁺, DES_SYNPO⁺, HAND2⁺, and NPPA_NPPB⁺ CMs. **j,** DKK3 expression dynamics. Smoothed expression of DKK3 along the CM pseudotime trajectory, highlighting state-specific regulation. **k,** WNT pathway–related genes along cardiomyocyte trajectories. Expression dynamics of SFRP1, AXIN2, and TNIK across CM states along pseudotime, illustrating coordinated regulation of WNT signaling components. **l,** Intracellular pathway activity (ICM vs. control). Heatmap summarizing inferred intracellular activity scores of key signaling pathways (e.g., Toll-like receptor, JAK–STAT, WNT, TGF-β, NF-κB, Hedgehog) across CM states, highlighting differential transcriptional response between ICM versus control hearts. **m,** WNT upstream signaling (ICM vs. control). Sankey plot display ligand-receptor interactions upstream of WNT signalling in CFLAR+ CM cells, comparing ICM with control hearts. Positive values indicate increased signaling strength in ICM, whereas negative values indicate reduced interaction strength. **n,** Cell-cell communication downstream of WNT signaling (ICM vs. control). Sankey plot displaying ligand–receptor interactions downstream of WNT pathway in CFLAR+ CM cells comparing ischemic cardiomyopathy (ICM) compared with controls; ligands correspond to transcriptional targets of WNT signaling. **o,** Disease-specific pathway activity. Heatmap of inferred intracellular pathway activity across CM states comparing ischemic cardiomyopathy (ICM), dilated cardiomyopathy (DCM), and acute myocardial infarction (AMI) with controls, focusing on Hippo, TGF-β, and WNT signaling pathways.

To validate this result we next performed FANS (fluorescent activated nuclei sorting) based on DAPI staining and resorted the nuclei to confirm the successful enrichment of diploid (2N), tetraploid (4N) and octaploid (8N) nuclei (**Fig. 3e, Extended Data Fig. 6e-i**). We performed this experiment on the tissue specimen of four patients where enough material was available and sorted for 2N, 4N and 8N DAPI content, respectively. This yielded eight enriched snRNAseq datasets (see schematic in **Extended Data Fig. 6e**). A median UMI count per nuclei of 3,233 and median of 1,640 genes per nuclei indicated that the FANS data had similar quality as our non-FANS isolation approach (**Supplementary Table 1**). Non-CM cells were mainly found in the diploid (2N) nuclei fraction, while 95% of nuclei in the polyploid fraction were derived from CM (**Extended Data Fig. 6f**). Clustering of all CMs across fractions revealed two distinct CM populations (**Extended Data Fig. 6g**). To connect this data to our main non-FANS snRNA dataset we performed enrichment analysis. The 8N fraction was strongly enriched for the CFLAR⁺ CM signature, while the 2N fraction scored high for the DES⁺ signature (**Fig. 3e, Extended Data Fig. 6i**). Nuclei in the 4N fraction were split into high DES+ and high CFLAR cells, which may represent a transitional cell state (**Fig. 3e, Extended Data Fig. 6h-i**). Taken together the DAPI and FANS experiments confirmed that the CFLAR⁺ CM gene signature is linked to the ploidy of the CMs, while the DES⁺ CM gene signature was more indicative of diploid CM.

While we could not identify the novel CM signatures in two previously published 3’ based snRNA datasets (**Extended Data Fig. 1e+f**), we searched for scRNA-seq or snRNA-seq studies of human heart tissue identifying similar heterogeneous CM states, independent of atrial vs. ventricular phenotypes. The only exception being a dataset from Wehrens et al.^59^, where such CM clusters are described derived from human HCM patients after septal myectomy and whole CM sequencing using SORT-seq (**Extended Data Fig. 6j**). When we mapped the CFLAR^+^ CM and DES^+^ CM gene scores in the Wehrens scRNA-seq dataset, we could identify a clear enrichment and separation of the two scores, fitting to our observation of novel CM states in ICM and DCM patient samples (**Extended Data Fig. 6k**). To increase our confidence in this mapping we employed a dual approach. Using ClusterFoldSimilarity^60^ we calculated similarity scores between the two datasets, based on differential genes expressed for each cluster. In addition, we performed a hyper geometric test (500 genes per cluster), which confirmed an overlap of expression patterns between Wehrens cluster 5 and our polyploid HAND2⁺ and CFLAR⁺ CMs (**Extended Data Fig. 6l**).

We next mapped structural, proliferative, metabolic and chromatin programs across CM states (**Fig. 3f+g, Extended Data Fig. 7a-j**). Contractile sarcomere genes diverged in a CM state-resolved manner: thin-filament and titin/Z-disc gene sets separated clearly across subtypes, indicating remodeling of the sarcomere. In particular *OBSCN*, *LDB3* and *TTN* were upregulated in HAND2⁺ and CFLAR⁺ CMs (**Fig. 3f**). Metabolic pathway analysis revealed a coordinated metabolic re-wiring (**Fig. 3g, Extended Data Fig. 7a-d**). Diploid DES⁺ CMs demonstrated upregulated β-oxidation genes, whereas polyploid HAND2⁺/CFLAR⁺ states showed increase in glycolytic and glutaminolysis gene expression (**Fig. 3g, Extended Data Fig. 7a-c**). Over representation analysis comparing DES⁺ and CFLAR⁺ CMs revealed significant enrichment of mitochondrial, oxidative phosphorylation, and aerobic respiration–related gene sets in DES⁺ CM (**Extended Data Fig. 7d**).

We next compared the expression of cell-cycle regulatory genes (**Extended Data Fig. 7e-f**). In adult CMs three reactivation modes emerged. HAND2⁺ CMs exhibited G1/S-associated gene expression (*MYC*, *CDK4*, *CDK6*, *CCNE1*, *CCNE2*), with relative enrichment for S-phase signatures compared to other states. Polyploid CFLAR⁺ CMs re-expressed late G2/M genes (*CDK1*, *CCNB1*, *AURKA*, *PRC1*, *HMMR*) in the absence of G1/S induction, a pattern compatible with endoreplication-like genome duplication without productive cytokinesis (**Extended Data Fig. 7e-f**). DES_SYNPO⁺ CMs also upregulated cell-cycle associated genes, including *CCND1*/*CCND2* (G1), *MKI67* (pan-proliferation), *TOP2A* (S/G2), and *CDK1* (mitotic entry)-consistent with partial engagement of cell-cycle transcriptional programs (**Extended Data Fig. 7e-f**).

Ion-channel programs segregated sharply by CM state (**Extended Data Fig. 7g**). Polyploid CFLAR⁺ subsets showed the strongest pacemaker/diastolic-depolarization signature (*HCN1*) together with an L-type Ca²⁺ module (*CACNA1C* with *CACNB2*) and broad TRP upregulation (*TRPC1, TRPC3, TRPM4, TRPM7*) (**Extended Data Fig. 7g**). In contrast, diploid subtypes (NPPA_NPPB⁺, SLC25A4⁺, DES⁺) preferentially expressed the noncanonical connexin GJA3 (C*X*46), whereas polyploid CM states expressed GJC1 (CX45), indicating a potential junctional coupling shift at the ICD. Together, these patterns point to a polyploid-biased increase in depolarizing drive (*HCN1*), Ca²⁺ influx capacity (*CACNA1C*/*CACNB2*), mechanosensitive entry (*TRPM4*/*7*; *TRPC1/3*), and an ICD connexin switch (*GJA3*→*GJC1*) (**Extended Data Fig. 7g**). Chromatin and nuclear stability gene programs were likewise CM state-specific (**Extended Data Fig. 7h**). Polyploid CFLAR⁺ CMs, particular the HAND2⁺ state, were enriched for histone acetylation/methylation genes, chromatin-remodeling factors, and nuclear-stability components, consistent with broad epigenomic reorganization accompanying metabolic shift (**Extended Data Fig. 7h)**. This was further supported by the unique upregulation of nuclear receptor gene expression involved in metabolism in CMs, like *PPARα and PPARγ* (**Extended Data Fig. 7i).**

Gene set analysis revealed a clear enrichment of cardiac disease terms in polyploid CMs (CFLAR⁺ and HAND2⁺) including hypertrophic cardiomyopathy (HCM) and dilatative cardiomyopathy terms (DCM) as well as the associated maladaptive cell signalling pathways in cardiac remodelling including WNT signalling pathway (**Fig. 3h, Extended Data Fig. 7j**). NPPA-NPPB⁺ CMs demonstrated enrichment in systolic heart failure gene signature and cardiac hypertrophic response. DES⁺ diploid CMs enriched in metabolic terms oxidative phosphorylation and fatty acid metabolism as well as myogenesis and cardiac muscle contraction (**Fig. 3h, Extended Data Fig. 7d and 7j**).

Based on single-nucleus-resolved RNA expression and open chromatin information from our previous work^11^ we constructed an enhancer based gene regulatory network (eGRN) with scMEGA^61^. The estimated eGRN detected two main regulatory modules **(Supplementary Table 4)**. One module was associated with cells at the beginning of the trajectory (DES⁺ CMs) and one connected to cells at the end of the trajectory (CFLAR⁺, DES-SYNPO⁺ and NPPA-NPPB⁺ CMs). *CTCF*, which is crucial in chromatin organization of the developing heart^62^, was the most well connected transcription factor (TF) (as determined by PageRank centrality) in the GRN module related to DES⁺ CMs (**Supplementary Table 4**). The integrated dataset was then used to construct a cell differentiation tree with PHLOWER^63^. By using DES+ CM cells as root, PHLOWER indicated branches leading to DES-SYNPO⁺ CMs and another branching leading to CFLAR⁺ or NPPA-NPPB⁺ CMs (**Fig. 3i**). We observe lower TF activity of *CTCF* in disease related CM sub-states such as CFLAR^+^, NPPA_NPPB+ and DES_SYNPO^+^ CMs (**Extended Data Fig. 7k**). Along this trajectory, CFLAR⁺ CMs displayed dynamic regulation of WNT pathway modulators, including a decrease expression of the WNT antagonist *DKK3* (**Fig. 3j**) and *SFRP1* (**Fig. 3k**), alongside induction of canonical WNT target genes such as *AXIN2* and the kinase *TNIK* (**Fig. 3k**). The main transcription factor regulated to DES-SYNPO⁺ branch was *WT1* (**Supplementary Table 4**), which is a known epicardial regulator^64^ and is known to directly regulate synaptopodin gene expression e.g. in podocytes in the kidney^65^. Its knock-out is known in zebrafish to lead to a loss of CM expression programs and upregulation of epicardial gene expression^66^. For CFLAR⁺ CMs, predicted regulators include the sonic hedgehog pathway TF *GLI3*, which has a known muscle repair–associated angiogenesis role in myocytes^67^ and *NR2C2* (or *TR4*), a reporter repressor of PPARα nuclear receptor signaling^68^ (**Supplementary Table 4**).

We next analyzed the expression of ligands and receptors of known canonical fetal cell-cell communication and fibrosis related pathways in CMs (**Extended Fig. Data 8a-c**). CFLAR⁺ and HAND2⁺ CMs showed the highest expression among CMs of receptor and ligands associated with maladaptive remodelling including the receptor *NOTCH2* and ligands *JAG2*, *DLL1* together with non-canonical WNT ligands (*WNT5A*, *WNT11*). Among these, HAND2⁺ uniquely presented upregulated *SHH* expression, whereas CFLAR⁺ CM states demonstrated high *PTCH1/SMO* levels. In contrast, DES⁺, NPPA_NPPB⁺, and SLC25A4⁺ CMs preferentially expressed WNT antagonists (*DKK3*, *SFRP1*/*2*) and showed generally attenuated Notch/HH signatures, consistent with a contractile/stress-adapted phenotype that restrains canonical Wnt activity (**Extended Fig. data 8a**). We next analyzed fibrogenic cell-cell communication pathways (TGFβ, PDGF, endothelin, *VEGF*/*ANGPT*, FGF/NRG axes) across CM states (**Extended Fig. data 8b**). DES⁺, NPPA_NPPB⁺, SLC25A4⁺, and especially DES_SYNPO⁺ were enriched for distinct profibrotic and proinflammatory ligands, including *TGFB* isoforms, *PDGFR-A/B*, *VEGFB* and *VEGFC*, *IL6* and *CXCL12* consistent with a CM-derived “injury signal” that can activate fibroblasts and vascular/perivascular cells (**Extended Fig. data 8b-c**). In contrast, CFLAR⁺ and HAND2⁺ CMs preferentially expressed growth-factor receptors (*TGFBR1/2*) together with *ERBB2/ERBB4* and the profibrotic ligands EDN1 and FGF2, indicating a divergent send-receiver profibrotic phenotype between CM states (**Extended Fig. Data 8b-c**).

In order to consider both cell-cell colocalization and changes in ligand-receptor expression across disease conditions, we performed a cell-cell communication analysis with scSeqComm^69^ and CrossTalkeR^70^ by focusing on colocalized cell pairs predicted by NicheSphere (**Fig. 2a; Methods**). We first used CrossTalkeR pagerank to identify highly influential cell types in ICM disease. This indicates the importance (higher communication in ICM) of diploid CM clusters (SLC25A4⁺, DES_SYNPO⁺, DES+⁺) with a highest score for the SLC25A4⁺ CMs and immune cells like neutrophils, macrophages (**Extended Data Fig. 8d**). We next analyzed differences in intracellular signalling in CM from disease samples. scSeqComm calculates differential cellular communication scores, which reflect how cell-cell communication affects down-stream pathways and transcriptional regulation. This indicates an upregulated activity of WNT, NOTCH and Hedgehog signalling in CFLAR⁺ CMs in the ICM vs. control contrast (**Fig. 3l**).

If we consider the signals related to WNT activation towards CFLAR (**Fig. 3m-n**), we observe increased signalling from myofibroblasts including *CTHRC1*^71^*, CCN2 and WNT11* (**Fig. 3m**). In ICM myofibroblasts secreted *CTHRC1*^71^ can bind *FZD6* on CFLAR⁺ CMs to potentiate non-canonical Wnt signaling. These predictions are consistent with our prior spatial analysis showing interaction of CFLAR⁺ CM with myofibroblast in all disease conditions and indicates the activation of WNT signalling in CFLAR⁺ CM cells. Reciprocally, CFLAR⁺ CM show an increased secretion of VEGFA towards myofibroblasts (**Fig. 3n**). Other signals that drive the fibrotic niche created by CFLAR+ CMs are *AREG*, *DLL1* and *FAT4* (**Extended Data Fig. 8e-f**).

Our colocalization analysis (**Fig. 2g**) indicated an unique role of DES-SYNPO⁺ CMs, which formed a DCM specific niche not detected in ICM samples. Moreover, DES-SYNPO⁺ CMs had the strongest proinflammatory gene signatures including IL6, CCL2 and CXCL8 (**Extended Data Fig. 8c**). We identified a reciprocal signalling axis of neutrophils and NK cells to DES-SYNPO⁺ CMs involving *GNAI2* and *CCL5* highlighting a distinct immune-CM axis in DCM (**Extended Data Fig. 8h-i**). When evaluating the intracellular pathway activity across contrasts, we observe an increase in WNT signalling in CFLAR+ CM in all disease conditions in particular in DCM samples (**Fig. 3o**).

Collectively, our results suggest the existence of distinct CM states characterized by polyploidy linked remodeling of contractile, electrophysiological, metabolic, and chromatin programs. These CM state-specific programs appear to converge further on specific trajectories, raising the hypothesis that cardiomyocyte genome multiplications may serve as a central organizing principle of CM-linked remodeling in human heart disease.

### Fetal vs. adult human cardiomyocytes

A central observation in cardiac transcriptomic analysis is that postnatal hearts adopt a fetal-like transcriptional state, also referred to as the “fetal gene hypothesis” in response to stress or chronic injury^72^. Given the expression of cardiac fetal genes like *MYH7B* and *HAND2* in polyploid CMs, we next addressed how conserved this gene expression signature is. To this end, we generated snRNA data using fresh frozen human fetal heart tissues from five timepoints preserved in RNAlater solution (p.c.w. 12 - 22) (**Fig. 4a**). In total we recovered 146,025 nuclei (**Extended Data Fig. 9a**). The nuclei contained a median of 3,531 UMIs and expressed a median of 1,886 genes. Our analyses focused on CMs of the left ventricle and identified six major cell states, which recapitulated the divergent expression profiles of CFLAR⁺ and DES⁺ populations observed in the adult dataset (**Fig. 4b-c**, all cells shown in **Extended Data Fig. 9a-b** and **Supplementary table 2**).

**Figure 4.**
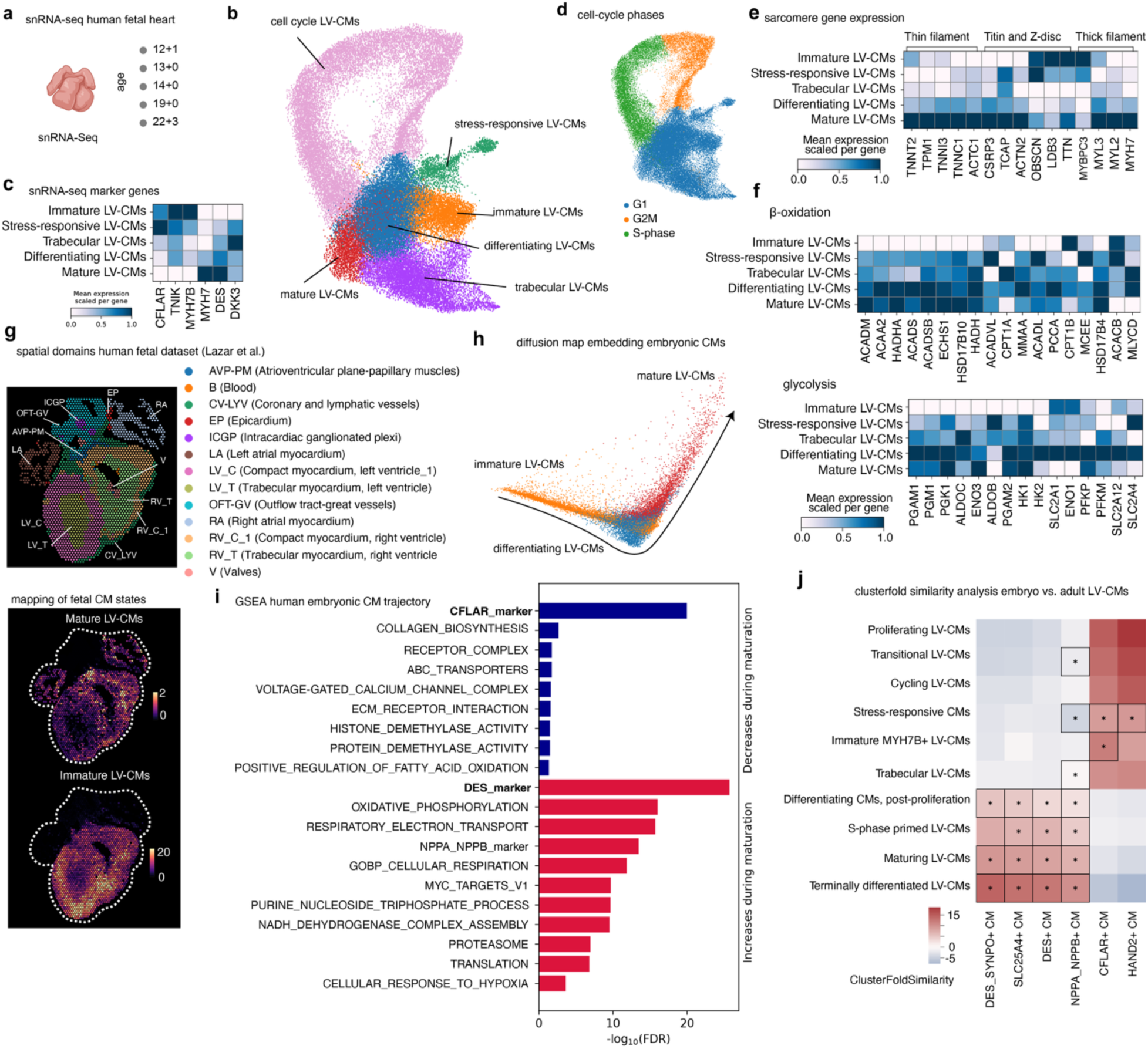
Single-nucleus and spatial dissection of human embryonic cardiomyocyte differentiation. **a,** Overview of human fetal heart samples profiled by snRNA-seq across developmental stages (12–22+ post-conception weeks, pcw). **b,** UMAP embedding of snRNA-seq data from human fetal hearts, revealing distinct cardiomyocyte (CM) populations including proliferating, cycling, differentiating, immature, stress-responsive, trabecular, and terminally differentiated left-ventricular (LV) CMs. **c,** Heatmap of selected marker genes defining major embryonic LV-CM states (scaled mean expression). **d,** Cell-cycle phase annotation of embryonic CMs projected onto the UMAP, highlighting G1, S-phase, and G2/M populations. **e,** Expression of sarcomeric gene modules (thin filament, titin/Z-disc, thick filament) across LV-CM maturation states. **f,** Metabolic gene expression programs associated with β-oxidation and glycolysis across LV-CM states. **g,** Reference spatial domains from the human embryonic heart dataset (Lazar et al.), annotated by anatomical and functional regions. Spatial mapping of embryonic CM states identified in this study onto the reference human embryonic heart sections, showing regional enrichment of immature, mature, and trabecular LV-CMs, alongside major non-CM lineages (estimated counts per Visium spot). **h,** Diffusion map embedding depicting the continuous differentiation trajectory from immature through differentiating to mature LV-CMs. **i,** Gene set enrichment analysis (GSEA) along the human embryonic CM trajectory, highlighting pathways decreasing (blue) or increasing (red) during maturation. **j,** Cluster-to-cluster similarity analysis comparing embryonic LV-CM states (y-axis) to adult human LV-CM populations (x-axis), revealing selective correspondence between embryonic differentiation states and adult CM phenotypes. Stars indicate significance (p-value <0.05) with an hypergeometric test.

Compared to other fetal heart datasets^73,74^, our data for the first time revealed all stages of fetal CM proliferation (**Fig. 4d**). The subclusters cycling and proliferating CMs mapped to the cell-cycle S-phase and G2/M-phase respectively (**Fig. 4d**, **Extended Data Fig. 9b**). We next compared our novel human adult CM states with the cell-cycle scores (S and G2/M) of our human fetal snRNA data (**Extended Data Fig. 9c-d**). While we found the vast majority of adult human CMs in G1-phase and 1% in G2M, none of them were enriched in cell cycle genes to the same level as the proliferating cells in the fetal heart (**Extended Data Fig. 9d**).

In addition to these cell-cycle CMs, which we grouped for downstream analysis, we were able to identify CMs in multiple stages of maturation based on the canonical maturation markers *HOPX*^75^ and *TCAP*^76^ (**Fig. 4b**, **Extended Data Fig. 9e**). Gene-set enrichment analysis revealed an increase of cell-proliferation terms in cycling CMs, WNT pathway terms and glycolysis for immature CMs and myofibril-assembly and cardiac contraction for mature CMs (**Extended Data Fig. 9f**), which corroborated our annotation. Interestingly, we could detect a similar shift in sarcomere gene expression of fetal immature vs. mature CMs (**Fig. 4e**) and changes in β-oxidation and glycolysis regulating genes compared to adult human CFLAR⁺DES⁺ and DES⁺CFLAR⁺ CM (**Fig. 4f**). In order to study the spatial arrangement of these fetal CM states we mapped their location in the tissue using a publicly available spatial transcriptomics dataset^74^. We saw the expected enrichment of our atrial and trabecular cells in their respective compartments (**Extended Data Fig. 9g-h**). Immature and mature CM were more evenly spread throughout the heart, although there was a higher abundance of immature cells around the right ventricle and a higher abundance of mature cells in atrioventricular plane papillary muscle, which might relate to differing mechanical strain in these distinct tissue regions (**Fig. 4g, Extended Data Fig. 9g**).

Next we investigated gene expression changes during CM maturation. We partitioned our data into immature, differentiating and mature CMs and calculated a diffusion map, to visualize the continuous trajectory of the maturation process (**Fig. 4h**). By assigning cells a pseudo-temporal ordering we were then able to study expression trends along this trajectory. We found that an increase of the DES⁺ and a decrease of CFLAR⁺ signatures, derived from the adult snRNA dataset, were among the most significant results (**Fig. 4i**). Maturation was also accompanied by increased cellular respiration and oxidative phosphorylation (**Fig. 4i**).

In the next step we used ClusterFoldSimilarity^60^ and a hypergeometric test to compare the embryo subclusters to the adult human CMs. We found that adult HAND2⁺ and CFLAR⁺ were significantly enriched in proliferating fetal CMs (**Fig. 4j**). Mapping of fetal cell types into the PHLOWER cell differentiation tree based on adult CMs confirmed that immature LV-CMs are related to the CFLAR+ CM branch **(Extended Data Fig. 9i**).

Overall, our data uncover a conserved fetal differentiation trajectory of CMs that provides a reference framework for adult CM states in human heart disease. Polyploid HAND2⁺ and CFLAR⁺ CMs transcriptionally revert toward immature fetal programs, whereas diploid DES⁺ and DES–SYNPO⁺ CMs represent terminally differentiated, contractile end states. Notably, polyploid CMs do not reacquire fetal proliferative signatures, supporting a model of incomplete fetal reprogramming rather than regenerative proliferation.

### Temporal disease association and HF targets in CMs

To study the novel CM heterogeneity and their dynamics with spatiotemporal resolution, we performed snRNA-seq and Xenium in mice after LAD ligation across four post-MI time points from day 3 to day 28 (**Fig. 5a**). In total, we recovered 75,825 nuclei from snRNA-seq and 543,586 nuclei from spatial transcriptomics (**Fig. 1g; Supplementary Table 2**), which allowed the detection of six transcriptionally distinct cardiomyocyte states (**Fig. 5b**). Marker-gene expression of the novel human CM clusters demonstrated conservation in the mouse dataset (**Fig. 5c, Extended Data Fig. 10a**), with a major cluster high in *Des* and a second one expressing fetal genes like *Myh7b*, although *Cflar* was expressed at much lower levels than in the human.

**Figure 5.**
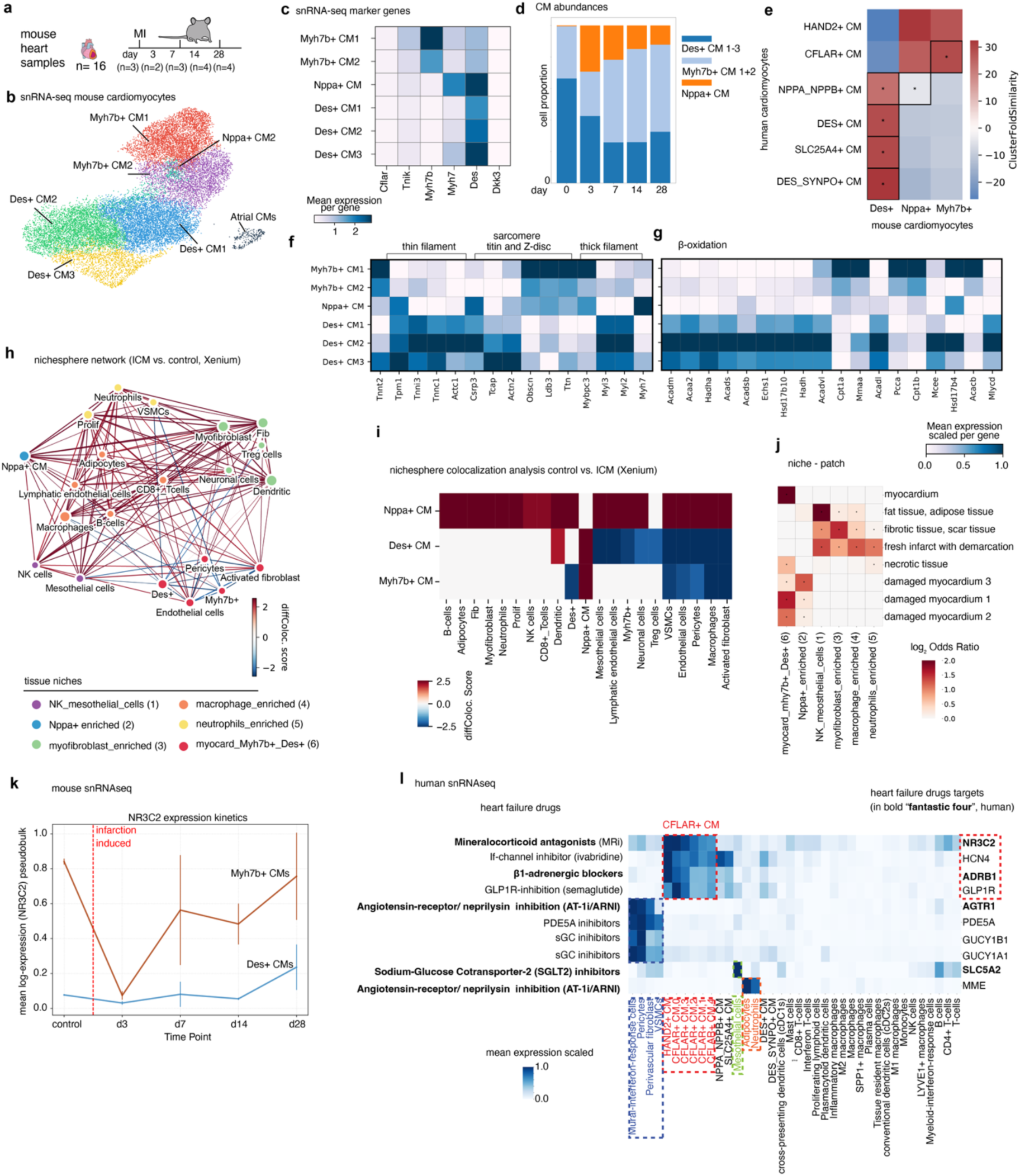
Spatiotemporal organization of cardiomyocyte states after myocardial infarction. **a,** Experimental design and sampling scheme of murine hearts subjected to myocardial infarction (MI), with snRNA-seq performed at days 0, 3, 7, 14, and 28 post-MI. **b,** UMAP embedding of snRNA-seq profiles from murine cardiomyocytes, identifying Myh7b⁺, Nppa⁺, and Des⁺ cardiomyocyte (CM) subclusters, as well as atrial CMs. **c,** Heatmap of marker gene expression defining CM subclusters, highlighting differential expression of contractile, stress-response, and cytoskeletal genes. **d,** Temporal dynamics of CM subcluster abundances following MI, showing early expansion of Nppa⁺ CMs and progressive enrichment of Des⁺ CM states during remodeling. **e,** Cross-species similarity analysis mapping murine CM subclusters to human CM states, including HAND2⁺, CFLAR⁺, NPPA/ NPPB⁺, DES⁺, SLC25A4⁺, and DES_SYNPO⁺ cardiomyocytes. **f,** Scaled mean expression of sarcomeric gene modules (thin filament, titin/Z-disc, thick filament) across CM subclusters, indicating progressive structural remodeling. **g,** Scaled mean expression of β-oxidation–related genes, revealing reduced oxidative metabolic signatures in Myh7b⁺ and Nppa⁺ CMs relative to Des⁺ CM states. **h,** Nichesphere interaction network derived from Xenium spatial transcriptomics comparing ischemic cardiomyopathy (ICM) and control myocardium, highlighting CM-centered tissue niches enriched for immune, stromal, and vascular cell interactions. **i,** Differential spatial colocalization scores for CM subclusters with non-myocyte cell types in ICM versus control hearts, indicating selective niche rewiring of Nppa⁺ and Des⁺ CMs. For **h** and **i**, only significant interactions are shown(adjusted Wilcoxon’s rank sums test; p-value < 0.05). **j,** Enrichment of CM-associated tissue patches across anatomical and pathological regions, including fibrotic scar, infarct border zones, necrotic tissue, and damaged myocardium. Significance was determined by Fisher’s exact test with adjusted p-values < 0.05. **k,** Temporal dynamics of heart-failure–relevant drug targets across murine CM states following MI. **l,** Expression landscape of clinically established heart-failure drug targets across human cardiac cell types, with the “fantastic four” therapeutic classes highlighted, demonstrating preferential target enrichment in distinct CM and non-CM compartments.

We next quantified shifts in CM state abundances across the post-MI time course^77^. Both Des⁺ and Myh7b⁺ CMs were reduced at day 3 following the acute MI, whereas NPPA_NPPB⁺ CMs represented the only CM population that increased at this early injury time point (**Fig. 5d, Extended Data Fig. 10b**). By day 7, the trajectories of Des⁺ and Myh7b⁺ CMs diverged: Des⁺ CMs remained low, while Myh7b⁺ CMs expanded progressively and became the dominant CM population at later stages (**Fig. 5d**). This delayed but pronounced enrichment of Myh7b⁺ CMs after injury highlights them as an adaptive CM state that emerges during post-ischemic remodelling.

To link mouse and human CM states, we applied ClusterFoldSimilarity^60^ and enrichment test, which showed that the Des⁺ CM nuclei from mouse overlapped in their expression with diploid CMs of the adult human heart (DES⁺, SLC25A4⁺, DES_SYNPO⁺, NPPA_NPPB⁺), whereas the Myh7b⁺ CM mouse nuclei were more similar to human polyploid HAND2⁺ and CFLAR⁺ CM nuclei (**Fig. 5e**). Nppa⁺ mouse CMs overlapped significantly with NPPA⁺ human CMs but also shared markers with human HAND2⁺ and CFLAR⁺ CMs (**Fig. 5e**). We then compared the mouse clusters to our fetal human data. Des+ CMs were most similar to terminally differentiated CM, while Myh7b⁺ CM showed correspondence to fetal immature MYH7B+ cells, a similar trend to what we observed in our adult to fetal comparison (**Extended Data Fig. 10c**).

Next we compared transcriptional programs between mouse CM subtypes (**Fig. 5f-g and Extended Data Fig. 10d–i**). Sarcomere-associated genes, including *Obscn*, *Ldb3*, *Ttn* and *Mybpc3*, were significantly upregulated in Myh7b⁺ CMs, whereas genes involved in β-oxidation regulation were upregulated in Des⁺ CMs mirroring the human polyploid pattern (**Fig. 3f-g, 5f-g**). Analyses of glycolytic and glutaminolysis genes (**Extended Data Fig. 10d-e**) showed that Myh7b⁺ CMs demonstrated reduced gene expression consistent with a metabolically rewired, stress-adapted state, whereas Des⁺ CMs maintained a more oxidative, metabolic increased profile (**Extended Data 10d-f**). Because our human data implicated polyploid CMs in fibrotic and developmental signalling, we next systematically assessed these pathways in the mouse model (**Extended Data Fig. 10g-i**). Our analysis shows that canonical profibrotic ligands and receptors including Tgfb isoforms, *Tgfbr1/2, Pdgfa/b, Pdgfra/b, Edn1/Ednra/Ednrb*, *Vegfa* and *Il6/11*,were strongly enriched in Nppa⁺ and Myh7b⁺ CMs compared to Des⁺ CMs, while Des+ CMs preferentially expressed *Vegfb* and *Vegfc* (**Extended Data Fig. 10g-i**). This pattern is only partially concordant with the human heart, where polyploid CFLAR+/HAND2+ CMs demonstrated higher upregulated gene expression of fibrosis signalling ligands and receptors (**Extended Data Fig. 8b, 10g-i**). However a marked bias of WNT (particularly WNT5/11) and Hedgehog signaling components towards Myh7b⁺ CMs remained: Myh7b⁺ cells expressed multiple WNT ligands and receptors, NOTCH ligands and receptors, and Sonic hedgehog pathway members, whereas Des⁺ CMs predominantly expressed inhibitory modulators such as *Sfrp1* and *Dkk3* (**Extended Data Fig. 8g+i).**

Intracellular pathway analysis of mouse CM states revealed notable differences compared to human CMs (**Extended Data Fig. 10j**). Specifically, WNT, TGFβ, and Notch signaling were predominantly upregulated in murine Nppa+ CMs rather than Myh7b+ CMs, highlighting a species- or model-dependent divergence in signaling. Despite this, the general functional stratification was conserved: the adaptive Nppa+ state in mice concentrated profibrotic and developmental signaling capacities, whereas Des+ CMs retained a restrained, homeostatic profile similar to that of humans (**Extended Data Fig. 10j**).

Next, we performed spatial colocalization analysis, using NicheSphere of Xenium mouse data, contrasting controls and ICM conditions (days 7,14 and 28), which detected six major tissue niches (**Fig. 5h**). In the mouse heart, Des⁺ and Myh7b⁺ CMs largely shared a common myocardial niche (niche 6 - myocardium) indicating that diploid and polyploid CMs coexist within the same anatomical compartments (**Fig. 5h-i**). Cells in this niche tended to lose their colocalization to other cell-types upon diseases. By contrast, NPPA⁺ CMs dominated a distinct niche that showed a gain of colocalization with myofibroblasts in ICM, consistent with their transcriptional injury response program (**Fig. 5i**). This organisation differs from the human heart, where Des⁺ and polyploid CFLAR⁺/HAND2⁺ CMs segregate into two spatially distinct niches suggesting that niche partitioning of diploid versus polyploid CMs is, at least in part, time- and species-dependent. NPPA⁺ mouse CMs in niche 2 mapped predominantly to the “damaged myocardium 3” region in the heart patch atlas, in which also the human NPPA_NPPB CMs enriched (**Fig. 5j**). This underscores conserved features of the injury-associated NPPA+ states across myocardial tissue from different mammalian species. Next we performed a PILOT trajectory analysis on spatial niches, which indicates the decrease of myocardium (niche 1) and increase of Nppa+ (niche 2) and the myofibroblast_enriched niche (niche 3) among others and revealing an overall increase in periostin expression fitting to our previous data (**Fig. 5i, Extended Data Fig. 11a-b**).

Finally, we analyzed the expression of heart failure drug targets in our mouse snRNA data (**Extended Data Fig. 11c**). This revealed a conserved, dichotomous expression pattern: three out of four major heart-failure targets, β1-adrenergic receptor (*Adrb1*), mineralocorticoid receptor (*Nr3c2*) and angiotensin II receptor (*Agtr1*) were predominantly expressed in Myh7b⁺ CMs, while *Pde5a* and *Gucy1a1* were enriched in pericytes and vascular smooth muscle cells (**Extended Data Fig. 11c**). *Glp1r* expression was mainly restricted to T cells. Adrenoceptor (β1) expression overall was higher in Myh7b⁺ than in Des⁺ CMs. Comparing the dynamics of mineralocorticoid receptor (*Nr3c2*) expression, we noted a marked expression dynamics only in Myh7b+ CMs (**Fig. 5k).** We next asked if this pattern of HF drug target expression is recapitulated in our human snRNA data and curated a panel of established drug targets and well-supported HF mediators and profiled their expression across all cardiac cell subtypes (**Fig. 5l**). Of the four standard of care drugs - β1-blockers, ACEi/ARB, mineralocorticoid-receptor (MR) antagonists, and SGLT2 inhibitors, we observed a selective enrichment of transcripts encoding two of these targets (MR and adrenergic receptor β1) in polyploid CMs (HAND2+/CFLAR+) (**Fig. 5l, Extended Data Fig, 11d).** Additionally, we found a significant enrichment of *GLP1R* in polyploid CMs, which in mice was restricted to Treg and CD8-T-cells (**Fig. 5l, Extended Data Fig. 11c**). *AGTR1* was primarily expressed in cardiac pericytes and fibroblasts (**Fig. 5l**) as well as transcripts for *PDE5A* and the soluble guanylyl cyclase α1 subunit (*GUCY1A1*).

Taken together, our HF target gene analysis indicates that polyploid CMs constitute a major —previously unrecognized— target cell for heart failure therapies, while pharmacologic control points for the cGMP pathway are predominantly enriched in the vascular/perivascular compartment, and not the endothelium. In sum, our analysis suggests a compartment-specific division of therapeutic action in human heart failure treatment, with polyploid CMs as a major myocardial drug-addressable population.

### TNIK inhibition ameliorates maladaptive remodelling by targeting polyploid CMs

Based on prior evidence implicating TNIK in WNT-dependent fibrotic programmes^78^, we examined TNIK expression across our multispecies snRNA-seq atlas and observed consistent enrichment in CFLAR+ or Myh7b+ CMs across human, mouse, and rat (**Extended Data Fig. 12a-d**). While TNIK expression was not strictly CM-restricted, being detectable in human mast cells and select murine T-cell populations, the rat heart displayed the highest CM specificity (**Extended Data Fig. 12d**). This enrichment suggested a potential role for TNIK in regulating polyploid CM states.

To investigate the functional consequences of TNIK inhibition in CMs, we assessed its effects both *in vitro* using human iPSC-derived CMs (**Fig. 6a-g)** and *in vivo* in a rat MI model (**Fig. 6h-p, Extended Data Fig. 12e-j**). Long-term differentiated beating CMs from two independent iPSC lines were treated for seven days with the TNIK inhibitor NCB-0846^79^ or vehicle and profiled by snRNA-seq. Unsupervised clustering identified two major CM populations (**Fig. 6b**), which we annotated as mature and immature based on enrichment analysis (**Fig. 6c**). The more immature CM showed the CFLAR signature, while the mature cluster closely resembled DES⁺ CM. ClusterFoldSimilarity^60^ and a hypergeometric test verified the similarity to DES⁺ and CFLAR+ clusters from the adult dataset (**Fig. 6d)**.

**Fig. 6.**
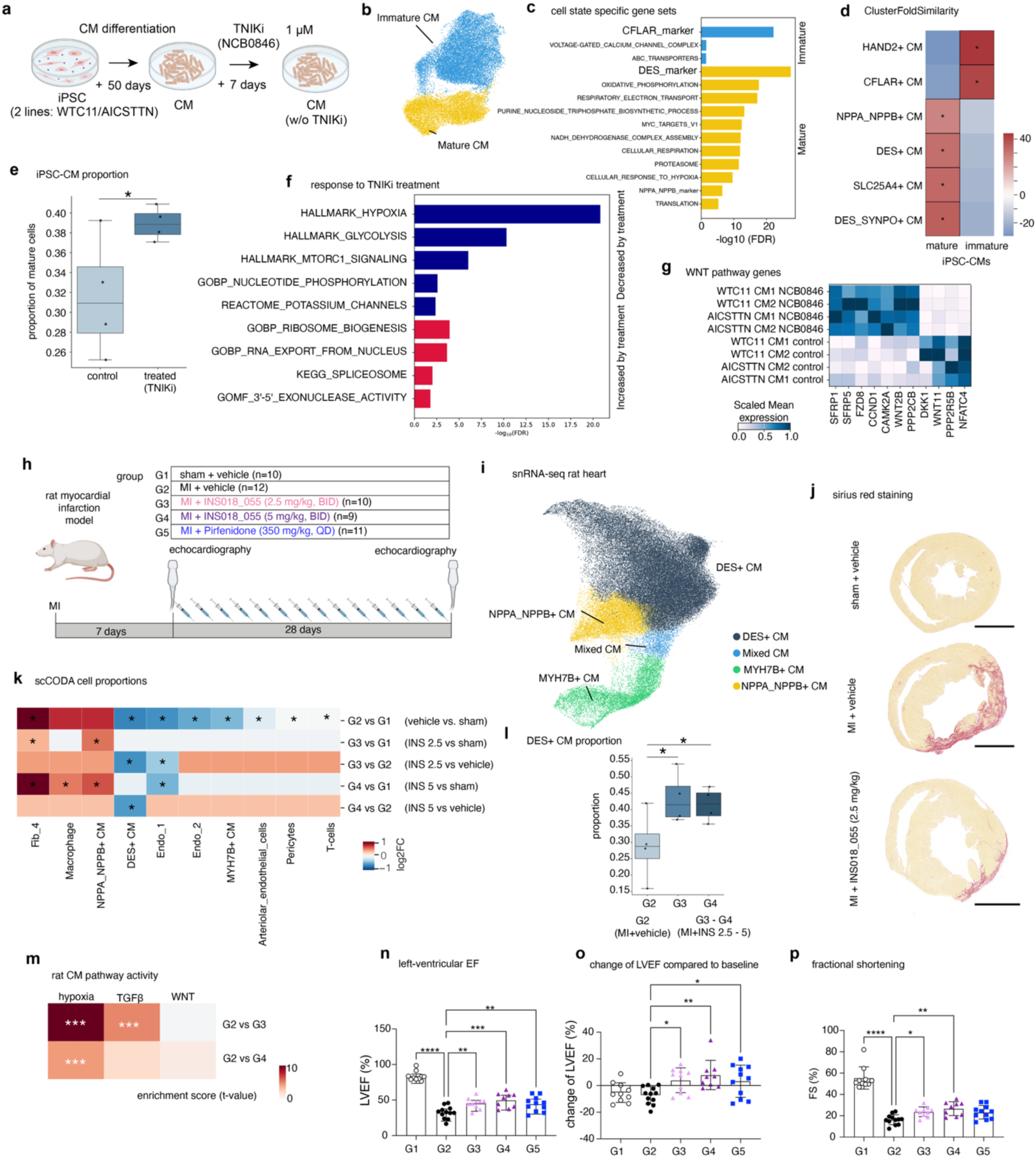
TNIK inhibition promotes cardiomyocyte maturation and improves post-MI cardiac function. **a,** Experimental workflow for iPSC-derived cardiomyocyte (CM) differentiation and TNIK inhibition with NCB-0846. **b,** UMAP of iPSC-CMs showing segregation into immature and mature CM states. **c,** Gene set enrichment analysis highlighting immature (CFLAR-associated) versus mature (DES-associated) CM programs. **d,** Transcriptomic similarity between iPSC-CMs and adult clusters, color indicates ClusterFoldSimilarity, *p-value < 0.05 hypergeometric test. **e,** Increased proportion of mature CMs following TNIK inhibition in iPSC-CM, * indicates FDR < 0.1 for the scCODA abundance test. **f,** Pathways decreased or increased upon TNIK inhibition in iPSC-CMs. **g,** Expression of WNT pathway genes across control and TNIK-inhibited iPSC-CMs. **h,** Rat myocardial infarction (MI) study design and treatment groups. **i,** snRNA-seq of rat hearts identifying CM subpopulations after MI and treatment. **j,** Sirius Red staining and quantitative analysis demonstrating reduced fibrosis with TNIK inhibition, scale bars indicate 4 mm. **k)** Changes in cell abundance between treatment groups,* indicates FDR < 0.1 for the scCODA abundance test. **l)** Increased proportion of mature CMs following TNIK inhibition in rats, * indicates FDR < 0.1 for the scCODA abundance test. **m)** Compatarative enrichment analysis of selected pathways. ***p-value < 0.001. **n-p),** Improved cardiac function following TNIK inhibition, including left ventricular ejection fraction, and fractional shortening. Data are shown as mean ± s.e.m.; *p-value < 0.05, **p-value < 0.01, ***p-value < 0.001.

Cell-type abundance analysis revealed a significant increase in mature CM states following TNIK inhibition (**Fig. 6e**). Gene set enrichment analysis demonstrated a marked reduction in hypoxia-associated transcriptional programmes, metabolic changes and changes in RNA transport and splicing after TNIK inhibition (**Fig. 6f**). Although global WNT pathway activity was not significantly altered (**Extended Data Fig. 12g**), TNIK inhibition resulted in coordinated expression changes of non-canonical WNT components, including decreased *WNT11* and *NFATC4* expression and induction of the WNT antagonists *SFRP1* and *SFRP5* (**Fig. 6g**). Together, these findings indicate that TNIK inhibition promotes maturation-associated CM states alongside reduced hypoxia signalling and selective remodelling of WNT-related transcriptional programmes.

We next investigated whether TNIK inhibition confers therapeutic benefit in the injured heart in vivo. Rats underwent permanent LAD ligation or sham surgery and, following a one-week recovery period, received vehicle, the clinical-stage TNIK inhibitor INS018_055 (rentosertib, NCT05938920, NCT05975983) at two doses, or the antifibrotic agent pirfenidone as a reference therapy for 28 days (**Fig. 6h**). Rentosertib is a novel AI-designed TNIK inhibitor whose efficacy was recently demonstrated in a phase IIa trial for idiopathic pulmonary fibrosis^81^. Single-nucleus RNA-seq was performed on FFPE rat heart tissue at endpoint and identified similar CM populations to the human and mouse infarct datasets (**Fig. 6i)**. TNIK inhibition resulted in a reduction of fibrosis, as shown by sirius red staining (**Fig. 6j, Extended Data Fig. 12h**). Abundance analysis revealed a significant, dose-dependent increase in DES⁺ cardiomyocytes following TNIK inhibition, accompanied by a reduction in fibroblast and macrophage populations (**Fig. 6k-l**). Furthermore, treatment with both doses of rentosertib resulted in a marked decrease of hypoxia and TGFβ signalling pathway in CMs (**Fig. 6m**), while WNT signatures did not significantly change between any of the groups. iPSC-derived CMs and rat CM shared a large overlap in genes that were down-regulated upon treatment. More than 400 hypoxia-associated genes were decreased in both cohorts, including transcripts implicated in cross-talk between HIF-1α signalling and WNT/β-catenin or TGF-β pathways, such as KCTD11 (REN), SEMA4B, PDK1, TGFBI and ARRDC3. Rats treated with rentosertib additionally demonstrated improved cardiac function as detected by echocardiography, which reached comparable levels to the group treated with antifibrotic control pirfenidone (**Fig. 6n-p, Extended Data Fig. 12i-j**). Collectively, the concordant human iPSC-CM and rat MI data demonstrate that pharmacological TNIK inhibition mitigates maladaptive post-infarction remodelling, in part by shifting (*in vitro*) or leading (*in vivo*) to increased mature CM transcriptional states and suppressing hypoxia- and fibrosis-associated signalling programmes. Together, these results support the concept that pharmacological modulation of disease-associated polyploid CM states can favorably modulate post-infarction remodelling of the heart.

## Discussion

In this study, we provide a multi-scale, multimodal view of how CM heterogeneity, ploidy state and microenvironmental context shape cardiac remodeling in human heart failure. We integrate single-nucleus transcriptomics, chromatin profiling, high-resolution spatial transcriptomics and histology across ischemic and non-ischemic cardiomyopathies, human development, and in three mammalian species. Our analysis revealed that disease progression is not only encoded in changes of cell-type composition and fibrosis, but in the emergence of distinct diploid and polyploid cardiomyocyte states that shape spatial niches and drive cardiac remodeling.

We systematically dissect diploid versus polyploid CMs in the human heart. Using DAPI-based nuclear profiling, FANS-enriched ploidy fractions and integrative snRNA-seq, we show that polyploid CMs form distinct transcriptional states, marked by CFLAR and HAND2, which are enriched in 4N/8N nuclei, engage specific metabolic and chromatin-remodeling programs, and display a disease-biased remodeling signature. In contrast, predominantly diploid DES⁺ CMs preserve oxidative phosphorylation–dominated metabolic profiles and mature sarcomere programs. Direct comparison to human fetal hearts indicates that HAND2⁺ and CFLAR⁺ polyploid CMs align with proliferating and immature fetal states, whereas DES⁺ and DES_SYNPO⁺ CMs resemble terminally differentiated, contractile end states. Together, these findings provide a cellular and ploidy-based resolution to the fetal gene hypothesis. We demonstrate that developmental reactivation in human heart disease is partial, selective, and strongly shaped by CM ploidy, rather than representing a generic fetal-like state.

At the tissue level, CM states organize into distinct spatial niches that differ between ischemic and non-ischemic cardiomyopathies. Polyploid CFLAR⁺/HAND2⁺ CMs preferentially localize to healthy-appearing myocardium and damaged myocardium, whereas DES⁺ and DES_SYNPO⁺ states colocalize with myofibroblasts and tissue-resident macrophages in fibrotic and chronically remodeled regions. The use of deep learning models for histology analysis revealed distinct damaged cardiomyocyte regions conserved in both mouse and human, that could not be identified even by expert pathologists. These distinct areas have differences in both their abundance of damaged NPPA_NPPB CMs and lymphocytes.

Subcellular spatial transcriptomics further reveals a discrete ICD transcriptome compartment, enriched for mRNAs encoding junctional regulators, cytoskeletal linkers and local mRNA-regulatory components. Differential analysis of this ICD program across cardiomyocyte states and ploidy suggests that altered mRNA localization and junctional remodeling may contribute to conduction and mechanical dysfunction in heart disease. All these technical advances are released as community resources, i.e. machine learning models that empower researchers to segment CMs in microscopy and spatial transcriptomic datasets, identify regions of interest in H&E sections, and detect cardiomyocyte polyploidy from DAPI images.

Ligand–receptor and drug-target mapping position these CMs states within a structured signaling and therapeutic landscape. Polyploid CFLAR⁺ and HAND2⁺ CMs act as major receivers and senders of distinct developmental and fibrotic cues, with enriched Notch, non-canonical Wnt and Hedgehog receptors, as well as TGFβ and ERBB signaling components. More differentiated DES⁺, NPPA_NPPB⁺ and SLC25A4⁺ CMs preferentially express Wnt antagonists.

Analysis of heart failure drug targets demonstrated that molecular targets of standard-of-care therapies, such as β-blockers and mineralocorticoid receptor antagonists, show marked enrichment in polyploid CMs. These data suggest a compartmentalized division of therapeutic action and highlight polyploid CMs as a major drug-addressable myocardial population in patients.

This study is limited by the observational nature of human end-stage cardiac tissues, targeted spatial gene panels, and reliance on transcript-based inference for signaling and drug-target activity. In addition, ploidy assignment is based on DAPI content and enrichment rather than direct lineage tracing. Furthermore, our pharmacological perturbation of TNIK in an *in vivo* model is not CM-subtype specific. Future work combining CM-subtype-specific genetic perturbations, higher-order multiplexing spatial profiling, and functional assays of electrophysiology and force generation will be required to test causal roles of individual CM states and to define how genome multiplication mechanistically constrains -or enables- cardiomyocyte plasticity.

Finally, through cross-species analyses we nominated TNIK as a central regulator in polyploid cardiomyocyte remodeling. TNIK inhibition in a rodent model of myocardial infarction ameliorated remodelling, reduced fibrosis and improved myocardial function. We anticipate that these insights will serve as a reference to dissect when and how diploid CMs transition into maladaptive polyploid states, and to guide the design of interventions enabling the development of novel cardiac therapeutic strategies, such as TNIK inhibition, that selectively reprogramming pathological CM states to restore myocardial function.

## Methods

### Experimental procedures

All human tissue protocols were approved by the local ethics committees of Ruhr University Bochum (Bad Oeynhausen; Reg.-No. 21/2023), RWTH Aachen University (EK 24-337), University of Chicago (STU00216865), University Medical Center Utrecht (UCC-UNRAVEL #12-387; TCBio 25-242), Amsterdam University Medical Center (METC 2016_285; Dutch Fetal Biobank^81^), and Washington University in St. Louis (No. 201104172). Human myocardial tissue was obtained from non-transplanted donor hearts, from patients with myocardial infarction undergoing heart transplantation, and from patients receiving a total artificial heart or left ventricular assist device (LVAD) implantation. The study complied with the Declaration of Helsinki and with the Dutch Code of Conduct for the Responsible Use of Human Tissue. All participants provided written informed consent. This study was conducted within the framework of the UCC-UNRAVEL biobank^82^ (University Medical Center Utrecht; www.unravelrdp.nl). Tissue collection was approved by the scientific advisory boards of the UMC Utrecht Biobank (UNRAVEL) and the Amsterdam University Medical Center (Dutch Fetal Biobank).

### Human tissue processing and screening

Mainly FFPE heart tissues were used. Most samples were obtained between 2021 and 2023. 5 µm tissue sections were performed and the H&E stainings were analyzed by a cardiac pathologist. For RNA quality control, two scrolls of 25 µm tissue sections were used as input for the Qiagen FFPE RNA isolation kit. DV200 measurements were subsequently performed using TapeStation (Agilent). For MERFISH spatial transcriptomics (Vizgen) all samples underwent QC and only samples with a DV200 value of 30% or higher were included. For Xenium and Flex (10XGenomics) no DV200 measurements were performed. For H&E and IF imaging a Zeiss Axioscan was used.

### Animal studies

Mouse experiments were performed according to the animal welfare guidelines and German national laws and were authorized by the local authority (Regierungspräsidium Darmstadt, Hessen, Germany and Baden-Württemberg, Germany) under the animal experiment license FU/2084. C57BL/6J wildtype animals were obtained from Janvier Labs. Female and male mice were used at the age of 10 to 14 weeks. Myocardial infarction was induced by ligation of the left anterior descending coronary artery as described previously^83^. Experiments were performed in both females and males. Animals were held at 23°C ambient temperature and 60% humidity in 10 h/14 h light/dark cycle. At the end of each experiment, animals were euthanized by cervical dislocation after isoflurane inhalation (3–3.5 vol%).

Rat studies were performed by WuXi AppTec (Nantong, China, IACUC protocol number GP02-QD079-2023v1.2). Sprague–Dawley rats (male, 6-8 weeks, purchased from Vital Rivers Animal Technology Co., Ltd., weighted around 200-300 g) were obtained and housed at the WuXi AppTec (Nantong) facility under controlled conditions (22–25 °C, 40–70% humidity). A timed lighting system maintained a 12 h light/12 h dark cycle (06:00–18:00). Rats had ad libitum access to chow and water. Left anterior descending coronary artery (LAD) ligation was performed on SD rats to induce heart failure. On the day of model induction, rats were anesthetized with a combination of Zoletil (50 mpk, i.m.) and xylazine (8 mpk, i.p.). The trachea was connected to a ventilator for assisted respiration. A thoracotomy was performed between the third and fourth ribs; the pericardium was opened, and the LAD was ligated with 6-0 silk suture. The ribs and skin were closed with 4-0 silk sutures, and rats were placed on a heating pad postoperatively until recovery from anesthesia. In the sham group (G1), all procedures were identical except the ligation of the LAD. Animals were administered meloxicam (1 mg/kg, i.m.) and gentamicin hydrochloride (8 mg/kg, i.p.) postoperatively for analgesia and infection prophylaxis. One week later, successfully modeled rats (30% or more reduction in LVEF) selected by echocardiographic measurement, were grouped by LVEF into four groups G2 (Model), G3 (INS018-055, 2.5 mg/kg, BID), G4 (INS018-055, 5 mg/kg, BID), and G5 (pirfenidone, 350 mg/kg, QD). Echocardiography was performed on Vevo®1100 (Fuji), readout includes Left ventricular ejection fraction (LVEF), fractional shortening (FS), left ventricular end-diastolic diameter (LVEDD), left ventricular end-systolic diameter (LVESD), left ventricular end-diastolic volume (LVEDV), left ventricular end-systolic volume (LVESV), stroke volume (SV), cardiac output (CO), left ventricular mass (LV mass), left ventricular anterior wall thickness in systole/diastole (LVAWTs/d), and left ventricular posterior wall thickness in systole/diastole (LVPWTs/d). After grouping, rats received vehicle or test drugs by oral gavage for 4 weeks: G1-G4 were dosed twice daily (12 h apart), and G5 once daily. INS018-055 was given in 50 mM citrate buffer, pirfenidone in 0.5% CMC-Na. General condition was monitored and abnormalities recorded throughout. After 4 weeks of treatment, endpoint echo measurements and sample collection were performed. The hearts were fixed in 4% paraformaldehyde (PFA) and subjected to Sirius Red (SR) staining to evaluate myocardial fibrosis. Sirius red staining was analyzed by Halo AI 3.6. 4 samples each from groups 1-4 were then selected for snRNA processing.

### Human iPSC culture

WTC-11 (Male, Coriell, GM25256) and AICS-TTN (Male, Coriell, AICS-0048-039) human iPSC lines were used in this study. iPSCs cultures were maintained in Matrigel coated plates (Corning, 356231) in mTeSR™ Plus medium (STEMCELL, 100-0276) and passaged upon reaching 70% confluency. Briefly, iPSCs, with no signs of differentiation, were dissociated chemically using TryPLE Select (ThermoScientific, 12563029) and seeded at 1:4 ratio onto Matrigel-coated plates with mTeSR™ Plus medium, supplemented with 10 μM Y-27632 dihydrochloride (MedChemExpress, HY-10071). iPSCs cultures were regularly tested for mycoplasma contamination.

### Human iPSCs derived cardiomyocyte differentiation protocol

iPSCs cultures at 70-80% confluency and with no apparent signs of spontaneous differentiation, were differentiated to cardiac lineage by following established conditions^84,85^. Briefly, to promote gastrulation on the iPSC cultures at Day 0, the culture media, switched to RPMI 1640 (Gibco, 21875034) basal media supplemented with 1% penicillin/streptomycin, 2% B27 supplement minus insulin (Gibco, A1895601, hereafter referred to as RPMI/B27−), and 6 μM CHIR99021 (Cayman Chemical, 13122). Culture was directed on Day 2 towards cardiac mesodermal lineage, by supplementing to RPMI/B27− basal media with 5 μM IWP2 (Tocris, 3533). On day 4, the medium was changed to RPMI/B27−. On day 6 and 8, the medium was changed to RPMI 1640 basal media supplemented with 1% P/S and 2% B27 supplement (Gibco, 17504044; hereafter referred to as RPMI/B27+). Cardiomyocytes were enriched until day 10, by switching the medium to glucose-free RPMI 1640 (Gibco, 11879020) supplemented with 0.1% (v/v) sodium DL-lactate (Sigma, L4263-100 mL). On day 13, the medium was reverted to RPMI/B27+, and thereafter, medium changes were performed every 2 days.

### TNIK perturbations on hiPSCs derived cardiomyocytes

At day 50 of differentiation, beating cardiomyocyte cultures were treated with 1 μM NCB-0846 (MedChemExpress, HY-100830), with DMSO-treated cultures serving as controls, for 7 days.The medium was changed every two days^78,86^. On day 7 of treatment, iPSC-derived cardiomyocytes (iPSC-CMs) were mechanically harvested for nuclei isolation. Nuclei were isolated using the 10x Nuclei Isolation kit (10XGenomics) with bulk cardiomyocyte cultures handled according to the tissue protocol.

### Tissue dissociation and nuclei isolation

Two to three 50 µm sections were taken from FFPE samples and stored at 4°C until processing. To isolate single nuclei from FFPE tissue, samples were deparaffinized with Xylene (3x) for 10 min each to remove the paraffin (Sigma-Aldrich, 108298) and rehydrated in an decreasing ethanol concentrations (100%, 70%, 50%, 30%) (prepared from Fisher Scientific, BP2818500), nuclease-free water (invitrogen, 10977035) and 1X PBS (Gibco, 11594516). To digest the tissue sections, they were processed with digestion buffer (1.68 ml RPMI 1640 medium (Thermo Scientific, A1049101), 420 µl Liberase TH (final concentration 1mg/ml, Roche, 5401135001), 21 µl Collagenase D (final concentration 1mg/ml, Roche, 11088858001), 8.4 µl RNase inhibitor (final concentration 0.2 U/µl, New England Biolabs, M0314L)) on the gentleMACS™ Octo Dissociator (Miltenyi Biotec, 130-096-427) with the 37C_FFPE_1 protocol. After digestion, the samples were transferred to glass tubes, cell lysis mix was added (796 µl nuclei EZ lysis buffer (Sigma Aldrich, N3408), 200 µl 10% BSA (Miltenyi Biotech, 130-091-376), 4 µl RNase inhibitor), pestled with A and B (10-15x), and incubated for 5 min to dissociate the tissue further. Afterwards, the nuclei/cell solution was filtered with a 70 µm filter (PN: 431007060, PluriSelect) and washed. To reduce debris, a density centrifugation step was performed at 700 rcf for 10 min at 4°C in debris removal buffer (550 µl debris removal reagent, PN-2000560, Chromium Nuclei Isolation Reagents (PN-1000447), 10X Genomics), 0.55 µl reducing agent B (10X Genomics, Reducing Agent B (PN-1000450), PN-2000087)). The nuclei suspension was then washed and resuspended in resuspension buffer (2733.5 µl 0.5X PBS, 5.5 µl 10% BSA, 11 µl RNase inhibitor). Nuclei concentration was determined by staining an aliquot of the nuclei suspension with ethidium homodimer 1:100 (Invitrogen, E1169) and counting on the Invitrogen™ Countess™ 3 FL Automated Cell Counter (Fisher Scientific, 16832556). Only samples with over 100,000 nuclei were input for probe hybridization.

### Single-Nucleus RNA- sequencing (Flex)

Single-nucleus RNA sequencing (snRNA-seq) was performed following the 10x Genomics protocol CG000527 (Rev. E). Fixed nuclei or cells were hybridized and incubated overnight (16-24 h) with mRNA transcript probe sets targeting approximately 18,000 genes per species, with probes carrying a sample-specific barcode in the left handle. After hybridization, nuclei were counted and pooled in equal proportions, followed by multiple washes with post-hybridization wash buffer (10X Genomics) to remove unbound probes. The pooled nuclei were then counted again, and a volume corresponding to a target recovery of 20,000 nuclei or cells per sample was combined with a master mix containing enzymes for probe ligation and extension during GEM incubation. The reaction mixture was loaded onto a NEXT GEM Chip Q together with gel beads and partitioning oil, and GEMs were generated using the Chromium X system. Following GEM recovery, cDNA was amplified and used for library preparation. A fraction of the amplified material was subsequently re-amplified using library-specific P5 and P7 indices.

### QC and Sequencing

All libraries were quantified with the Qubit fluorometer’s high sensitivity dsDNA reagents (Thermo Fisher Scientific) and sized on the Agilent 2200 Tapestation with high-sensitivity tapes and reagents. All libraries were run on the NextSeq2000 and NovaSeq X (Illumina). Sample sequencing was conducted to achieve a depth of 20,000 reads per nuclei/cell as recommended by 10XGenomics.

### Fluorescent-activated nuclei sorting (FANS)

Nuclei were stained with DAPI (1:10,000, 1 mg/dl stock, Sigma-Aldrich) and diluted in 1X PBS and analyzed on the FACS SH800 (SONY) using standard laser configuration and a 100 µm microfluidic chip (SONY) for sorting. Events were first gated on FSC and SSC profiles, followed by singlet gating using FSC-H and FSC-A and subsequent gating on DAPI as indicated. Sorts were completed with semi-purity mode, and a resort was always performed to check sorting efficiency.

### MERFISH

FFPE samples were sectioned to 4.5 µm, placed on a functionalized slide (Vizgen, 20400100) and gently pressed onto the slide to enhance tissue adhesion. Subsequently, the slide was dried at room temperature for 60 minutes, then baked at 55°C for 30 minutes and placed at room temperature again for 3 hours before storage at 4°C until use. After equilibrating the slides to room temperature, deparaffinization was performed with deparaffinization buffer (Vizgen, 20300112) in two cycles of 55°C for 5 minutes, followed by a single cycle at room temperature for 5 minutes. The tissue was rehydrated by three washes with 100% ethanol (Fisher Scientific, BP2818500) and one consecutive wash with 90% and 70% ethanol. The slides were dried at room temperature for 30 minutes, then incubated with decrosslinking buffer (Vizgen, 20300115) at 90°C for 15 minutes. Before further processing, FFPE samples were split into MERFISH chemistry V1 and V2. For V1, conditioning buffer (Vizgen, 20300116) was added to the slide for 30 minutes at 37°C. For pre-RNA anchoring, 5 µl pre-anchoring activator (Vizgen, 20300113) and 5 µl RNase inhibitor (Roche Diagnostics, 09537643103) were diluted in 100 µl conditioning buffer and incubated on the tissue for 2 hours. Multiplex protein staining steps were performed before adding formamide wash buffer (Vizgen, 20300002) at 37°C for 30 min and an overnight incubation in 100 µl RNA anchoring buffer (Vizgen, 20300117). RNA anchoring buffer was removed from the tissue by adding formamide wash buffer at 47°C for 15 min and sample prep wash buffer (Vizgen, 20300001) afterwards. Polyacrylamide gel was prepared by mixing 5 ml gel embedding premix (Vizgen, 20300118) with 25 µl 10% ammonium peroxodisulfate (Karl Roth, 9592.2) and 5 µl N,N,N′,N′-Tetramethylethylendiamin (Sigma-Aldrich, T7024-25ML). The slides were coated in gel solution for 5 min, then the solution was removed and 50 µl gel solution added to the center of the tissue. The gel was evenly spread over the tissue with a coverslip (Vizgen, 30200004) coated with gel-slick solution (Lonza, 50640) and incubated until fully polymerized. Afterwards, the coverslip was carefully removed and the tissue digested overnight at 47°C by adding 5 ml clearing premix (Vizgen, 20300114) with 50 µl Proteinase K (New England Biolabs, P8107S). To reduce tissue autofluorescence, the tissue was bleached with blue LED light for 3 hours (Vizgen, 10100003). After three washes with sample prep wash buffer and one with formamide wash buffer at 37°C for 30 min, 100 µl custom gene probes were added onto the gel and spread evenly by placing a small piece of parafilm on top. Probe hybridization with V1 probes was performed for 48 h at 37°C. To remove unbound gene probes, two washes with formamide wash buffer were performed at 47°C for 30 min each, followed by two washes in sample prep wash buffer. MERFISH chemistry V2 provided an improved RNA detection rate and overall increased sensitivity by optimizing anchoring mechanisms, gene probes and augmented readout-probes. For this version, cell staining was performed immediately after decrosslinking, followed by an anchoring pretreatment with a conditioning buffer (Vizgen, 20300116) incubation for 15 min and a pre-anchoring buffer incubation (100 µl conditioning buffer, 5 µl pre-anchoring activator (Vizgen, 20300113), 5 µl RNase inhibitor) overnight. RNA anchoring was performed the following day for 2 hours at 37°C. The samples were then embedded into gel and cleared with tissue clearing solution at 47°C overnight. After autofluorescence quenching and probe hybridization with V2 probes, an additional enhancer hybridization step was added after the post-hybridization wash with formamide buffer. For this, pre-warmed enhancer (Vizgen, 30300491) was added onto the embedded sample and incubated at 37°C overnight, followed by two additional washing steps with enhancer wash buffer (Vizgen 20300192). All slides were consecutively stained with DAPI and PolyT staining reagent (Vizgen, 20300021) in the dark for 15 min, then washed again with formamide wash buffer and sample prep wash buffer before proceeding to imaging.

Fresh frozen samples were cut to 10 µm sections, placed on functionalized slides (Vizgen, 20400001) and equilibrated to -20°C for 5 min to improve tissue adherence. Then, slides were dried at 47°C for 3 min and immediately fixed with pre-warmed 4% PFA (Electron Microscopy Sciences, 15714-S) at 47°C for 30 min. PFA was removed by washing in 1X PBS (Gibco, 11594516) three times, then the tissue was permeabilized in 70% ethanol at 4°C overnight. After a wash with 1X PBS, multiplex protein staining was performed. The slide was then incubated in formamide wash buffer at 37°C for 30 min before adding a 50 µl custom gene panel and placed at 37°C for 48 h for probe hybridization. Excess probes were removed by two incubations in formamide wash buffer at 47°C for 30 min each, before the tissue was embedded in a polyacrylamide gel prepared from gel embedding premix (Vizgen, 20300004) and cleared in 5 ml clearing premix (Vizgen, 20300003) with 50 µl Proteinase K at 47°C overnight. Photobleaching for 3 hours was initiated the next day. Analogous to the FFPE sample preparation, the slides were washed with sample prep wash buffer, before staining with DAPI and PolyT in the dark for 15 min and a final wash in formamide wash buffer and sample prep wash buffer before imaging.

For multiplex protein staining, the slides were blocked with blocking buffer C premix (Vizgen, 20300100) for 60 min before adding the primary antibody mix (Anti-Desmin 1:50 (Novus Biologicals, AF3844), Anti-Myeloperoxidase 1:100 (Abcam, ab300651), Anti-Vimentin 1:100 (Abcam, ab24525), cell boundary primary stain mix 1:100 (Vizgen, 20300010) diluted in blocking buffer C premix) for 60 min. Unbound primary antibodies were washed away with 1X PBS before secondary antibody mix (Anti-goat Aux 6 protein stain 1:100 (Vizgen, 20300103), Anti-rat Aux 7 protein stain 1:100 (Vizgen, 20300104), Anti-chicken Aux 9 protein stain 1:100 (Vizgen, 20300106), cell boundary secondary stain mix 1:33 (Vizgen, 20300011) diluted in blocking buffer C premix) was added onto the slide for 60 min. After further washing with 1X PBS, a second blocking step with diluted rabbit IgG (Thermo Fisher, #02-6102) was performed for 30 min. During this time, a conjugate of Anti-Connexin 43 (Proteintech, 26980-1-AP) and a custom 5’acrydite-Aux 5-conjugated Anti-rabbit nanobody (Massive Photonics) was prepared and diluted in blocking buffer C premix to a final antibody concentration of 1:100. After washing away excess rabbit antibody with 1X PBS, the antibody-nanobody mix was incubated on the tissue for 60 minutes. Rnase inhibitor was added to all steps to ensure RNA preservation. Afterwards, more washes with 1X PBS and a light fixation with 4% PFA for 15 min were performed. The slide was then washed again with 1X PBS and sample prep wash buffer. Imaging Imaging cartridges (Vizgen, 20300019) were activated by adding a mix of 250 µl imaging buffer activator (Vizgen, 20300022) and RNase inhibitor (New England Biolabs, M0314L) to the cartridge port and layering mineral oil (Sigma-Aldrich, M5904-500ML) on top to avoid oxidation. The slides were placed into the flow cell, both imaging cartridge and assembled flow cell were connected to the instrument. After ROI selection and focus acquisition, the measurement was initiated and processing of the raw data performed according to Vizgen’s proprietary processing pipeline.

### Xenium spatial transcriptomics

Spatial in situ sequencing data of human and mouse heart tissues were generated using the Xenium Prime In Situ Gene Expression workflow for FFPE tissue (10X Genomics) with the Xenium Prime 5K Human or Mouse Pan Tissue & Pathways Panel (10X Genomics, 1000724/1000725, RevE). Samples were sectioned to 5 µm and carefully placed in the sample area of Xenium slides following the Tissue Preparation Guide (10X Genomics, CG000578). Slides were dried for 3 h at 42°C, then placed in a desiccator at room temperature overnight. Xenium slides were fixed and permeabilized according to the Xenium In Situ Gene Expression Fixation and Permeabilization protocol (10X Genomics, CG000580, RevE) prior to probe hybridization, ligation, rolling circle amplification and cell segmentation staining according to the manufacturer’s instructions in Xenium In Situ Gene Expression with optional Cell Segmentation Staining (10X Genomics, CG000760, RevA). For human heart samples, a custom add-on probe panel was added to the priming hybridization and probe hybridization step. Slides were imaged on the Xenium Analyzer (10X Genomics, CG000584, RevF) and an H&E staining was performed afterwards following the manufacturer’s recommendations (10X Genomics, CG000613, RevB). A post-Xenium WGA-FITC staining was performed by incubating 1:100 WGA-FITC on the Xenium-slide in 1X PBS for 30 min, followed by washing in 1X PBS for 3x 5min. Imaging was performed using an AxioScan (Zeiss) at a magnification of 20x.

### Bioinformatics Analysis Data Acquisition

Spatial transcriptomics data was acquired with onboard software versions 234c (MERSCOPE) and 4.0.1 (Xenium). Microscopy images were acquired with Zeiss Zen 3.1.9. snRNA data was aligned with Cellranger 7.1.0. Reads were aligned to the human reference genome GRCh38 using the pre-built reference package version 2024-A provided by 10x Genomics. This reference incorporates the GENCODE v44 annotation and the GRCh38 primary assembly sequence. Reads were aligned to the mouse reference genome GRCm39 using the pre-built reference package version 2024-A provided by 10x Genomics. This reference incorporates the GENCODE vM33 annotation and the GRCm39 primary assembly. Single-nucleus RNA-seq data were aligned to the Rat reference genome mRatBN7.2 with annotations from Ensembl Release 110.

### snRNA processing

To preprocess and analyse the snRNA-seq dataset we created a standardised Snakemake^87^ pipeline that includes the following steps: ambient RNA detection and removal with SoupX^88^, quality control and normalisation with Scanpy^89^, doublet detection with scrublet^90^ and doubletdetection^91^, integration with Harmonypy^92^ and differentially expressed marker genes analysis with Presto^93^. Each dataset underwent identical preprocessing, followed by a semi-manual cell type annotation with support from CellTypist^94^. Human, mouse and rat data annotations were harmonized on the cell type level, while cell state annotation remained dataset specific.

Cells were filtered using a multi-step quality control procedure. First, cells expressing fewer than 2 genes were removed. Subsequently, outlier cells were identified using a median absolute deviation (MAD)-based approach across three quality metrics: total counts (log-transformed), number of genes detected (log-transformed), and percentage of counts in the top 20 genes. For each metric, cells deviating more than 5 MADs from the median were flagged as outliers (similar to Germain et al.^95^). The SoupX ambient RNA detection was then applied with default parameters with the exceptions of the iPSC snRNA-seq data due to the extremely homogeneous nature of the cell line data, as suggested by the package authors. For the integration with Harmony, we selected 4000 highly variable genes across samples using a batch-aware approach with the unique sample identifier provided as the batch key. Expression counts were normalised to the total library size, log-transformed, and scaled without zero-centring. Then we performed principal component analysis (PCA) using the default number of 50 principal components (PCs) on the scaled data and used Harmony for the batch correction with the sample label as the integration variable and a maximum of 100 iterations. Nearest-neighbour graphs were constructed using the Harmony-corrected principal components. Cell clustering was performed with the Leiden algorithm at two resolutions 1.5 and 3 for further manual annotation and cluster assignments were stored separately. Then we generated two-dimensional embeddings using PaCMAP and UMAP applied to the Harmony-corrected PCA space. We performed automatic cell type annotation using CellTypist on the integrated dataset, following the authors instructions. Additionally, we calculated cell cycle phase scores for S-phase and G2/M-phase gene expression using predefined gene sets^96^.

We identified marker genes of the integrated datasets for both clustering resolutions using a two-sided Wilcoxon rank-sum test implemented in the Presto framework. Genes were ranked based on the Wilcoxon AUC statistic. Marker genes were filtered to retain the top 100 genes per cluster with an AUC of at least 0.5, expression in all cells of the cluster, and an adjusted p-value below 0.05. For all datasets during the initial manual cell type annotation process we used overclustered resolution (leiden res = 3) to annotate major cell types and removed all small low quality, high in doublets subclusters that had a mixed marker gene signatures. Additionally, during the preprocessing of the Embryo human heart snRNA-seq dataset we first annotated all cell types then subsetted, reclustered and reannotated only CMs to achieve higher annotation accuracy. During this step some low-quality, high in doublets mixed clusters were removed.

### Enrichment analysis and gene expression comparison

All gene expression comparisons (e.g. expression heatmaps) were created from normalized, log1p transformed UMI matrices (see processing details above). Differential gene expression and enrichment analysis was performed on the pseudobulk level with edgeR^97^. Pseudobulks were formed at the donor and cell type level as required by the individual analysis and were based on the sum of the count matrix after ambient RNA removal. Enrichment analysis was performed using the competitive gene set test CAMERA^98^ implemented in edgeR^97^ with default settings. The enrichment (t-statistic) of selected gene sets was calculated with a univariate linear model as implemented in the python version of the decoupler library^99^ based on differential genes calculated by edgeR (quasi-likelihood negative binomial generalized log-linear model). Gene sets were retrieved from msigdb^100^ and PROGENy^101^. Complete enrichment scores are given in **Supplementary Table 2**.

### Gene signature scoring per nucleus

We computed per-nucleus signature scores using Scanpy’s gene scoring function for DES⁺ and CFLAR⁺ CMs. Then we analyse the signature scores in relation to cellular ploidy using the polyploid annotated data. Pairwise comparisons between ploidy groups 2N, 4N, and 8N were performed separately for each signature using the two-sided Wilcoxon rank-sum test.

### Abundance testing

Cell-type abundance differences were analyzed using scCODA, a Bayesian framework for compositional single-cell data^102^. In all analyses, control samples were specified as the reference condition, resulting in direct comparisons between control and disease samples, as well as between untreated and treated iPSC samples. For human and mouse snRNA-seq data, iPSC derived samples and rat samples, credible changes in cell-type abundances were identified using an estimated false discovery rate (FDR) of 0.1. Changes in cell-type abundances were quantified as scCODA-estimated log2-fold changes relative to control samples and visualized using heatmaps with markers indicating credibly changing cell types.

### Comparison of cardiomyocyte substates across data sets

We use two complementary approaches to contrast cardiomyocyte sub-states/clusters across novel (human adult, human embryo, mouse, rat) and public data sets^59^ The first approach, ClusterFoldSimilarity^60^, quantifies the similarity of cell clusters across datasets by their gene fold changes measured against the remaining clusters within the respective dataset and comparing these to the fold change in clusters from a reference dataset^60^. As the resulting similarity score is dependent on the specific context, we refrain from comparing the magnitude of specific values across pairs of considered datasets and only interpret these as indicators of matching or distinct cell types.

As a second approach, we perform a one-sided hypergeometric test to measure the statistical significance of the overlap between cell markers for each cluster^103^. Based on the hypergeometric distribution, the test calculates the probability of observing at least as many overlapping genes by chance when drawing two random sets from the shared background of genes expressed in at least 5% of CMs in the compared datasets. To perform this test, the differentially expressed genes of the subtypes are ranked within each considered dataset (tie-corrected two-sided Wilcoxon test). The resulting top 500 differentially expressed genes of each CM subtype are then compared pairwise across all datasets to determine the number of matching genes. The match of top ranked genes between two subtypes is considered as significant, if the random chance of obtaining at least as many matching genes under the hypergeometric distribution is less than alpha=0.01. As we are only interested in contrasting CM populations, all marker genes and fold similarities described above are based on contrasting only CM cells.

### Spatial transcriptomics panel design

An initial MERFISH V1 500 gene panel was designed based on publicly available scRNA datasets of the human heart and human heart disease^11,104–108^ in addition to canonical marker genes. Cell type annotation was harmonized between public datasets. We then selected the most informative genes using the Spapros^1^ probe selection pipeline. The selection procedure was carried out in three steps. Spapros was first run jointly on all datasets, selecting informative genes based differential expressed (DE) gene statistics. Afterwards, the tool was run on the individual datasets to select informative genes for cell states based on PCA loadings. The performance of the panel was confirmed by using Spapros’ evaluation mode which trains a random forest classifier to distinguish cell types in scRNA data based on subsets of genes. In cases where cell types were commonly confused by the classifier (i.e. the correct cell type was selected less than 60% of the time) additional genes were selected with NSForest^2^, a marker gene selection tool based on a random forest classifier. To reduce the chance of optical crowding the panel adhered to an overall FPKM (Fragments Per Kilobase Million) limit of 9000 in the manufacturers reference RNAseq dataset. Genes with very high FPKM were replaced with genes that showed similar expression patterns but lower FPKM values. This replacement was based on Pearson correlation values.

We generated a second panel for MERFISH (v2) by including genes related to newly discovered CM states, which were not previously defined in the previous analysis. For the MERFISH v2 dataset the procedure was as follows: The snRNA dataset was subset to only cardiomyocytes. Cells were then assigned to the major clustered CFLAR+ and DES+ and marker genes were selected with the tie-corrected two-sided Wilcoxon rank sum test, as implemented in Scanpy^89^. The bottom 15 genes in the initial panel, as ranked by importance according to Spapros, were replaced with CM state marker genes. For the MERFISH v2 panel we added an additional step to further decrease the chance of optical crowding occurring. The final MERFISH v2 codebook, which assigns binary barcodes to transcripts^109^, was shuffled in a way where highly expressed genes in the same cell type were not hybridized with fluorescent probes during the same detection round. This optimization procedure^110^ relied on simulated annealing and used our human snRNA dataset as input. All MERFISH panels were purchased from Vizgen Inc.

For the Xenium experiments (10X Genomics) we used the preconfigured 5000 pan tissue panel in the mouse and human variant (PO 1000725 and 1000724). A custom 100 gene human addon panel optimized for cardiomyocyte state differentiation. The procedure was the same as for MERFISH, but only considered 100 top genes. This panel was added to all measurements of human samples. A detailed description of all panels can be found in **Supplementary Table 3**.

### Segmentation of spatial transcriptomic datasets

For MERFISH, the segmentation of heart muscle cells was extensively optimized. We fine-tuned a Cellpose 3 cell membrane model, “heartbreaker”, on 216 (1000 x 1000 px) expert annotated images from our MERFISH measurements, specifically Vizgen’s “Cell Boundary Stain 3” and the DAPI channel. Images were split into 200 training and 16 test images. The model was initialized with “cyto3” weights and then trained with a learning rate of 0.1 and a weight decay of 1e-05 for 200 epochs. In a benchmark we compared the performance of our model to several image and transcript-based segmentation algorithms. For image-based we considered the default MERSCOPE segmentation based on Cellpose 2, Cellpose 3 “Nuclei” model and our model. We additionally tested a configuration where the result of heartbreaker was combined with the output of the “Nuclei” Cellpose 3 model.

All image-based tools were run on boundary stain 3 from the Vizgen MERSCOPE cell boundary stain kit. Transcript-based segmentation methods tested included Baysor^111^, BIDCell^112^ and Proseg^113^. Performance was judged by comparison to a total of 10 hand-annotated image sections (1500 x 1500 px) from two MERFISH datasets. Tools were scored based on how similar their output was to the expert annotation. For this purpose the aggregated jaccard index (AJI)^114^ and mean average precision at intersection over union of at least 50% (mAP IoU 50%) were used. We further ranked tools on the percentage of all transcripts assigned to cells and based on the number of segmented cells in the two tested MERFISH samples. As the total true number of cells in a sample is unknown, we compared the number of output cells to the number retrieved from nuclei segmentation. The absolute value of the difference between the estimated total number and number output by a tool was min-max scaled between the tools.

Heartbreaker achieved the overall best performance, combination with a second round of nuclei segmentation did not improve results (**Extended Data Fig. 2b**). Cellulose segmentation of the cell boundary stain closely adhered to human annotation, while Baysor, BIDCell and Proseg tended to oversegment large heart muscle cells.The default MERSCOPE segmentation mostly segmented nuclei and split cardiomyocytes into a large amount of small round cells. Nuclei segmentation with Cellpose reliably segmented nuclei (**Extended Data Fig. 2c**), but did not assign a large number of transcripts (**Extended Data Fig. 2b**). Based on these results all MERFISH datasets were segmented with heartbreaker.

For Xenium, datasets were run with the “Multimodal Cell Segmentation” kit, based on ATP1A1, E-Cadherin and CD45 antibodies for membrane staining and recommendation in CG000750. Compared to the MERSCOPE, the membrane stain was less well-suited for cardiomyocyte segmentation and suffered from lower contrast between foreground and background (**Extended Data Fig. 2d**). To minimize segmentation errors we therefore opted for nuclei segmentation with the Cellpose “Nuclei” model for all Xenium data except the TMA used in the subcellular analysis experiment (**Supplementary Table 3**).

Subcellular analysis (**Fig. 2f-k**) required us to precisely segment cells and their nuclei in our Xenium data. Here, we employed a new approach. A single TMA with 26 samples was treated with a post-run WGA and DAPI stain, as described above. This resulted in sharper boundaries with less background compared to the 10x segmentation kit (**Extended Data Fig. 2e**). The post-run stain was aligned with the images acquired during the Xenium measurements by affine transformation. Matching landmarks from both DAPI channels were picked in the Xenium explorer software. The WGA staining was then run through a white top hat filter^115^ and segmented by morphological watershed in Napari^116^.

### Spatial transcriptomic processing

All datasets were processed in a standardized fashion with Sopa^8^ which performed segmentation, assignment of transcripts, calculations of protein intensity and conversion to the Spatialdata^9^ format. Cells that contained less than 20 transcripts (50 for human Xenium samples) or expressed less than 3 unique genes were not considered in further analysis tasks. Samples where a majority of cells were filtered this way were not considered for further analysis. The data was then processed in a similar fashion as described above for the snRNA data. For the Xenium data 2000 variable genes were calculated, while this step was skipped for the MERFISH data due to the limited panel size.

Cell type annotation was performed by label transfer from the snRNA dataset of the corresponding species. Detailed annotation on the cell state level was used for this transfer (**Supplementary Table 2**). Annotation was performed with Insitutype, TACCO^40^, SingleR, Phi-Space^42^ and RCTD^39^. For every cell majority voting was performed and the number of tools in agreement was calculated. These values were used to calculate the average number of method agreements per cell state. For cell states where this number was below three, the cell state’s annotation was merged with the alternative states proposed by the dissenting tools. This resulted in datasets with annotation that was as detailed as possible, while still being reproducible by the different annotation tools. As an additional quality control step we calculated Spearman’s correlation and Jensen–Shannon divergence between spatial transcriptomic samples and snRNA samples from the same patients. This showed strong correspondence between cell type proportions (Spearman’s ρ > 0.8). The cell type annotation per dataset is further described in **Supplementary Table 3.**

### HeartNet patch based histology analysis

We jointly analyzed the histology of our human and mouse samples. This included WSIs from 53 human and 15 mouse Xenium samples, as well as 23 MERFISH samples. We obtained a patch level embedding (1536 dimensions) by extracting 1,135,625 non-overlapping 224 × 224 pixel (38 µm × 38 µm) patches and running them through the UNI foundation model^28^. Next we assigned expert annotations to the patches. We applied PCA, (50 components) to the embedding followed by Harmony integration^92^. We then performed Leiden clustering^29^ with the resolution 0.3 on the integrated embeddings, which revealed nine distinct patch clusters. These clusters were subsequently annotated by expert assessment of representative examples for every cluster.

To enable scalable annotation of new incoming samples, we developed HeartNet, a deep learning classifier trained on 1,135,625 patch embeddings with expert-annotated clusters from our integrated multi-platform dataset. HeartNet is implemented as a fully connected deep neural network operating on patch-level embeddings. The architecture consists of three hidden layers with 256, 128, and 64 neurons, respectively, each followed by batch normalization, ReLU activation, and dropout (rate = 0.3) to improve training stability and reduce overfitting. The network is trained using the Adam optimizer with weight decay (1e-4) and cross-entropy loss. Early stopping based on validation accuracy and adaptive learning-rate scheduling are employed to ensure robust generalization. Using a train/validation/test split (70%/10%/20%) on balanced classes, the model achieved 93.4% accuracy on independent test data, outperforming traditional machine learning approaches (Logistic Regression: 86.9%, Random Forest: 90.4%). We then applied the trained model to annotate 526,242 patches across 9 MERFISH samples using consistent feature extraction pipelines. In total, our comprehensive analysis encompassed 100 samples and 1,661,867 patches across all platforms.

To relate patches with cellular and nuclear information of spatial transcriptomic experiments, we performed a spatial alignment between transcriptomic data and H&E histology. Two different strategies were deployed, based on the difficulty of the task. For the human Xenium samples we used affine transformation as implemented in the Xenium explorer. Mouse Xenium samples showed higher levels of distortion after H&E staining, while MERFISH alignment was difficult because the H&E staining was performed on a subsequent tissue slice. Here we applied a more advanced alignment strategy. We picked matching landmarks in the BigWarp^117^ Fiji^118^ plugin. We fit thin plate splines^119^ to the landmarks and transformed the centers of cells or nuclei from spatial transcriptomic coordinates to the H&E image space. For MERFISH this was only possible for 9 samples. The overall average number of nuclei per patch for mouse, human Xenium data and Merfish was 5.16.

### NicheSphere based Differential Colocalization

We used NicheSphere to detect condition specific niches in Xenium experiments (https://github.com/CostaLab/Nichesphere). For this, NicheSphere first calculates pair-wise colocalization probabilities for different cell type pairs by considering the k-nearest neighbors in space (k=5) for each sample. Afterwards, we compared the distribution of colocalization probabilities for each cell type pair in two contrasts (Controls vs. ICM,Controls vs. DCM, Controls vs. AMI) via two-sided Wilcoxon tests^120^. Only significant edges are kept in the colocalization graph and the signed weights are set to the W statistic. Then we computed an adjacency matrix of the graph by measuring an Euclidean distance on the weights and used this as input for Louvain graph clustering^121^. Clusters indicate spatial niches. Finally, we use the niche information to perform a graph embedding for visualization purposes using the community layout (https://github.com/alexodavies/CommunityLayout).

### Patient-level analysis

Next, we used NicheSphere niches for sample level trajectory analysis using PILOT^43^. Due to the use of the disease contrasts to define the niches, PILOT provides a semi-supervised trajectory sorting samples according to how well they have niches across the two contrasts, i.e. healthier samples have a lower pseudo-time and more diseased samples have higher pseudo-time and milder cases are in between. PILOT uses optimal transport to calculate distances between pairs of samples based on cell type proportions per sample. Then, a sample trajectory is inferred based on those distances. We modified PILOT, which originally works on transporting cell clusters distributions, using cells similarly as transport cost. Here, PILOT uses the niche sizes as distributions and the cosine distance (PILOT’s default option) on the proportion matrix is used as cost function.

Next, we used linear and non-linear regression models in PILOT to characterize niche proportion changes and gene expression changes associated with the disease progression. In other words, for each feature in gene space and niche abundances, we fitted regression models of increasing complexity, including linear, quadratic, and combined linear–quadratic forms. Model significance was evaluated using the F-statistic, with corresponding p-values calculated to test the null hypothesis that all regression coefficients are simultaneously zero. Features whose best-fitting model produced a p-value exceeding 0.05 were deemed statistically non-significant and subsequently excluded from further analysis. Gene selection was performed in a cell type specific way. Moreover, for gene features, we use a Wald Test to contrast the cell type specific patterns with the pattern from other cells. Due to the panel characteristic of the Xenium data, we used a lenient threshold (Wald statistics > 1.8) and a fold change larger than 0.5.

### Subcellular pattern identification

After nuclei and cells were segmented we applied a strict preprocessing to the Xenium TMA measured for subcellular analysis. All nuclei with less than 10 transcript counts, an isoperimetric quotient (measure of circularity) of less than 4 and an average DAPI intensity of less than 200 were removed. After label transfer from snRNA, as described above, we additionally removed all cells and nuclei which were not labeled as cardiomyocytes. All cells which were smaller than twice the 95th percentile of nuclei area were removed. In addition all cells with an area larger than 300,000 square pixels were removed. We then removed all nuclei not intersecting cells and all cells intersecting more than 2 nuclei and cells with less than 50 transcript counts.

We then performed the subcellular analysis with the Bento^46^ package. A raster grid with a square length of 10 pixels was laid over the segmented cells. We then computed a square-wise vector defined as the difference between the local neighborhood gene composition and the total cell composition. These vectors were normalized and projected into a lower-dimensional space using truncated singular value decomposition as a measure of intracellular spatial variance. In the next step we clustered this embedding using Self-Organizing Maps. The optimal number of clusters was determined dynamically by evaluating the quantization error across a range of 2 to 14 clusters and selecting the inflection point (elbow method). Only squares with at least 100 transcripts were considered for the clustering. This resulted in 4 final domains being assigned to the data.

To interpret these clusters we relied on public APEXseq data^47^ which assigned RNA molecules likelihoods for occurring in different cellular compartments. 500,000 squares were randomly selected. We then performed enrichment analysis using the weighted aggregate method implemented in decoupler^99^. We labeled the domains as nuclear, perinuclear, cytoplasm and intercalated disc based on the enrichment scores and overlay with nuclear (DAPI) and cell membrane (WGA) stainings.

### Nuclear morphology analysis

We analyzed nuclear morphology based on DAPI staining on the strictly preprocessed TMA described above. 167 classical morphological features related to DAPI intensity and cell shape were calculated. Calculations were performed with squidpy^122^ and cp_measure^123^ which implement features offered by Cellprofiler^124^ and scikit image^119^. The raw feature by nucleus matrix was then post-processed. We used Pycytominer^125^ to perform robust median absolute deviation transformation and selected features based on variance and correlation thresholds (default values). This resulted in 154 features being retained. We then formed a pseudo-bulk dataset for every cardiomyocyte state based on the mean of every feature for that cell state and calculated PCA^89^.

Morphological features were compared between DES⁺ CM and CFLAR⁺ CM populations. Numeric features were min-max scaled to the unit interval [0,1]. Differential analysis was performed using Ordinary Least Squares regression with sample included as a fixed effect. To ensure valid inference in the presence of intra-sample correlation, heteroscedasticity-consistent covariance matrices were estimated using the sandwich estimator clustered by sample (R packages sandwich^126^ and lmtest^127^). The derived t-statistics were used to compute p-values, which were subsequently adjusted using the Benjamini-Hochberg procedure (FDR). Effect sizes were derived from the regression coefficients and pooled standard deviations to generate Cohen’s *d*. Shown are only selected features based on variance and Cohen’s d, all features can be found in **(Supplementary Table 3).**

In addition we trained classifiers to predict DES⁺ CM /CFLAR⁺ CM status from the morphology of a nucleus. We compared classification performance based on the morphological features described above to the embedding generated by the self-supervised transformer Dinov3^128^. Binary classification of nuclei morphology was performed using a Logistic Regression classifier with L1 regularization (Lasso). The regularization parameter (*C*) was set to 0.1 to encourage model sparsity and generalization. To address potential data imbalance, class weights were automatically adjusted inversely proportional to class frequencies. Model performance was assessed using stratified 5-fold cross-validation, and results were reported using mean, precision, recall, and F1-scores. Classification of nuclei based on the Dinov3 embedding was more reliable (higher recall, precision and F1 score).

### Cell-cell communication

To investigate changes in cell-cell communication across conditions, we integrated NicheSphere-defined niches with cell-cell communication inference tools. In short, we used scSeqComm to estimate both ligand-receptor (LR) interactions and intracellular activity downstream of these LR interactions. Finally, LR interactions between two conditions were given as input for CrossTalker for a differential cell-cell communication analysis.

In detail, we first extracted gene expression matrices from the snRNA-seq data for each experimental group (controls, ICM, ICM_AMI and DCM). We then applied scSeqComm tool^69^ using default parameters to infer LR interaction strengths independently for each experimental group, applying the ConnectomeDB2020^129^ human and CellTalkDB^130^ mouse LR database. Subsequently, we applied CrossTalkeR^70^ to identify changes in cell–cell communication by comparing the predicted LR interactions between each disease condition and the control (i.e. ICM vs control, ICM_AMI vs control, DCM vs control). The interactions considered by CrossTalkeR were filtered from the scSeqComm output using a threshold of LR strength S_inter > 0.5 and restricted to pairs of cell types identified as significantly colocalized based on NicheSphere analysis for the corresponding spatial contrast.

To estimate differences in intracellular signaling activity between each experimental group and control samples for each signaling pathway, we applied the scSeqComm differential intracellular scoring scheme^131^, leveraging the KEGG pathway resources^132^ with pathway integrations from STRINGDB^133^ (for WNT signaling) and TTRUSTv2^134^, RegNetwork^135^, and HTRIdb^136^ databases for transcriptional regulatory networks. The method quantifies differential intracellular activity based on evidence of altered transcriptional regulation of target genes, assessed using a two-sided Wilcoxon rank-sum test, within the same cell-type in disease versus control conditions downstream of receptor-mediated interactions.

### Multi-ome Integration, Trajectory Analysis and Gene regulatory network

We employed an algorithm based on Fused Unbalanced Gromov-Wasserstein (FUGW)^137^ to integrate cardiomyocytes from the human single nucleus RNA-seq and cardiomyocytes from a previous single nucleus ATAC-seq data^11^, which builds upon SCOT framework^138^. First,we applied dimensionality reduction for both modalities. For RNA, Principal Component Analysis (PCA) was performed, retaining Principal Components (PCs) with an explained variance ratio exceeding 10^-3^. For ATAC, we utilized scOpen^139^ with the dimension set to 100. The intra-modal cost matrices were calculated as Euclidean distance matrices derived from the RNA PCA embedding and the ATAC scOpen embedding, respectively. To construct the inter-modal cost matrix, we first identified common genes shared between the RNA gene expression matrix and the ATAC gene activity matrix. After normalization, a PCA model was fitted on the RNA data restricted to these common genes. The ATAC data (common genes) was then projected into this shared space using the trained RNA PCA model to ensure alignment. We used the pairwise Euclidean distances between the two modalities in this shared PCA space as cost function. We assumed uniform marginal distributions for both modalities. The FUGW hyperparameters were set as follows: the weight balancing Wasserstein and Gromov-Wasserstein distances *alpha* was set to 0.1, the unbalanced constraint parameter *rho* to 1.1, and the entropic regularization parameter *epsilon* to 10^-5^. Finally, the resulting optimal transport plan was used to project the ATAC embeddings from the scOpen space into the RNA PCA space. Once aligned, each RNA cell was paired with its nearest ATAC neighbor based on Euclidean distance.

Subsequently, we employed the Phlower^63^ framework to infer differentiation trajectories of single-cell RNA-seq data. Initially, a k-nearest neighbor (k-NN) graph was constructed in the PCA space with K=15 to capture the local manifold geometry. To capture the global reachability and developmental timescales, we implemented a diffusion process on the graph followed by spectral decomposition, yielding 14 diffusion components. Taking the designated “DES+ CM” type cell group as the root, we applied a Hodge Laplacian (HL) decomposition to compute a continuous pseudotime ordering; Subsequently, 10⁴ trajectories were sampled by stochastic traversals of the kNN graph initiated from the root cells. Finally, cells were ordered along the inferred pseudotime and trajectory paths to reconstruct the dynamic expression kinetics of genes and transcription factors (TFs) throughout the differentiation process.

Based on the differentiation trajectories inferred by PHLOWER, we performed transcription factor enrichment analysis for each identified branch. Specifically, cells were first partitioned into branch-specific populations versus the remaining cell pool according to the trajectory topology. Differential expression analysis was then conducted to identify candidate regulatory factors. A TF was considered a potential regulator if it exhibited a significant upregulation in the target branch compared to other branches, defined by a log2 fold-change (log2FC) threshold of 0.13. The statistical significance (t-test) was further evaluated by assessing the expression consistency among intra-branch cells, yielding corresponding p-values. For all branches, TFs with a p-value < 0.05 were aggregated as the final set of candidate regulators for downstream analysis. We used the above method to select branch specific TFs (**Supplementary Table 4**).

To construct the enhancer based gene Regulatory Network (GRN), we utilized scMEGA^61^ to calculate gene-TF correlations. In the resulting network, TF nodes were annotated with pseudotime values derived from PHLOWER. We filtered for significant TFs by intersecting those selected by PHLOWER (p-value < 0.05) and only considering genes with a correlation > 0.3 when contrasting the gene expression and TF activity scores. Finally, edges were filtered to retain only those with a gene-TF correlation > 0.5. To mitigate network density for visualization, we further restricted the network to the top 500 edges per TF based on correlation strength. The final GRN was produced with a Fruchterman-Reingold force-directed network algorithm.

### Pseudo-time analysis of embryonal cardiomyocytes

For the differentiation of embryonal cardiomyocytes, the dataset was subset to “Differentiating LV-CMs”, “Immature LV-CMs” and “Mature LV-CMs”. A trajectory was then calculated with scMEGA^61^ using a diffusion map^140^ based on harmony-corrected PCA. The trajectory was defined in the order from “Immature LV-CMs” - “Differentiating LV-CMs” - “Mature LV-CMs” with archR^141^ and the first three dimensions of the diffusion map were input into the algorithm. ArchR constructs a cellular trajectory using the centroids of user-defined clusters in reduced dimension, aligning individual cells via the nearest cell-to-trajectory distance. We then analyzed gene expression along the trajectory. Pearson correlation was calculated between normalized log gene expression and the pseudo-time assigned to a cell and genes were ranked based on this value. Enrichment analysis was then performed with Camera^98^ in pre-ranked mode as implemented in edgeR^97^. The queried gene sets were gathered from the msigdb^100^ and encompassed the hallmark^100^ collection, KEGG^132^, Reactome^142^, biocarta and GO^143^. Additionally, we added three custom gene sets composed of 100 marker genes each for DES,CFLAR and NPPA cardiomyocytes to the test. The marker genes were calculated using a two-sided tie-corrected Wilcoxon rank-sum test.

### Spatial statistics

The spatial segregation index was calculated on the MERFISH v2 data (n=13) and was calculated with the dixon function^144^ as implemented in the dixon library^145^. Spatial segregation tests every cell against its nearest neighbor and assesses whether a cell of the same type is found more often than would be expected by chance. We specifically analyzed “self-enrichment” (homotypic neighbor relationships) to determine the degree to which niches form contiguous clusters versus scattered distributions. A comparison between the Z-values for the cell types of interest output by the segregation test was then calculated with the two-sided Wilcoxon rank-sum test. For the comparison of niches between ICM and DCM the mean Z score for niches in every sample was compared using a one-sided Wilcoxon rank-sum test (n=57).

### Analysis of public fetal Visium data

The dataset was retrieved from the original publication^74^. We performed deconvolution with our embryo data as reference with Cell2location^146^. We removed mitochondrial genes and selected genes of interest as suggested by the Cell2location authors. The deconvolution was performed with cells per location set to 15, detection alpha set to 20 and otherwise default parameters. We then calculated the mean number of cell types per Visium spot for every heart compartment annotated by the original authors.

### Analysis of rat in vivo measurements

Sirius red staining was quantified with Halo AI 3.6. Statistics on heart function were calculated in GraphPad Prism. A one-way ANOVA followed by Dunnett’s multiple comparisons test was used for all measurements except bodyweight, where a two-way ANOVA followed by Dunnett’s multiple comparisons test was used.

### Statistics and reproducibility

Sample sizes were determined based on sample availability and no statistical method was used to predetermine sample size. Data generation was randomized with respect to disease and control samples. The investigators were not blinded to allocation during experiments and outcome assessment. Criteria for data exclusion, including filtering of nuclei or cells based on size and gene counts, are detailed in the methods section. Microscopic images shown are representative examples and not exhaustive. Statistical tests and exact sample sizes are provided in the figure legends. Correction for multiple hypothesis testing was performed using the Benjamini–Hochberg procedure, unless specified otherwise. For heatmaps, the use of z-scoring or min-max scaling (0–1) is indicated in the figure legends. For boxplots, the median is indicated by the center line, with whiskers extending from the first and third quartiles to the farthest data point within 1.5× of the inter-quartile range. Violin plots represent a kernel density estimate of the underlying distribution; densities were normalized so that all violins within a panel have the same width.

### Reporting summary

Further information on study design is available in the Nature Portfolio Reporting Summary attached to this article.

### Data availability

Processed snRNA data and spatial transcriptomics are available from https://rwth-aachen.sciebo.de/s/c6oay6kCrAdXgM7. Data will be deposited in public repositories upon publication. Human Sequencing data will be deposited to the European Genome-phenome Archive (EGA), rat and mouse sequencing data will be deposited to the European Nucleotide Archive (ENA). Processed anndata files will be shared on Cell x Gene. Spatial transcriptomic datasets will be shared through the BioImage Archive. The data will additionally be presented in an interactive viewer at heartatlas.kuppelab.org.

### Code availability

Analysis code is available from https://rwth-aachen.sciebo.de/s/LSCsL5j7cnrHFrY. and will be made available on Github and Zenodo upon publication. Code and tutorials for NicheSphere are available at https://github.com/Costalab/NicheSphere. A model checkpoint for HeartNet is available at https://github.com/Costalab/HeartNet.

## Acknowledgements

This project has been funded by the German Research Foundation received by C.K. ( CRU344-417911533, CRU5011-445703531, Emmy Noether EN-459969915); received by I.C. (DFG; 3888802535, CRU5011-445703531) by the the German Ministry of Science, Technology and Space (BMFTR) consortia Graphs4Patients for C.K., M. S. and I.C and E:MED Fibromap and CureFib for I.C. and R.K. This work was further supported by grants to CK from the European Research Council (ERC-StG-101040726), the Boehringer Ingelheim Foundation, Else Kroener Fresenius Foundation (EKFS, Physician Scientist Professorship CK) and the Aventis Foundation. R.K. received support from the German Research Foundation (DFG: SFBTRR219 322900939, CRU344 4288578857858, CRU5011 445703531, KR4073/18-1 (561848128) & INST 222/1582-1), the European Research Council (ERC-CoG 101043403 & 101001791, ERC-PoC 101212974 and 10104303), the Leducq Foundation (ImmunoFibHF 20CVD02 and COMET 24CVD02). G.C. was partially supported by Fondazione Ing. Aldo Gini. Views and opinions expressed are those of the authors only and do not necessarily reflect those of the European Union or the European Research Council Executive Agency. Neither the European Union nor the granting authority can be held responsible for them. We thank Dr. Igor Efimov from the Northwestern University, Chicago, IL, USA, for providing fixed human donor heart samples. NM is supported by the Deutsche Forschungsgemeinschaft (German Research Foundation; TRR 219; Project-ID 322900939. We thank the IZKF Aachen Genomics Core facility for sequencing experiments. The Dutch Fetal Biobank received funding from Amsterdam UMC (Innovation Impuls) and the Amsterdam Reproduction and Development research institute.

We thank all supporting centers, physicians, midwives and nurses for the information they have provided to parents who considered donating their fetuses to science and in particular to the DFB. Most importantly, we express our gratitude to all mothers and couples who have donated their most precious gift to the scientific community.

This study is part of the UCC-UNRAVEL biobank (www.unravelrdp.nl). The study was approved by the local institutional ethics review board (University Medical Center Utrecht, protocol UCC-UNRAVEL #12-387 and TCBio 25-242) ERN UNRAVEL is Member of the European Reference Network for rare, low prevalence and complex diseases of the heart: ERN GUARD-Heart (ERN GUARD HEART; http://guardheart.ern-net.eu)

The data used in this publication was managed using the research data management platform Coscine with storage space granted by the Research Data Storage (RDS) of the DFG and Ministry of Culture and Science of the State of North Rhine-Westphalia (DFG: INST222/1261-1 and MKW: 214-4.06.05.08 - 139057).

The authors gratefully acknowledge the computing time provided to them at the NHR Center NHR4CES at RWTH Aachen University (project number p0020567). This is funded by the Federal Ministry of Education and Research, and the state governments participating on the basis of the resolutions of the GWK for national high performance computing at universities (www.nhr-verein.de/unsere-partner).

The authors wish to thank the many other people who assisted in the completion of the manuscript. Among them Louis Kümmerle for helping with spatial transcriptomic panel design, Clarence K. Mah for his advice on BENTO and Beth Cimini for input on cell segmentation and image alignment. In addition we thank Vizgen, Inc. for giving us early access to MERFISH v2 chemistry and technical support by Marc Reudelsterz and 10x Genomics, Inc. for giving us access to FLEX rat probes.

## Author Contributions

C.K. and I.G.C. conceived and designed the study and supervised the project. CK acquired the majority of funding for the study. D.P., E.S., O.G., S.F., S.S., X.L., M.v.S., M.E., T.S., B.K., J.A., K.L., B.S.d.B., J.H.-v.d.V., S.D., and H.M. organized patient tissue collection, biobanking, and patient consent. C.K. generated tissue microarrays. E.S. and O.G. performed spatial transcriptomics and single-nucleus RNA-sequencing experiments. T.L., D.R.M., and S.D. provided mouse myocardial infarction time-course tissues. M.Z., H.Z., F.R., and A.Z. generated and provided rat tissues and snRNA-sequencing datasets. Human iPSC-derived experiments, including differentiation, functional assays, and snRNA-seq, were performed by L.M., Z.F., and D.W., with guidance from A.A. and C.K. P.K. designed the MERFISH and Xenium panels, optimized the image segmentation approach, and performed subcellular spatial transcriptomic analyses. K.K. performed histological analyses; H&E-stained cardiac tissues were evaluated by K.K. and M.E. Computational analyses were structured as follows: M.R. performed niche analyses; G.C. conducted cell–cell communication analyses; M.J. performed PILOT and histology patch analysis. These analyses were advised by I.G.C. and C.K. All other computational analyses—including data preprocessing, quality control, cross-species integration, clustering, differential expression analysis, trajectory inference, multi-omic integration, segmentation optimization, and subcellular transcriptomic analyses—were conducted by P.K., M.J., D.P., G.C., M.R., D.K., K.P., and M.C., with guidance from I.G.C., M.S. and C.K. Data interpretation was performed by P.K., I.G.C., and C.K., with input from all co-authors. P.K., I.G.C., and C.K. wrote the manuscript and organized the figures. N.M., K.K., R.K., I.G.C., H.M., M.S., and C.K. critically revised the manuscript and advised on data interpretation. All authors read and approved the final manuscript.

---

**ED Data Figure 1.**
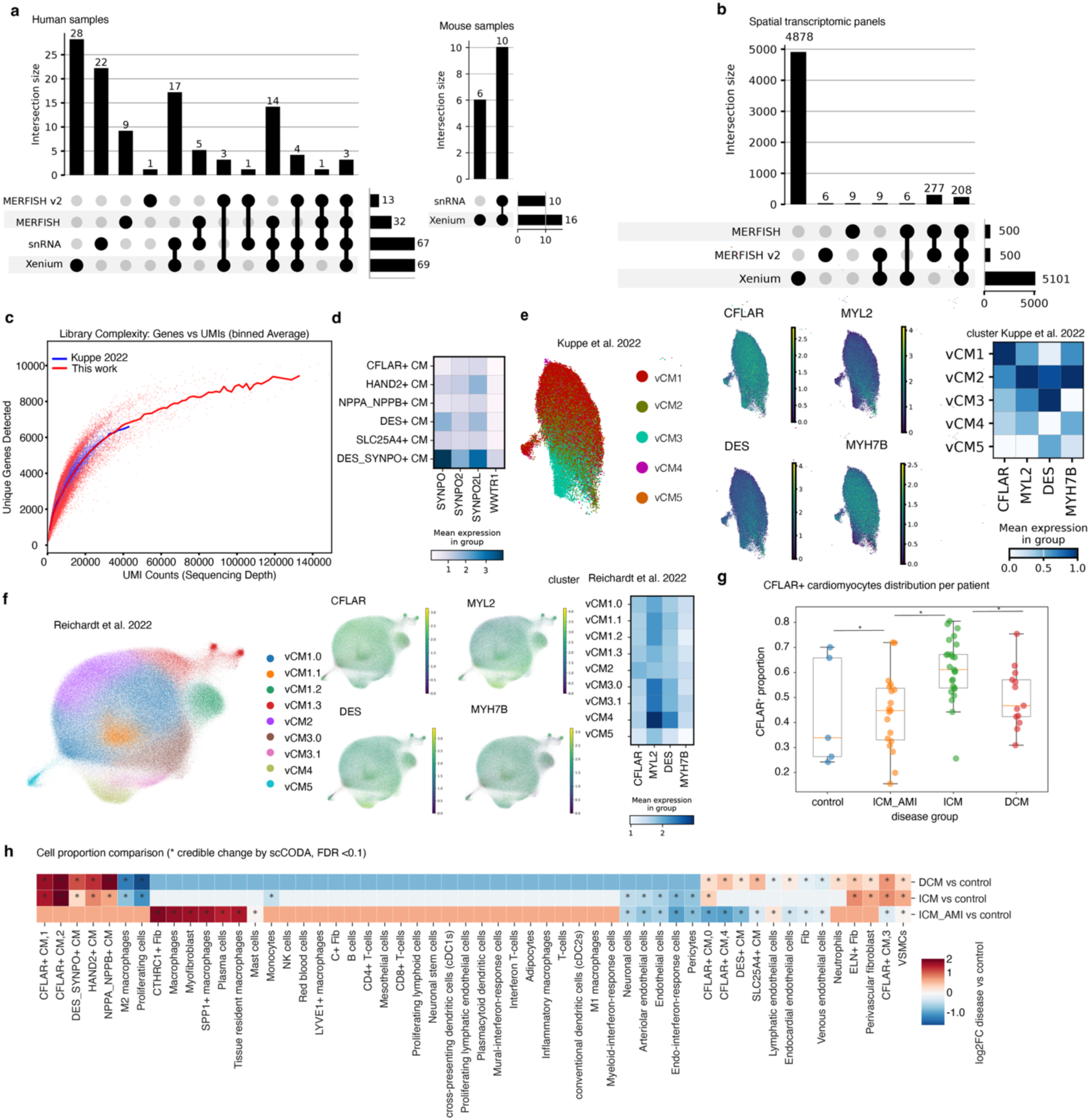
Cross-species identification and segregation analysis of CFLAR⁺ cardiomyocytes across spatial and single-nucleus datasets. **a,** Overview of human (left panel) and mouse (right panel) cardiac datasets used in this study. Bar plots indicate the number of samples per modality, including Xenium, snRNA-seq, MERFISH and MERFISH V2.0. UpSet-style plots summarize dataset overlap and availability across modalities in human and mouse hearts. **b,** Overview of the spatial transcriptomic panels used and their overlap. **c,** Comparison of library complexity (adult human snRNA) with Kuppe et al., 2022. Shown are unique genes per UMI (library complexity). The gene detected by counts per nucleus slope is comparable, while this dataset shows a higher top end. **d,** Heatmaps showing mean expression of SYNPO, SYNPO2, SYNPO2L, and WIF1 across major cardiomyocyte (CM) subtypes in human. DES_SYNPO⁺ CMs display selective enrichment of SYNPO family genes, indicating a distinct cytoskeletal-associated CM program conserved across species. **e,** Projection of cardiomyocyte nuclei from Kuppe et al., 2022 onto a low-dimensional embedding, annotated by CM clusters. Feature plots illustrate expression of CFLAR, MYL2, DES, and MYH7B. The accompanying heatmap summarizes mean gene expression per CM cluster, highlighting segregation of CFLAR⁺ CMs from classical contractile CM states. **f,** Independent validation in the Reichardt et al., 2022 human ventricular cardiomyocyte dataset. Left, UMAP embedding of ventricular CM subclusters, where clusters are name as in the original publication (vCM1–vCM5). Middle, feature plots showing expression of CFLAR, MYL2, DES, and MYH7B. Right, heatmap of scaled mean expression per vCM subtype, confirming selective enrichment of CFLAR and MYH7B in distinct CM populations. **g,** Quantification of the proportion of CFLAR⁺ cardiomyocytes across disease conditions (control, ischemic cardiomyopathy with acute myocardial infarction (ICM_AMI), ischemic cardiomyopathy (ICM), and dilated cardiomyopathy (DCM)). Each dot represents one patient sample, illustrating disease-associated shifts in CM state composition. Stars indicate a FDR<0.1 based on scCODA abundance test. **h,** Differential cell-type proportion analysis across disease groups. Heatmap depicts log-fold changes in relative cell abundance between pairwise disease comparisons. Asterisks indicate credible changes by the scCODA test (False Discovery Rate=0.1). CFLAR⁺ CM populations show selective expansion in cardiomyopathy compared to control hearts, while other CM and non-CM populations display distinct disease-specific remodeling patterns.

**Extended Data Figure 2.**
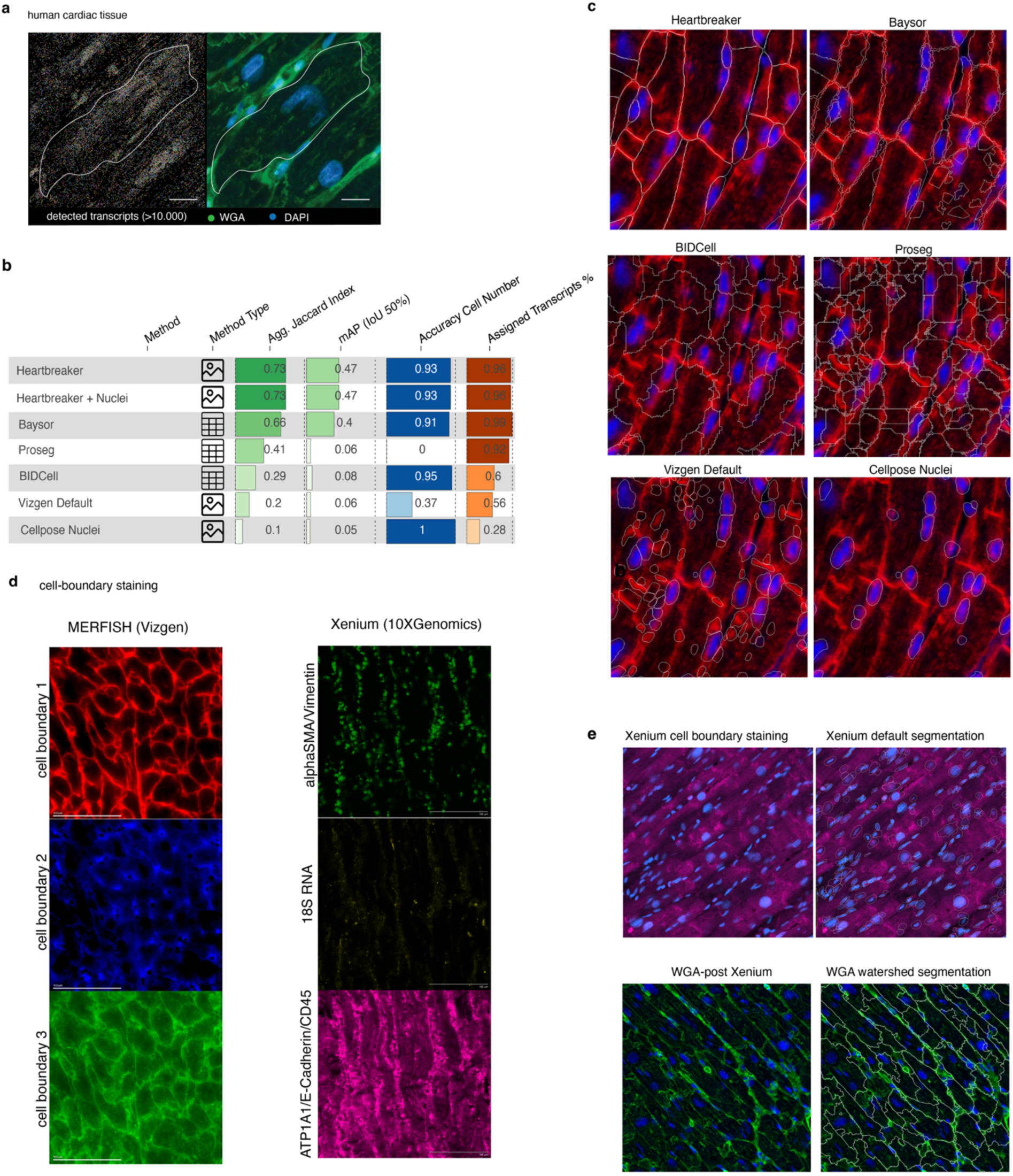
Cardiac tissue segmentation optimization. **a,** Representative human cardiac tissue section profiled by high-resolution spatial transcriptomics Xenium 5k. Left, spatial distribution of detected transcripts (>10,000) within a manually outlined cardiomyocyte. Right, corresponding immunofluorescence image showing wheat germ agglutinin (WGA, cell boundaries; green) and DAPI (nuclei; blue). Scale 10 um. **b,** Quantitative comparison of segmentation methods using image-based and feature-based metrics. Image metrics include aggregated Jaccard index and mean average precision (mAP, IoU ≥50%) relative to reference annotations. Feature metrics include accuracy of cell number estimation (vs nucleus segmentation) and fraction of assigned transcripts. Our heartbreaker model achieves the best overall balance between spatial accuracy and transcript assignment, whereas nucleus-only and default platform segmentations underperform in cardiomyocyte-dense tissue. **c,** Visual comparison of segmentation outputs overlaid on cardiac tissue images (segmented boundaries in white). Methods include Cellpose (membrane), Baysor, BIDCell, Proseg, Vizgen default segmentation, and Cellpose nuclei. Membrane-aware methods more accurately capture elongated cardiomyocyte morphology and reduce over-segmentation. **d,** Comparison of cell-boundary staining strategies for spatial transcriptomics platforms. MERFISH (Vizgen), left section, relies on intrinsic membrane signal, whereas Xenium (10X Genomics), right section, benefits from additional post-hoc membrane labeling to improve boundary definition in cardiomyocytes. **e,** Xenium cardiac tissue segmentation comparison. Top, default Xenium cell boundary staining and segmentation. Bottom, post-Xenium WGA staining followed by watershed segmentation, resulting in improved delineation of elongated cardiomyocytes and more accurate cell boundaries.

**Extended Data Figure 3.**
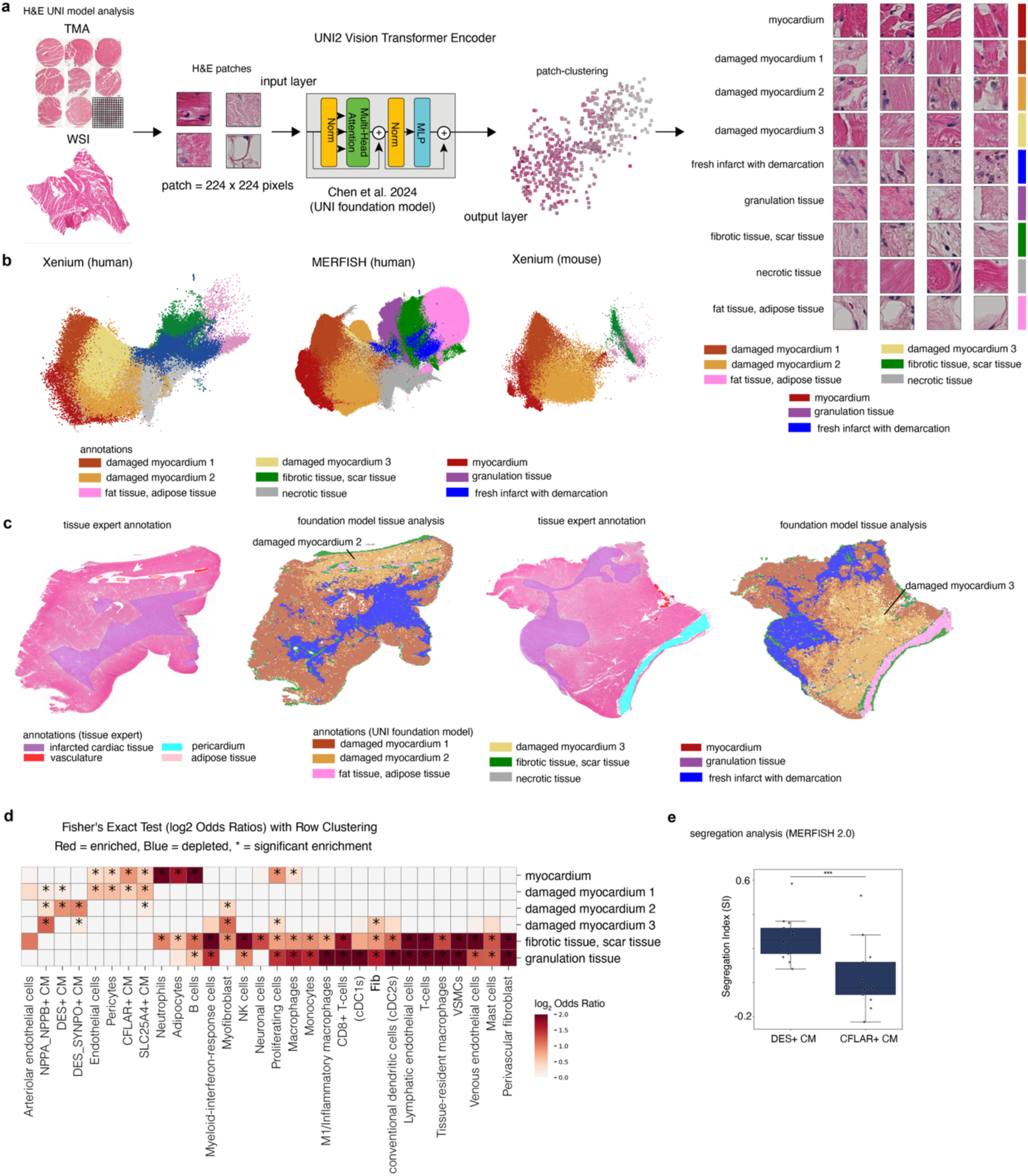
Histopathology foundation model–based tissue state annotation across spatial transcriptomics platforms. **a,** Schematic overview of the histology-based tissue classification workflow using a deep learning foundation model. Whole-slide images (WSI) and tissue microarrays (TMA) stained with haematoxylin and eosin (H&E) were subdivided into non-overlapping image patches (224 × 224 pixels) and embedded using the UNI transformer foundation model (Chen *et al.*, 2024). Latent representations were clustered in an unsupervised manner to define recurrent myocardial tissue states, including intact myocardium, multiple damaged myocardium states, fresh infarct with demarcation, granulation tissue, fibrotic scar tissue, necrotic tissue, and adipose tissue. Representative image patches for each tissue state are shown. **b,** Joint embedding of spatial transcriptomics data annotated by foundation model–derived tissue classes across platforms and species. UMAP projections of Xenium (human), MERFISH (human), and Xenium (mouse) datasets demonstrate consistent segregation of myocardial tissue states, indicating cross-platform and cross-species robustness of the histology-driven tissue classification. **c,** Comparison of expert tissue annotation and foundation model–based tissue analysis on whole heart sections. Left, tissue expert annotations highlighting infarcted cardiac tissue, vasculature, adipose tissue, and pericardium. Right, corresponding foundation model predictions resolving fine-grained myocardial damage states and infarct-associated tissue regions. Arrows indicate regions of concordance and refined subclassification by the model. **d,** Association between tissue states and cellular composition. Heatmap showing enrichment and depletion of cell types within foundation model–defined tissue regions using Fisher’s exact test (log₂ odds ratios). Red indicates enrichment, blue depletion; asterisks denote statistically significant associations after multiple-testing correction (adjusted p-values < 0.05). Fibrotic and granulation tissue regions are enriched for fibroblasts, macrophages, endothelial, and perivascular cells, whereas intact myocardium is selectively associated with cardiomyocyte populations. **e,** Segregation analysis of cardiomyocyte subtypes in MERFISH 2.0 data (n=13). Segregation index (SI) comparison shows significantly stronger spatial segregation (clumping) of DES⁺ cardiomyocytes compared with CFLAR⁺ cardiomyocytes, indicating differential spatial organization relative to tissue damage states. Box plots show median, interquartile range, and individual samples; *** p-value < 0.001 Wilcoxon rank-sum test.

**Extended Data Figure 4.**
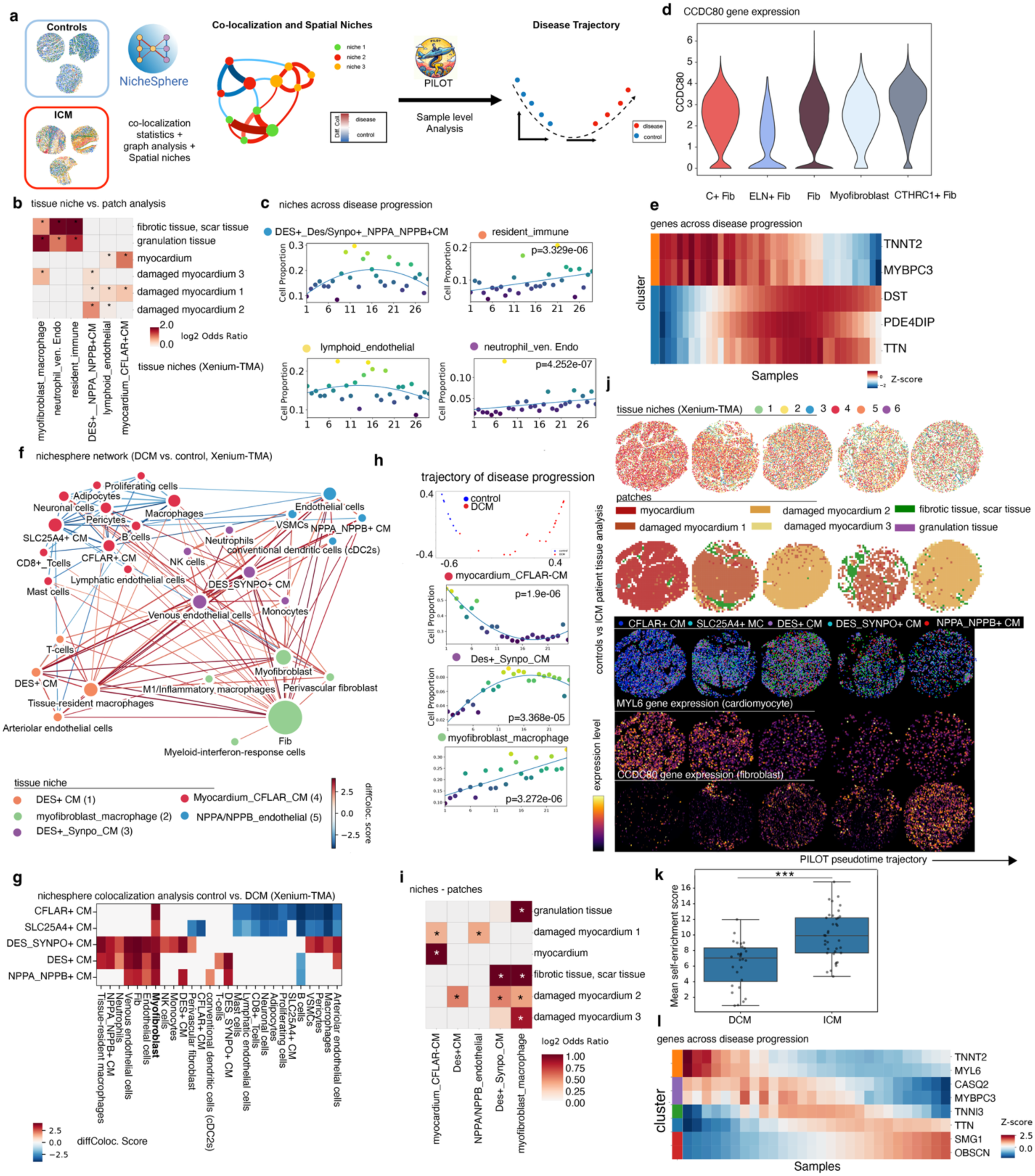
Spatial niche remodeling and disease trajectory in human cardiomyopathy. **a,** Overview of the spatial transcriptomics workflow. Xenium spatial transcriptomic data from control and ischemic cardiomyopathy (ICM) samples were analyzed using the NicheSphere framework to quantify cell–cell colocalization, graph-based interactions, and to define spatial tissue niches. TMA samples were subsequently ordered along a disease trajectory using PILOT pseudotime analysis in accordance to their niches. **b,** Comparison of tissue niche composition versus patch-based histological annotation. Heatmap shows enrichment of defined tissue niches (Xenium-TMA) across histopathological regions, including myocardium, damaged myocardium (subtypes 1–3), fibrotic/scar tissue, and granulation tissue. Asterisks indicate statistical significance (adjusted p-value < 0.05; Fisher’s exact test). **c,** Changes in niche composition (y-axis) across disease progression (x-axis). Scatter plots show proportions of selected niches along the inferred disease trajectory, including DES⁺/DES_SYNPO⁺/NPPA_NPPB⁺ cardiomyocytes, resident immune niches, lymphoid endothelial niches, and neutrophil–venous endothelial niches. Lines indicate smoothed trends; p-values from F statistic test denote significance of association with disease progression. **d,** Violin plots of CCDC80 gene expression (normalized log counts) across major fibroblast-related cell states in snRNA, including C7⁺ fibroblasts, ELN⁺ fibroblasts, general fibroblasts, myofibroblasts, and CTHRC1⁺ fibroblasts. **e,** Heatmap with genes selected (by PILOT) to be associated with the trajectory in CFLAR CM cells. They include cardiomyocyte structural and cytoskeletal genes across disease progression, illustrating coordinated downregulation of sarcomeric genes (e.g., *TNNT2*, *MYBPC3*) and upregulation of remodeling-associated genes (e.g., *DST*, *PDE4DIP*, *TTN*). **f,** NicheSphere interaction network comparing DCM and control samples (Xenium-TMA). Nodes represent cell types or niches, edges represent differential co-localization scores, and edge color indicates increased (red) or decreased (blue) interactions in disease. **g,** Differential co-localization heatmap showing changes in spatial associations between cardiomyocyte states and surrounding cell types in control versus DCM samples. Color scale indicates diffColoc scores. For **f** and **g**, only statistically significant interactions are shown(adjusted Wilcoxon’s rank sums test; p-value < 0.05). **h,** Top, disease trajectory using PILOT pseudotime analysis, Bottom, Proportional changes of selected cardiomyocyte-centered niches along disease progression, including myocardium_CFLAR⁺ cardiomyocytes, DES_SYNPO⁺ cardiomyocytes, and myofibroblast–macrophage niches. Trend lines and p-values indicate significant associations with pseudotime. **i,** Association between spatial niches and histological patches. Heatmap displays the fraction of each niche within defined tissue patches, highlighting preferential localization of cardiomyocyte niches to myocardium and damaged myocardium, and fibroblast–immune niches to fibrotic or granulation tissue. Significance was determined by Fisher’s exact test with adjusted p-values (p < 0.05). **j,** Representative Xenium-TMA sections (five selected samples along the Trajectory from early to middle and late stages) illustrating (top) spatial distribution of tissue niches, (middle) corresponding histopathological patches, and (bottom) cardiomyocyte subtype localization and marker gene expression. MYL6 expression highlights cardiomyocytes, whereas CCDC80 expression marks fibroblast-rich regions. Arrow indicates progression along the PILOT pseudotime trajectory. **k**, Comparison of spatial clustering of niches in ICM and DCM samples. Shown are mean self-enrichment scores (spatial segregation) per sample. (Wilcoxon rank-sum test: Z = 4.09, P < 0.0001) **l,** Heatmap of cardiomyocyte gene expression dynamics across disease progression, showing gradual loss of contractile gene programs (e.g., *TNNT2*, *MYL6*, *CASQ2*, *MYBPC3*) and induction of stress and remodeling-associated genes (e.g., *SMG1*, *OBSCN*).

**Extended Data Figure 5.**
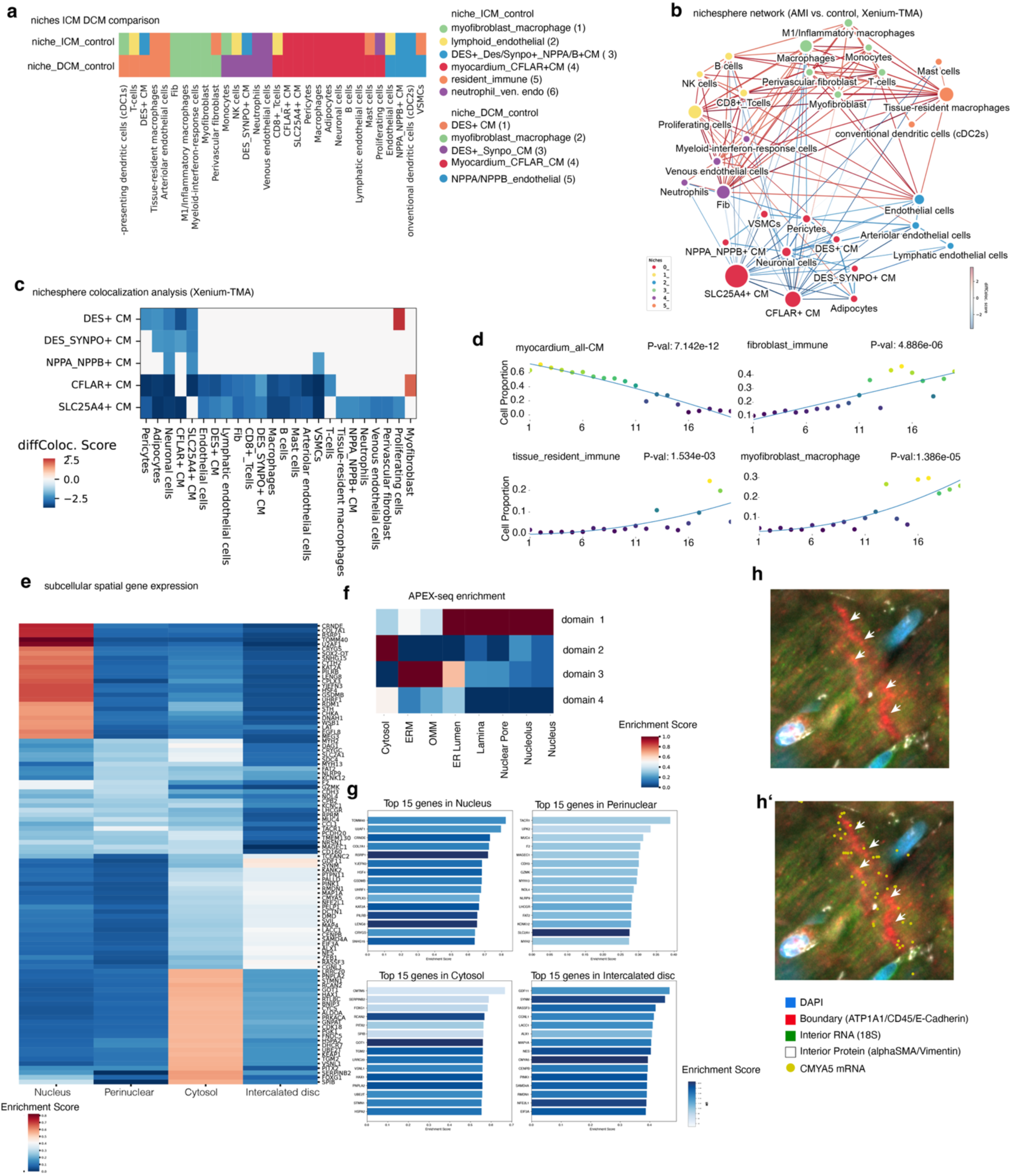
Differential spatial niches and subcellular RNA organization in cardiomyopathy. **a,** Heatmap summarizing differences in spatial niche composition between ischemic cardiomyopathy (ICM)/DCM and control samples. Columns represent annotated cell types, rows indicate identified spatial niches. **b,** NicheSphere interaction network comparing acute myocardial infarction (AMI) with control samples (Xenium-TMA). Nodes represent cell types, edges indicate differential spatial co-localization, with red edges denoting increased and blue edges decreased interactions in disease. Edge thickness corresponds to the magnitude of the differential co-localization score. **c,** Differential co-localization analysis of cardiomyocyte subtypes in Xenium-TMA data. Heatmap displays diffColoc scores for DES⁺, DES_SYNPO⁺, NPPA_NPPB⁺, CFLAR⁺, and SLC25A4⁺ cardiomyocytes with surrounding cell types, highlighting disease-associated remodeling of cardiomyocyte neighborhoods. **d,** Proportional changes of niches along disease progression. **e,** Heatmap of subcellular spatial gene expression patterns in cardiomyocytes. Genes are ordered by preferential enrichment across four subcellular compartments—nucleus, perinuclear region, cytosol, and intercalated disc—revealing compartment-specific transcript localization programs. **f,** Enrichment of APEX-seq–derived subcellular reference signatures across the four identified RNA domains. Heatmap shows concordance of spatial transcript localization with known cytosolic, ER membrane (ERM), outer mitochondrial membrane (OMM), ER lumen, lamina, nuclear pore, nucleolus, and nucleus-associated RNA signatures. **g,** Top 15 genes enriched in each subcellular compartment (nucleus, perinuclear region, cytosol, and intercalated disc), ranked by enrichment score, highlighting distinct functional gene sets underlying compartment-specific RNA localization. **h,** Representative high-resolution microscopy image illustrating cardiomyocyte subcellular architecture. White arrows indicate intercalated disc regions. Nuclei are counterstained with DAPI (blue). **h′,** Same field of view as in **h**, overlaid with detected CMYA5 mRNA molecules (yellow dots), demonstrating preferential localization of CMYA5 transcripts at intercalated disc regions. Boundary markers (ATP1A1/CD45/E-cadherin), interior RNA (18S), and interior protein markers (alphaSMA/Vimentin) are shown as indicated in the legend.

**Extended Data Figure 6.**
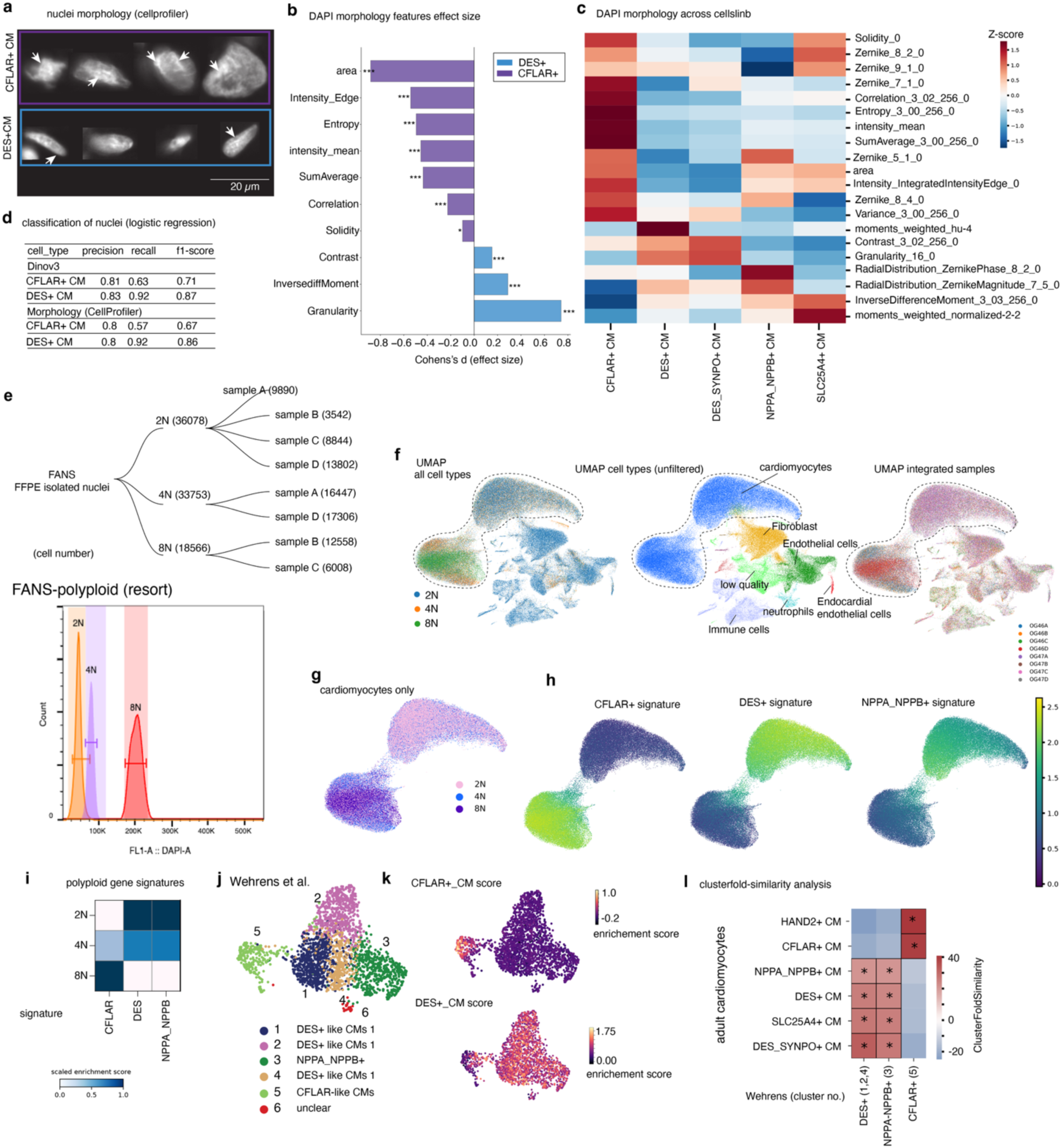
Nuclear morphology–based classification and polyploidy-associated cardiomyocyte programs. **a,** Representative nuclear morphologies. DAPI images of isolated cardiomyocyte nuclei from CFLAR⁺ and DES⁺ CMs illustrating distinct nuclear shapes and texture features (arrows). Scale bar, 20 µm. **b,** Quantitative nuclear morphology features. Comparison of selected CellProfiler-derived DAPI morphology features between DES⁺ and CFLAR⁺ CM nuclei. Bars indicate Cohen’s *d* effect size; positive values indicate enrichment in DES⁺, negative values enrichment in CFLAR⁺ nuclei. p-values from two-sided tests (p-value< 0.05; p-value < 0.001). **c,** Morphology feature heatmap. Heatmap of z-scored DAPI morphology features across cardiomyocyte states (DES⁺, DES_SYNPO⁺, CFLAR⁺, NPPA_NPPB⁺, SLC25A4⁺), highlighting systematic differences in size, texture, shape, and Zernike moments. **d,** Performance of morphology-based classification (logistic regression). Precision, recall, and F1-score for distinguishing CFLAR⁺ and DES⁺ CMs using either DINOV3 feature vectors (top) or CellProfiler based nuclear morphology measurements (bottom). **e,** Polyploid nuclei isolation by FANS. Experimental overview and cell numbers of fluorescence-activated nuclei sorting (FANS) from FFPE samples into 2N, 4N, and 8N fractions, including sample-specific contributions. Bottom panel shows representative DNA content distributions. **f,** Integrated single-nucleus landscape. UMAP embedding of all cardiac cell types (left), annotated cell types with cardiomyocytes highlighted (middle), and integrated samples colored by donor (right). **g,** Cardiomyocyte-only embedding. UMAP representation restricted to cardiomyocytes, capturing continuous transcriptional variation across CM states. **h,** Cell-state signature projection. UMAPs showing enrichment of CFLAR⁺, DES⁺, and NPPA_NPPB⁺ gene signatures across cardiomyocytes. **i,** Polyploid gene signatures. Heatmap summarizing enrichment of CFLAR, DES, and NPPA_NPPB signatures across ploidy classes (2N, 4N, 8N). **j,** External CM classification reference Wehrens et al.. UMAP of cardiomyocytes annotated according to the classification by Wehrens *et al.*, showing correspondence to DES-like, NPPA_NPPB⁺, CFLAR-like, and unclassified CM populations. **k,** Signature score validation. UMAPs displaying CFLAR⁺ CM and DES⁺ CM signature scores, confirming spatial separation of CM programs. **l,** Cluster fold-similarity analysis. Heatmap showing similarity between adult human CM states and reference DES⁺ or CFLAR⁺ programs. Asterisks denote significant enrichment (*P* < 0.05).

**Extended Data Figure 7.**
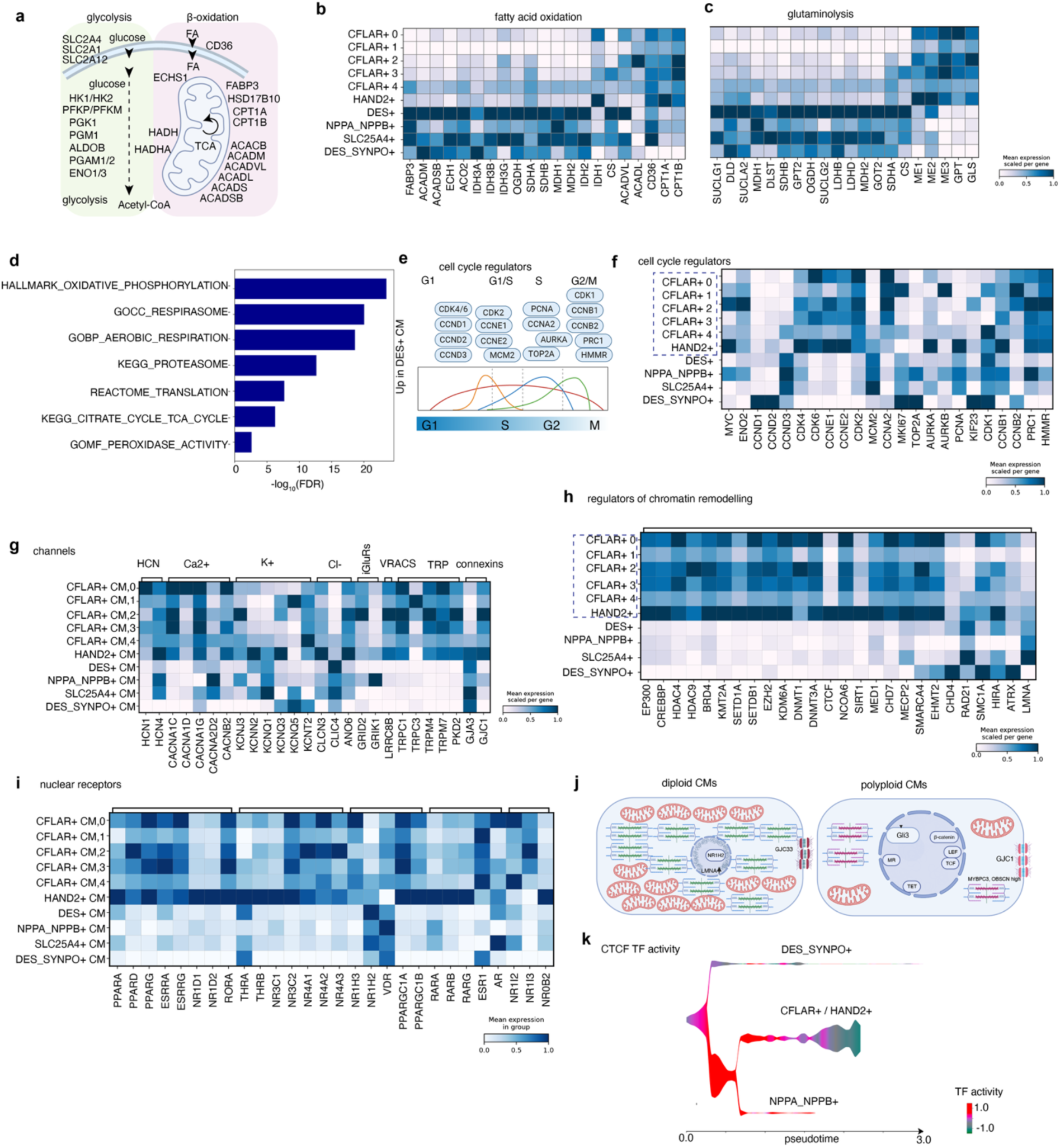
Metabolic, cell-cycle, electrophysiological, and epigenetic programs distinguish diploid and polyploid cardiomyocyte states. **a,** Schematic overview of central metabolic pathways analyzed in cardiomyocytes, highlighting glycolysis, fatty acid uptake and β-oxidation, and mitochondrial acetyl-CoA entry into the tricarboxylic acid (TCA) cycle. Key transporters and enzymes are indicated. **b,** Heatmap showing expression of fatty acid uptake and β-oxidation–related genes across cardiomyocyte states, including CFLAR⁺ subclusters, HAND2⁺, DES⁺, NPPA_NPPB⁺, SLC25A4⁺, and DES_SYNPO⁺ cardiomyocytes. Values represent mean expression scaled per gene **c,** Heatmap of glutaminolysis and TCA cycle–associated genes across cardiomyocyte states, illustrating differential utilization of anaplerotic pathways. **d,** Enrichment analysis -log10 (pvalues). Enriched terms are dominated by mitochondrial respiration, oxidative phosphorylation, and ATP synthesis pathways. **e,** Conceptual schematic of core cell-cycle regulators and their activity across G1, S, G2, and M phases, including MYC, cyclins, CDKs, and mitotic regulators. **f,** Heatmap of cell-cycle regulator expression across cardiomyocyte states, revealing selective enrichment of G1/S and G2/M regulators in specific CFLAR⁺ subclusters relative to other cardiomyocyte populations. **g,** Heatmap of ion channels, transporters, connexins, and receptors relevant for cardiomyocyte electrophysiology and intercellular coupling. Channels are grouped by functional class (e.g., HCN, Ca²⁺, K⁺, Cl⁻ channels, connexins). Mean expression is scaled per gene. **h,** Heatmap of genes involved in histone acetylation, histone/DNA methylation, chromatin remodeling, genome architecture, and nuclear stability across cardiomyocyte states, highlighting enhanced epigenetic and chromatin regulatory programs in CFLAR⁺ cardiomyocytes. **i,** Heatmap summarizing higher-order functional programs across cardiomyocyte states, including metabolic maturation and mitochondrial programs, postnatal maturation and hypertrophy, lipid/cholesterol and inflammatory crosstalk, developmental/regenerative signaling axes, and xenobiotic/stress responses. **j,** Conceptual model contrasting diploid and polyploid cardiomyocytes. Diploid cardiomyocytes are characterized by preserved metabolic maturation, mitochondrial organization, and WNT antagonism (e.g., DKK3, SFRP1), whereas polyploid cardiomyocytes display activation of Hippo, WNT, TGFβ, and SHH signaling, altered genome architecture, and stress-adaptive transcriptional programs. **k**, TF Activity of CTCF within the PHLOWER estimated CM trajectory.

**Extended Data Figure 8.**
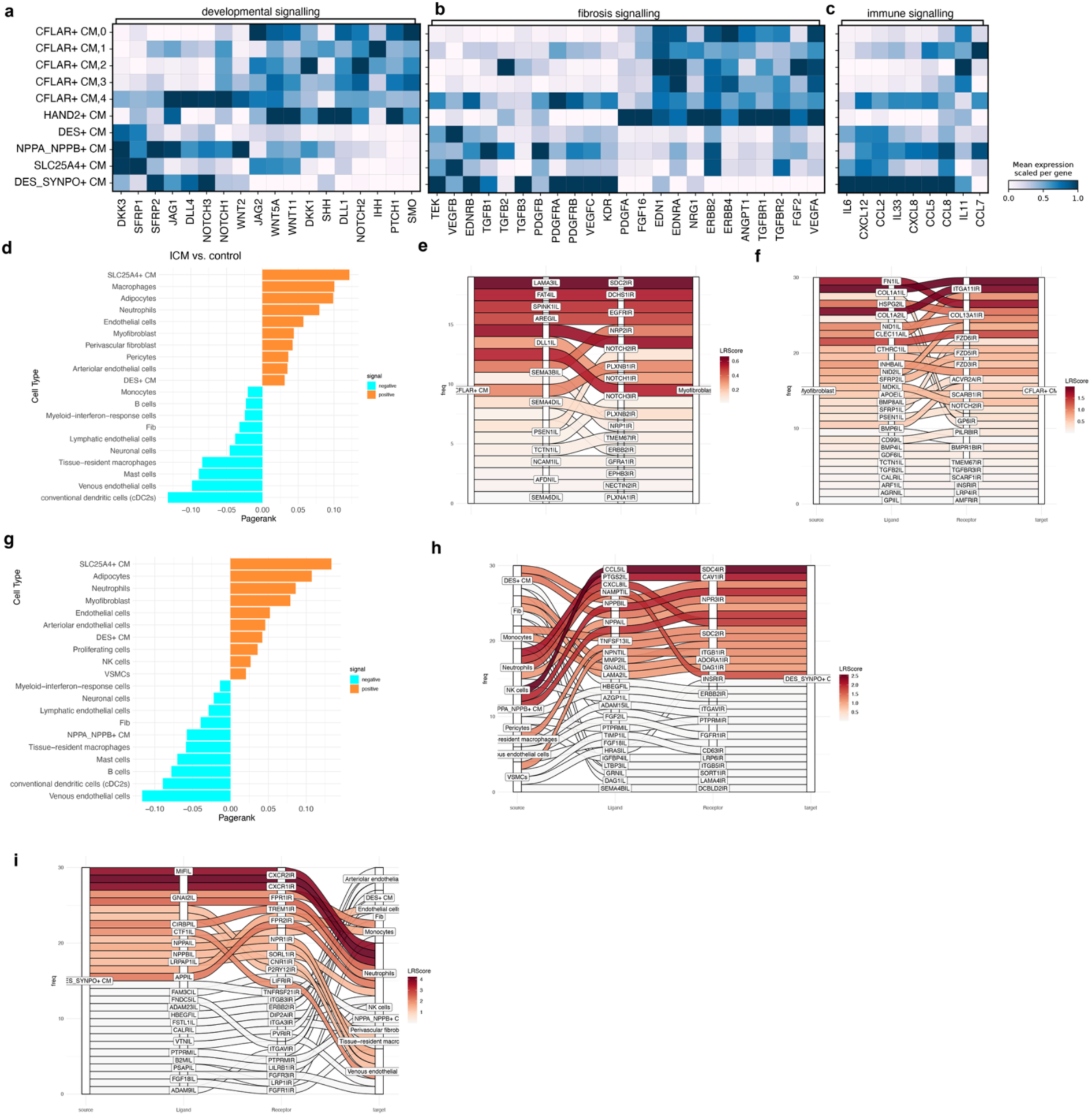
Disease-specific intercellular signaling programs linking cardiomyocyte states to fibrosis and immune remodeling. **a–c,** Heatmaps showing mean scaled expression of pathway-associated genes across cardiomyocyte states. **a,** Developmental signaling pathways, including WNT, SHH, and NOTCH components, display selective activation in CFLAR⁺ and HAND2⁺ cardiomyocytes. **b,** Fibrosis-associated signaling genes (e.g. TGFβ, PDGF, VEGF, FGF pathways) highlight enhanced profibrotic communication potential of specific cardiomyocyte states. **c,** Immune signaling genes, including chemokines and cytokines, indicate differential immune-modulatory capacity across cardiomyocyte populations. **d,** PageRank analysis of cell–cell communication networks in ICM versus control. Positive values indicate higher cell-communication in ICM and negative values indicate lower cell-cell communication in ICM. **e–f,** Sankey plot with top ligand–receptor interaction highlighting ongoing CFLAR⁺-myofibroblast specific communication relative to other cardiomyocyte states in ICM vs control. **e,** Sankey plot with top ligand-receptor interactions between CFLAR⁺ cardiomyocytes towards myofibroblasts, emphasizing fibrosis-related signaling axes. **f,** Reciprocal interactions from myofibroblasts, illustrating cardiomyocyte-driven profibrotic signaling. **g,** PageRank analysis of cell–cell communication networks in DCM versus control. Positive values indicate higher cell-communication in DCM and negative values indicate lower cell-cell communication in DCM. **h,** Sankey diagram with top ligand-receptor interactions with DES_SYNPO⁺ as target cell in DCM versus control. **i,** Sankey diagram with top ligand-receptor interactions with DES_SYNPO⁺ as source cell in DCM versus control.

**Extended Data Fig. 9.**
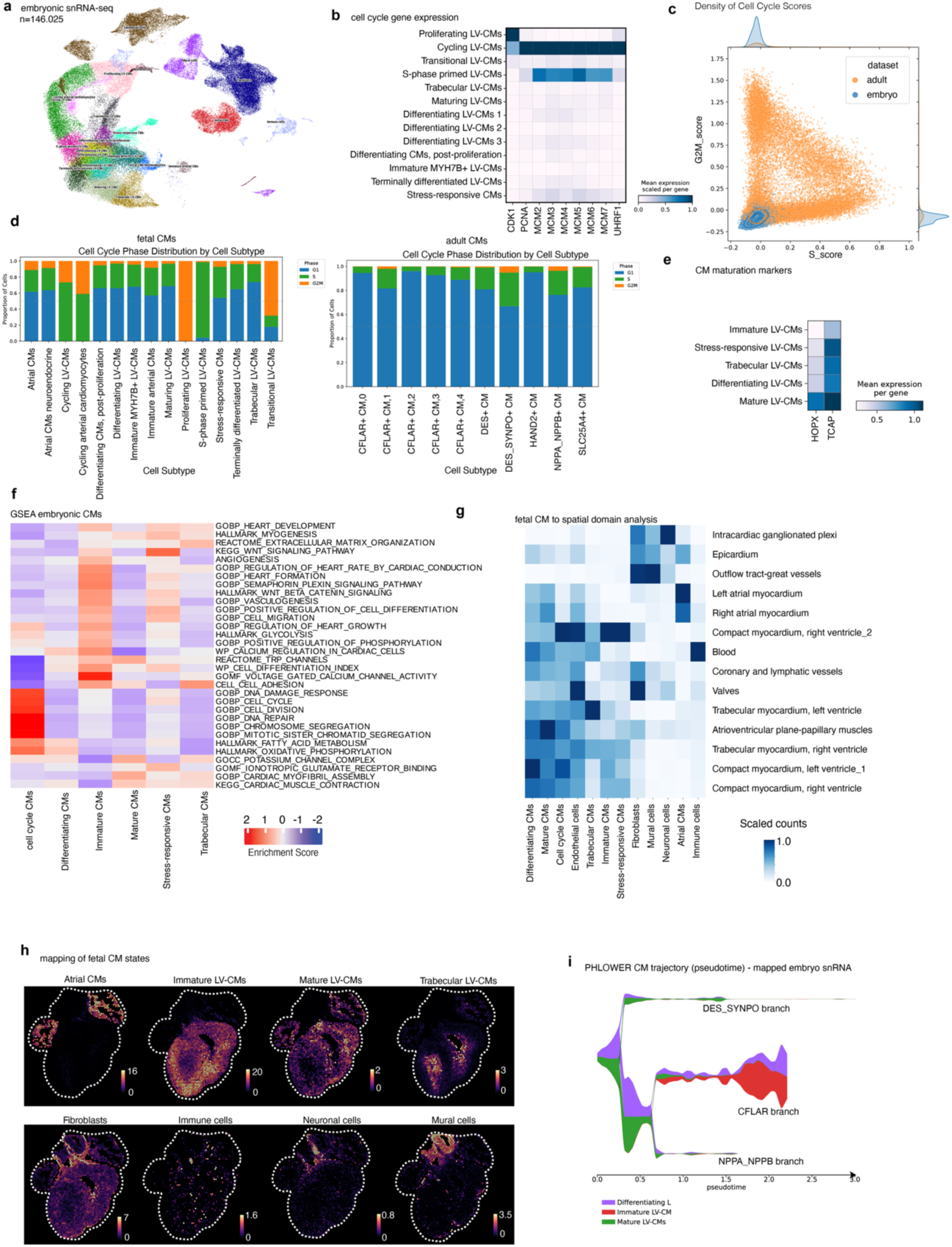
Cell-cycle activity and maturation trajectories of embryonic cardiomyocytes. **a,** UMAP of embryonic human snRNA-seq data highlighting cardiomyocyte differentiation states. **b**, Min-max scaled expression of canonical cell-cycle genes across embryonic LV-CM subtypes. **c**, Density and joint distribution of S- and G2/M-phase enrichment scores in embryonic versus adult cardiomyocytes. **d**, Cell-cycle phases across matched cardiomyocyte states in adult and embryonic datasets. **e**, Expression of cardiomyocyte maturation markers across embryonic LV-CM subsets. **f,** Gene set enrichment analysis (GSEA) of embryonic cardiomyocyte states highlighting developmental, metabolic, and cell-cycle programs. **g,** Quantitative association of embryonic CM states and non-CM cell types with anatomical spatial domains, illustrating coordinated patterning of CM maturation with cardiac tissue architecture, as measured by scaled cell counts per compartment estimated by deconvolution **h,** Embryonic cell states mapped to Visium data (estimated counts per Visium spot) on a representative sample. **i,** PHLOWER trajectory with selected embryonic cardiomyocytes cells mapped onto adult CM branches, indicating immature LV-CMs to be related to the CFLAR branch.

**ED Data Figure 10.**
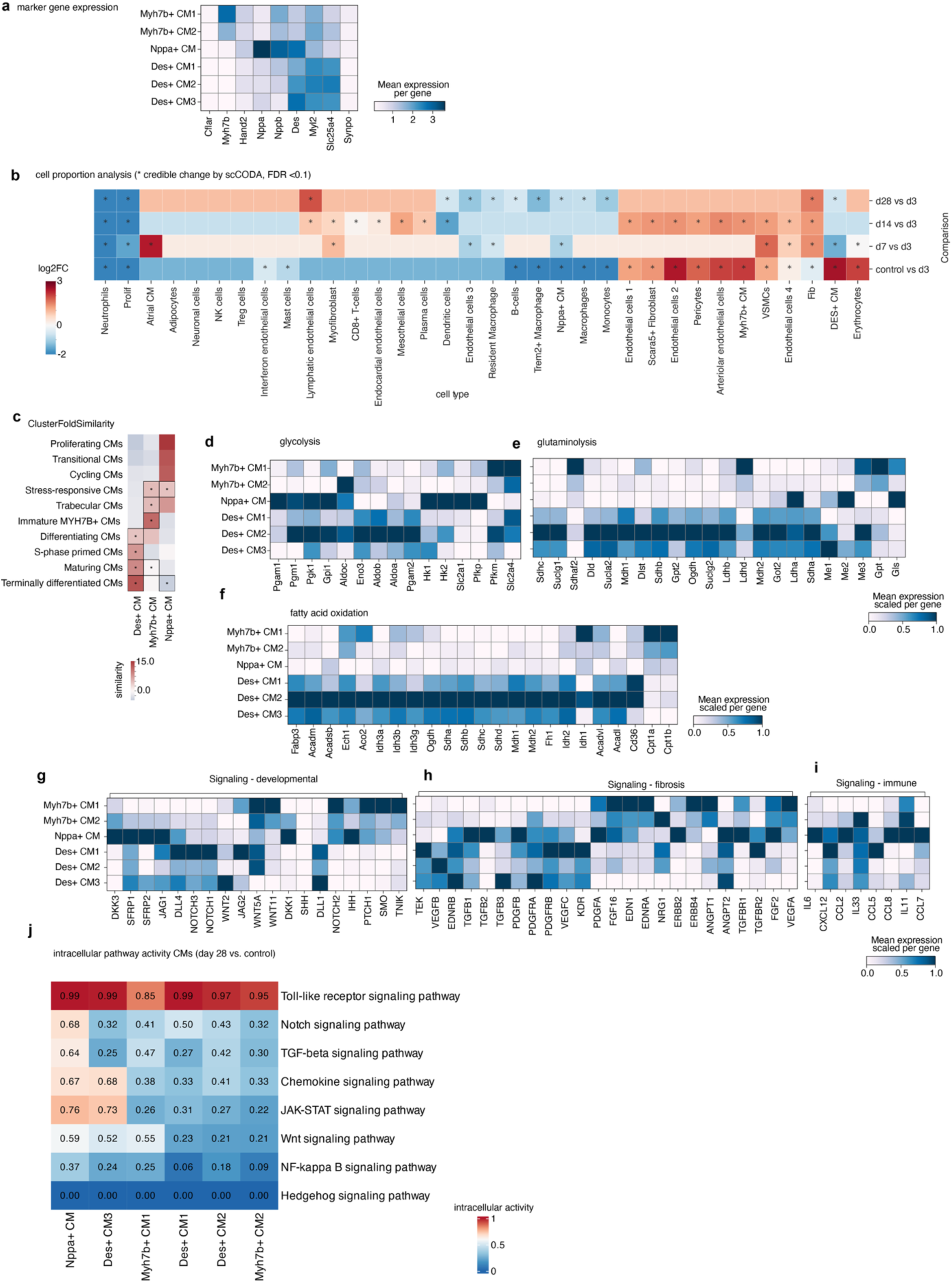
Cardiomyocyte state composition, metabolic rewiring, signaling programs, and spatial niches during disease progression in mice. **a,** Heatmap showing mean expression of selected cardiomyocyte (CM) marker genes across Myh7b⁺, Nppa⁺, and Des⁺ CM subclusters, confirming transcriptional identity of polyploid (Myh7b⁺, Nppa⁺) and diploid (Des⁺) CM states. Expression is scaled per gene. **b,** Differential cell proportion analysis across experimental conditions using scCODA. Heatmap indicates credible increases (blue) or decreases (red) in cell type abundance relative to control (FDR = 0.1). Asterisks denote statistically credible changes. Comparisons include control vs d3, d7 vs d3, d14 vs d3, and d28 vs d3. **c,** Enrichment of CM differentiation and cell cycle–related gene signatures across CM states. Polyploid Myh7b⁺ and Nppa⁺ CMs show enrichment of immature, stress-responsive, and cycling programs, whereas Des⁺ CMs align with terminal differentiation states. **d–f,** Heatmaps showing scaled mean expression of genes involved in **glycolysis** (d), **glutaminolysis** (e), and **fatty acid oxidation** (f) across CM subclusters. Polyploid CM states display increased glycolytic and glutaminolytic programs, while diploid Des⁺ CM states retain higher fatty acid oxidation signatures. **g–i,** Expression of signaling pathway components across CM states. **g,** Developmental signaling pathways (e.g., WNT, NOTCH, Hedgehog). **h,** Fibrosis-associated signaling (e.g., TGF-β, PDGF, VEGF, EGFR). **i,** Immune-related signaling and chemokine expression. Polyploid CM states preferentially activate developmental and pro-fibrotic signaling programs. **j**, Heatmap of intracellular pathway activity scores in cardiomyocyte (CM) subtypes at day 28 post-injury compared with controls. Pathways include Toll-like receptor, Notch, TGF-β, chemokine, JAK–STAT, Wnt, NF-κB, and Hedgehog signaling. Values represent normalized pathway activity, with clustering revealing differential inflammatory and developmental signaling across CM states (Nppa⁺, Des⁺, and Myh7b⁺ subsets).

**Extended Data Fig. 11.**
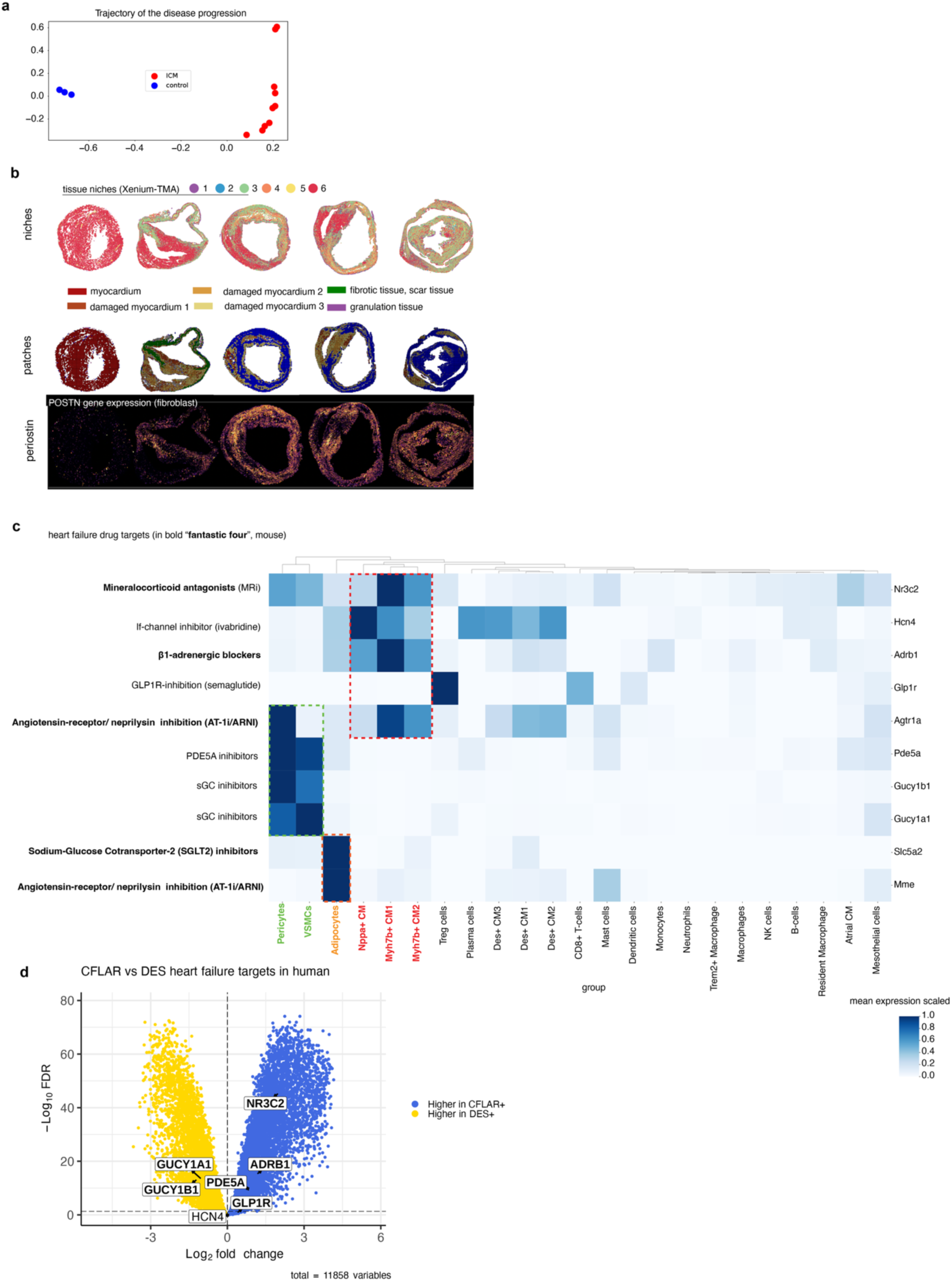
Spatial and transcriptional remodeling of cardiomyocyte states and therapeutic target engagement in ischemic cardiomyopathy. **a,** Low-dimensional embedding illustrating the trajectory of disease progression, separating control samples (blue) from ischemic cardiomyopathy (ICM; red), indicating a coordinated shift in global transcriptional states with disease. **b,** Spatial tissue organization inferred from five samples selected along the trajectory of **a**. Top: tissue niches (1–6) highlighting myocardium, damaged myocardium, fibrotic and scar tissue, and granulation tissue. Middle: spatial patches derived from unsupervised segmentation, reflecting recurrent micro-architectural patterns. Bottom: spatial expression of *Postn* in fibroblasts, illustrating fibrosis-associated gene expression across disease-affected regions. **c,** Cell type–resolved expression of major heart failure drug targets in mouse, including the “fantastic four” (β-blockers, MR antagonists, ARNI, and SGLT2 inhibitors; bold). Heatmap shows relative target gene expression across cardiomyocyte subtypes, vascular cells, adipocytes, and immune populations. Red dashed boxes highlight cardiomyocyte-enriched targets, while green dashed boxes indicate pericyte/VSMC-associated signaling nodes (e.g., PDE5A and sGC pathways), underscoring multicellular mechanisms of therapeutic action. **d**, Volcano plot between CFLAR+ and DES+ human CM. Highlighted are genes encoding heart failure targets. A line indicates the threshold of 0.05 FDR.

**Extended Data Fig. 12.**
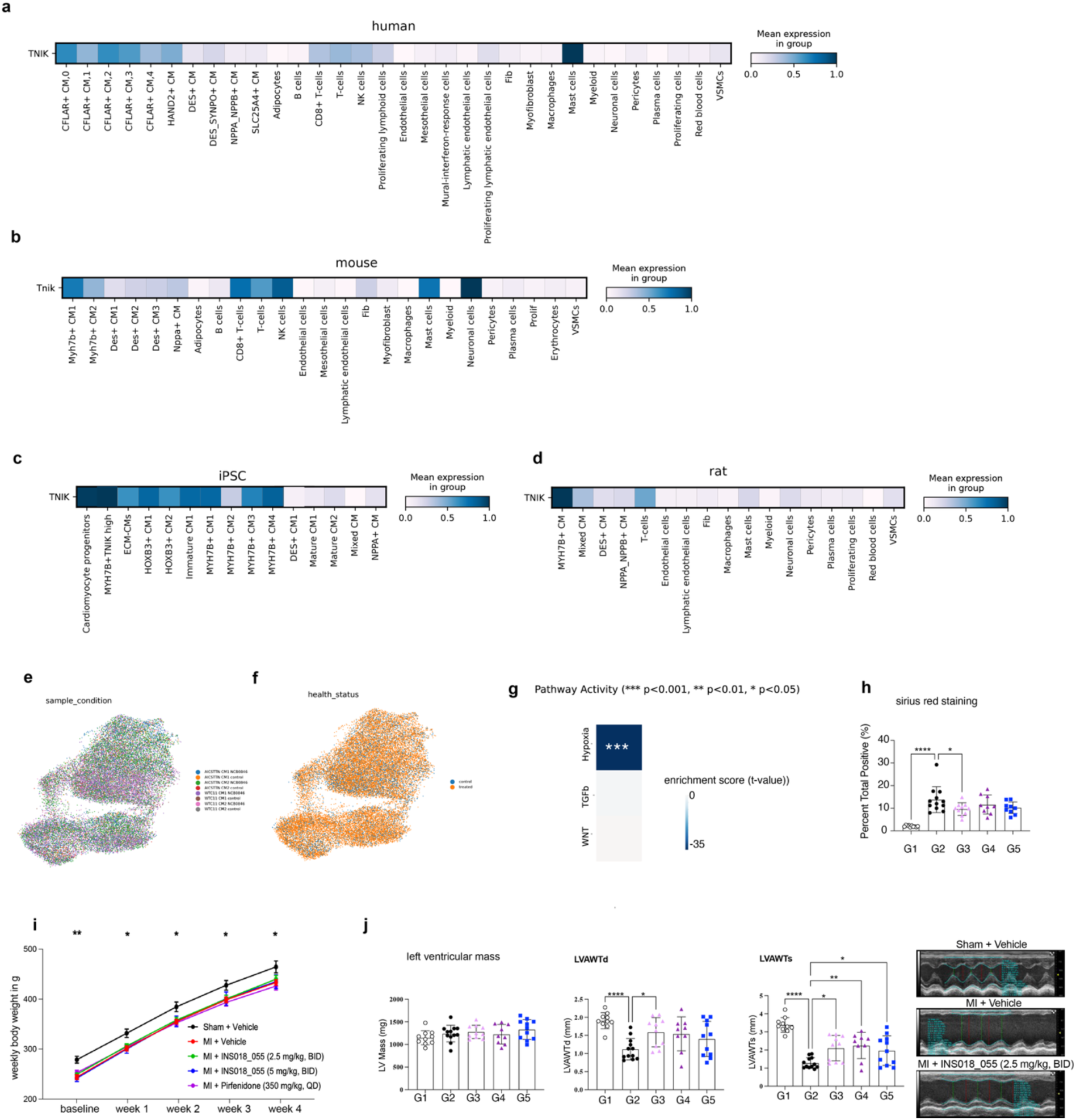
TNIK expression across species and models and functional effects of TNIK inhibition. **a,** Mean TNIK expression across human cardiac and non-cardiac cell types, highlighting enrichment in cardiomyocyte subpopulations, including CFLAR⁺ and HAND2⁺ cardiomyocytes, compared with stromal and immune compartments. **b,** Mean Tnik expression across mouse heart cell types, demonstrating conserved cardiomyocyte-biased expression relative to non-myocyte populations. **c,** TNIK expression across iPSC-derived cardiomyocyte differentiation stages and subtypes, with highest expression observed in immature and MYH7B⁺ cardiomyocyte states. **d,** Mean TNIK expression across rat cardiac cell populations, confirming cross-species conservation of cardiomyocyte-enriched TNIK expression. **e,** UMAP projection of iPSC cardiomyocytes colored by sample and experimental condition, showing robust integration across datasets. **f,** The same UMAP colored by health status, demonstrating overlap between control and diseased cardiomyocytes without condition-driven batch effects. **g,** Pathway enrichment scores inferred from cardiomyocyte transcriptomes of WTC11 and AICS-TTN iPSC-derived cardiomyocytes under control conditions or following TNIK inhibition (NCB-0846). *** p-value < 0.001 versus vehicle, two-sided Student’s t-test of the linear model slope **h**, Quantification of sirius red positive cells for all treatment groups **i,** Weekly body weight trajectories following myocardial infarction (MI) and treatment with vehicle, INS018_055 (two doses), or pirfenidone. TNIK inhibition does not induce weight loss compared with vehicle or antifibrotic control. Data are shown as mean ± s.e.m.; * p-value < 0.05, ** p-value < 0.01 versus MI + vehicle. **j,** Left ventricular mass, LVAWTd and LVAWTs for all treatments groups, Representative M-mode echocardiography images from sham-operated animals, MI + vehicle, and MI treated with INS018_055 (2.5 mg/kg, BID), illustrating improved cardiac contractile performance following TNIK inhibition.

