## Supplementary Information files for "Polyploid cardiomyocytes define disease-specific transcriptional states in the mammalian heart"

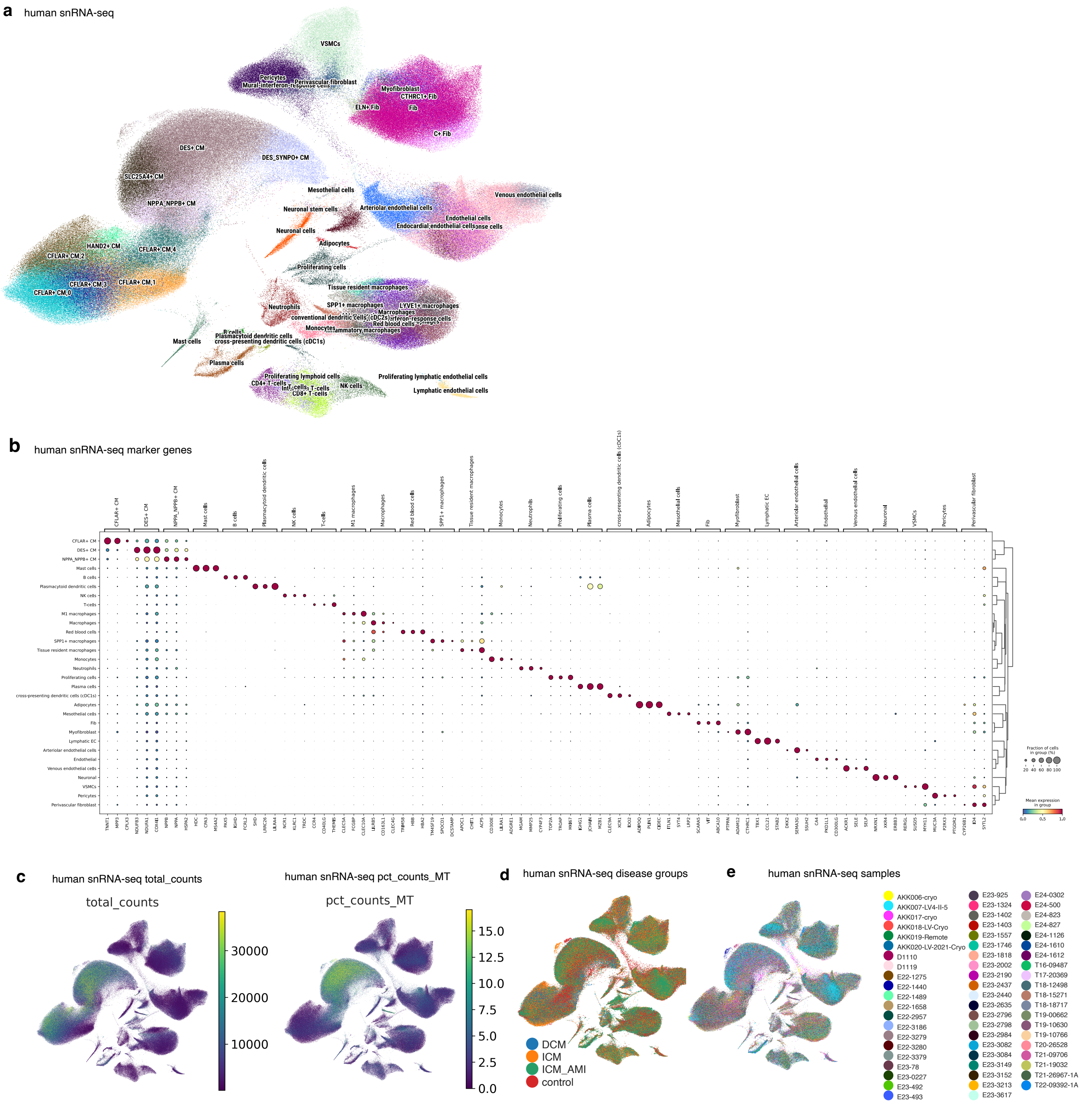
a
human snRNA-seq
VSMCs
Pericytes
Mural-interferon-response cells
Perivascular fibroblast
Myofibroblast
CTHRC1+ Fib
ELN+ Fib
Fib
C+ Fib
DES+ CM
DES_SYNPO+ CM
SLC25A4+ CM
NPPA_NPPB+ CM
Mesothelial cells
Neuronal stem cells
Arteriolar endothelial cells
Venous endothelial cells
Endothelial cells
Endocardial endothelial cells
Neuronal cells
Adipocytes
HAND2+ CM
CFLAR+ CM 4
CFLAR+ CM 2
CFLAR+ CM 3
CFLAR+ CM 1
CFLAR+ CM 0
Proliferating cells
Tissue resident macrophages
Neutrophils
SPP1+ macrophages
LYVE1+ macrophages
Macrophages
conventional dendritic cells (cDC2s)
Red blood cells
Monocytes
B cells
Plasmacytoid dendritic cells
cross-presenting dendritic cells (cDC1s)
Mast cells
Plasma cells
Proliferating lymphoid cells
CD4+ T-cells
T-cells
CD8+ T-cells
NK cells
Proliferating lymphatic endothelial cells
Lymphatic endothelial cells
b
human snRNA-seq marker genes
Fraction of cells in group (%)
20 40 60 80 100
Mean expression in group
0.0 0.5 1.0
c
human snRNA-seq total_counts
total_counts
30000
20000
10000
human snRNA-seq pct_counts_MT
pct_counts_MT
15.0
12.5
10.0
7.5
5.0
2.5
0.0
d
human snRNA-seq disease groups
DCM
ICM
ICM_AMI
control
e
human snRNA-seq samples
AKK006-cryo
AKK007-LV4-II-5
AKK017-cryo
AKK018-LV-Cryo
AKK019-Remote
AKK020-LV-2021-Cryo
D1110
D1119
E22-1275
E22-1440
E22-1489
E22-1658
E22-2957
E22-3186
E22-3279
E22-3280
E22-3379
E23-78
E23-0227
E23-492
E23-493
E23-925
E23-1324
E23-1402
E23-1403
E23-1557
E23-1746
E23-1818
E23-2002
E23-2190
E23-2437
E23-2440
E23-2635
E23-2796
E23-2798
E23-2984
E23-3082
E23-3084
E23-3149
E23-3152
E23-3213
E23-3617
E24-0302
E24-500
E24-823
E24-827
E24-1126
E24-1610
E24-1612
T16-09487
T17-20369
T18-12498
T18-15271
T18-18717
T19-00662
T19-10630
T19-10766
T20-26528
T21-09706
T21-19032
T21-26967-1A
T22-09392-1A

**Supplementary Data Figure 1 Overview and quality control of the human cardiac snRNA-seq data a,** Uniform Manifold Approximation and Projection (UMAP) of all nuclei profiled by single-nucleus RNA sequencing (snRNA-seq) from human cardiac tissue. Major cardiac cell types are annotated, including cardiomyocyte (CM) sub-states, fibroblast subsets, endothelial populations, vascular smooth muscle cells (VSMCs), immune cells, adipocytes, mesothelial cells, neuronal cells, and proliferating populations. **b,** Dot-plot showing canonical marker gene expression across annotated cardiac cell types identified in the human snRNA-seq dataset. Dot size represents the fraction of nuclei expressing the gene within each cell type, and color intensity indicates scaled average expression levels. Hierarchical clustering reflects transcriptional similarity between cell populations. **c,** Quality control metrics projected onto the UMAP embedding. Left: total UMI counts per nucleus (*total_counts*), illustrating sequencing depth distribution across cell populations. Right: percentage of mitochondrial transcripts (*pct_counts_MT*), indicating overall RNA integrity and low mitochondrial contamination across clusters. **d,** UMAP colored by disease group, including dilated cardiomyopathy (DCM), ischemic cardiomyopathy (ICM), ischemic cardiomyopathy with acute myocardial infarction (ICM_AMI), and non-failing control samples, demonstrating broad representation of disease states across all major cell types. **e,** UMAP colored by individual human samples, highlighting inter-sample integration and minimal batch effects following data harmonization.

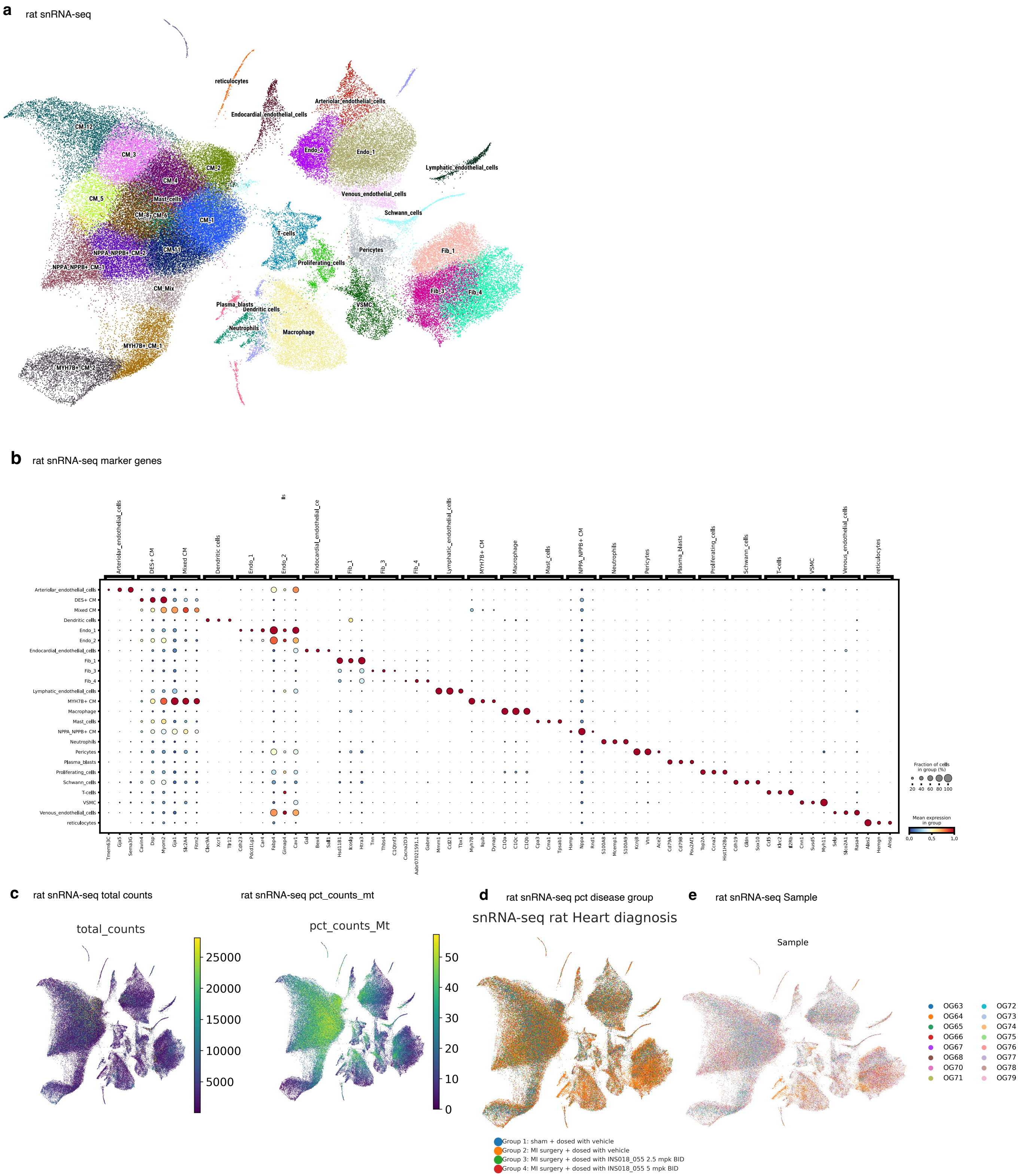
a
rat snRNA-seq
reticulocytes
Endocardial_endothelial_cells
Arteriolar_endothelial_cells
CM_12
CM_3
CM_2
CM_4
CM_5
Mast_cells
CM_8
CM_6
CM_1
Endo_2
Endo_1
Lymphatic_endothelial_cells
Venous_endothelial_cells
Schwann_cells
T-cells
Pericytes
Proliferating_cells
NPPA_NPPB+ CM_2
CM_11
NPPA_NPPB+ CM_1
CM_Mix
Fib_1
Fib_3
Fib_4
VSMC
Plasma_blasts
Dendritic cells
Neutrophils
Macrophage
MYH7B+ CM_1
MYH7B+ CM_2
b
rat snRNA-seq marker genes
Fraction of cells in group (%)
20 40 60 80 100
Mean expression in group
0.0 0.5 1.0
c
rat snRNA-seq total counts
total_counts
25000
20000
15000
10000
5000
rat snRNA-seq pct_counts_mt
pct_counts_Mt
50
40
30
20
10
0
d
rat snRNA-seq pct disease group
snRNA-seq rat Heart diagnosis
Group 1: sham + dosed with vehicle
Group 2: MI surgery + dosed with vehicle
Group 3: MI surgery + dosed with INS018_055 2.5 mpk BID
Group 4: MI surgery + dosed with INS018_055 5 mpk BID
e
rat snRNA-seq Sample
Sample
OG63
OG64
OG65
OG66
OG67
OG68
OG70
OG71
OG72
OG73
OG74
OG75
OG76
OG77
OG78
OG79

**Supplementary Data Figure 2 Single-nucleus RNA-seq dataset and quality control rat a,** Uniform Manifold Approximation and Projection (UMAP) of all nuclei profiled by single-nucleus RNA sequencing (snRNA-seq) from rat cardiac tissue. Major cardiac cell populations are annotated, including multiple cardiomyocyte (CM) sub-states, fibroblast subsets, endothelial cell types, vascular smooth muscle cells (VSMCs), pericytes, Schwann cells, immune populations, proliferating cells, and reticulocytes. **b,** Dot-plot showing expression of canonical marker genes used to annotate rat cardiac cell populations. Dot size represents the fraction of nuclei expressing each gene within a given cell type, while color intensity indicates scaled mean expression. Hierarchical clustering reflects transcriptional similarity among annotated cell types. **c,** Quality control metrics projected onto the UMAP embedding. Left: total UMI counts per nucleus (*total_counts*), illustrating sequencing depth across rat cardiac cell populations. Right: percentage of mitochondrial transcripts (*pct_counts_mt*), indicating RNA quality and low levels of mitochondrial contamination across clusters. **d,** UMAP colored by experimental group, including sham-operated controls and myocardial injury groups with vehicle or pharmacological intervention, demonstrating balanced representation of experimental conditions across cell populations. **e,** UMAP colored by individual rat samples, highlighting inter-sample integration and minimal batch effects following data harmonization.

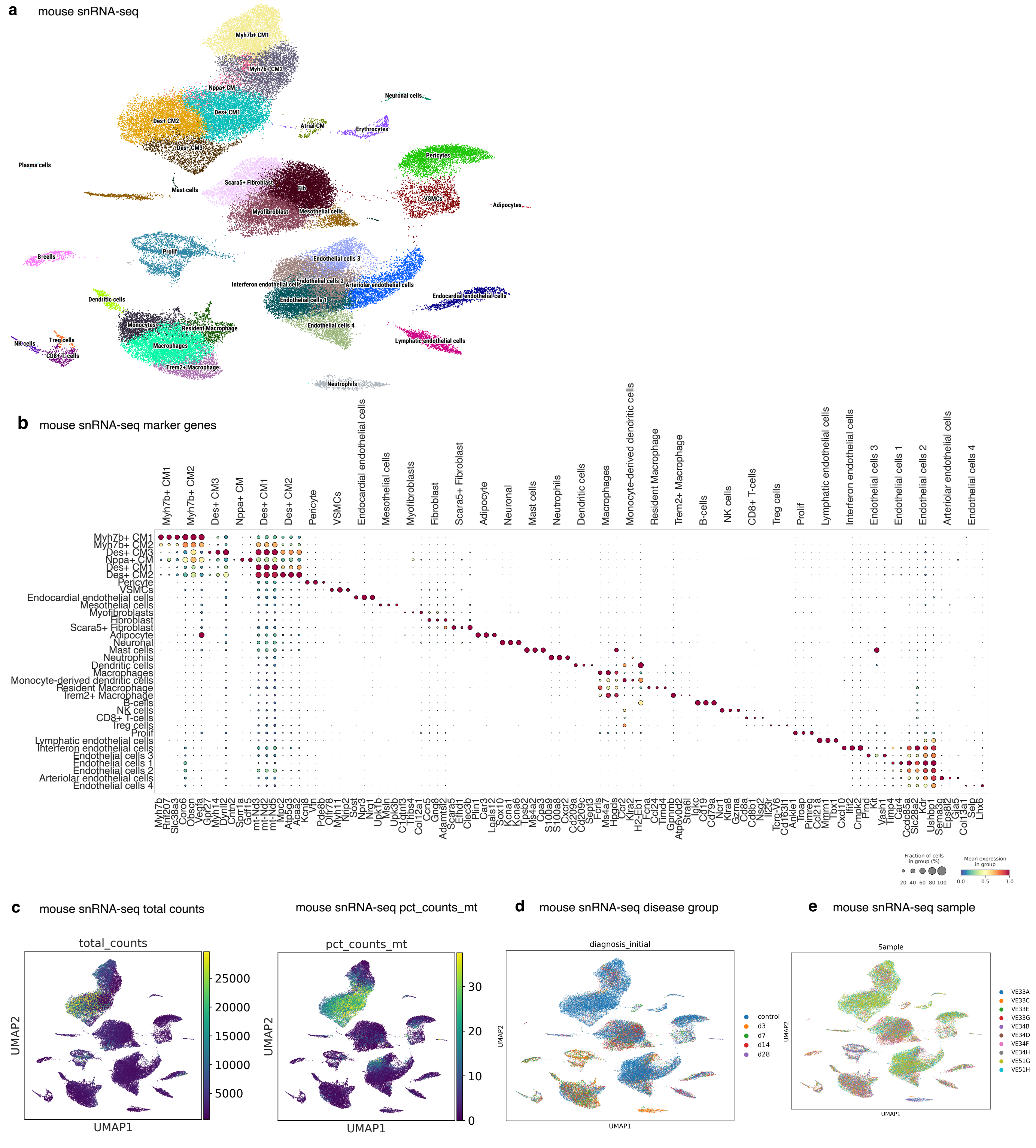
a
mouse snRNA-seq
Myh7b+ CM1
Myh7b+ CM2
Nppa+ CM
Des+ CM1
Des+ CM2
Des+ CM3
Atrial CM
Neuronal cells
Erythrocytes
Pericytes
VSMCs
Adipocytes
Plasma cells
Mast cells
Scara5+ Fibroblast
Fib
Myofibroblast
Mesothelial cells
Prolif
B-cells
Endothelial cells 3
Interferon endothelial cells
Endothelial cells 2
Arteriolar endothelial cells
Endothelial cells 1
Endocardial endothelial cells
Endothelial cells 4
Dendritic cells
Monocytes
Resident Macrophage
Macrophages
Trem2+ Macrophage
NK cells
Treg cells
CD8+ T-cells
Lymphatic endothelial cells
Neutrophils
b
mouse snRNA-seq marker genes
Myh7b+ CM1
Myh7b+ CM2
Des+ CM3
Nppa+ CM
Des+ CM1
Des+ CM2
Pericyte
VSMCs
Endocardial endothelial cells
Mesothelial cells
Myofibroblasts
Fibroblast
Scara5+ Fibroblast
Adipocyte
Neuronal
Mast cells
Neutrophils
Dendritic cells
Macrophages
Monocyte-derived dendritic cells
Resident Macrophage
Trem2+ Macrophage
B-cells
NK cells
CD8+ T-cells
Treg cells
Prolif
Lymphatic endothelial cells
Interferon endothelial cells
Endothelial cells 3
Endothelial cells 1
Endothelial cells 2
Arteriolar endothelial cells
Endothelial cells 4
Fraction of cells in group (%)
20 40 60 80 100
Mean expression in group
0.0 0.5 1.0
c
mouse snRNA-seq total counts
total_counts
25000
20000
15000
10000
5000
UMAP2
UMAP1
mouse snRNA-seq pct_counts_mt
pct_counts_mt
30
20
10
0
UMAP2
UMAP1
d
mouse snRNA-seq disease group
diagnosis_initial
control
d3
d7
d14
d28
UMAP2
UMAP1
e
mouse snRNA-seq sample
Sample
VE33A
VE33C
VE33E
VE33G
VE34B
VE34D
VE34F
VE34H
VE51G
VE51H
UMAP2
UMAP1

**Supplementary Data Figure 3 Single-nucleus RNA-seq dataset and quality control mouse a,** Uniform Manifold Approximation and Projection (UMAP) of all nuclei profiled by single-nucleus RNA sequencing (snRNA-seq) from rat cardiac tissue. Major cardiac cell populations are annotated, including multiple cardiomyocyte (CM) sub-states, fibroblast subsets, endothelial cell types, vascular smooth muscle cells (VSMCs), pericytes, Schwann cells, immune populations, proliferating cells, and reticulocytes. **b,** Dot-plot showing expression of canonical marker genes used to annotate rat cardiac cell populations. Dot size represents the fraction of nuclei expressing each gene within a given cell type, while color intensity indicates scaled mean expression. Hierarchical clustering reflects transcriptional similarity among annotated cell types. **c,** Quality control metrics projected onto the UMAP embedding. Left: total UMI counts per nucleus (*total_counts*), illustrating sequencing depth across rat cardiac cell populations. Right: percentage of mitochondrial transcripts (*pct_counts_mt*), indicating RNA quality and low levels of mitochondrial contamination across clusters. **d,** UMAP colored by experimental group and treatment condition, including sham-operated controls and myocardial injury groups with vehicle or pharmacological intervention, demonstrating balanced representation of experimental conditions across cell populations.**e,** UMAP colored by individual mouse samples, highlighting inter-sample integration and minimal batch effects following data harmonization.

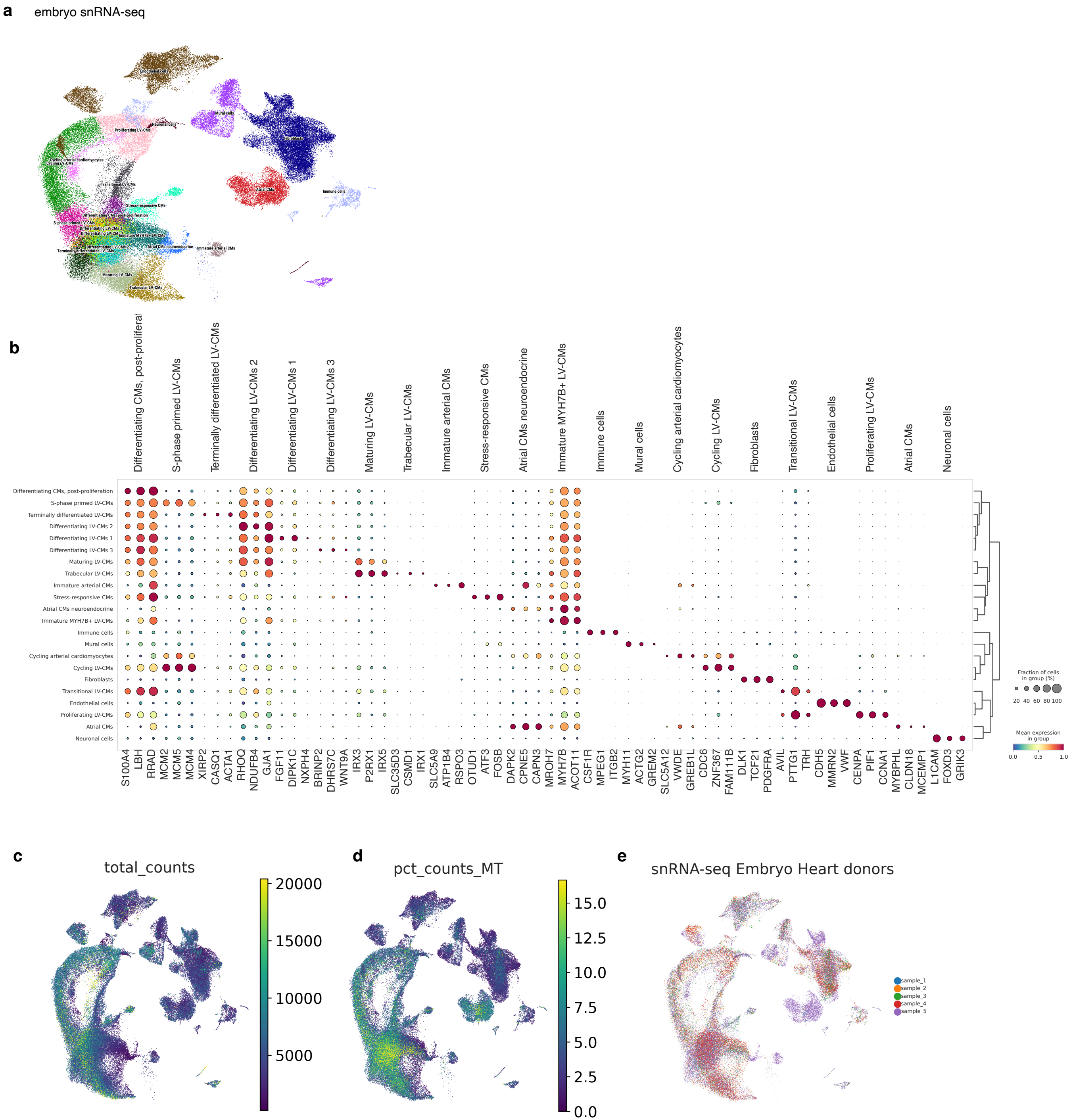
a
embryo snRNA-seq
Endothelial cells
Mural cells
Neuronal cells
Proliferating LV-CMs
Fibroblasts
Cycling arterial cardiomyocytes
Cycling LV-CMs
Transitional LV-CMs
Atrial CMs
Immune cells
Stress-responsive CMs
Differentiating CMs, post-proliferation
S-phase primed LV-CMs
Differentiating LV-CMs 3
Differentiating LV-CMs 1
Immature MYH7B+ LV-CMs
Differentiating LV-CMs 2
Atrial CMs neuroendocrine
Immature arterial CMs
Terminally differentiated LV-CMs
Maturing LV-CMs
Trabecular LV-CMs
b
Differentiating CMs, post-proliferat
S-phase primed LV-CMs
Terminally differentiated LV-CMs
Differentiating LV-CMs 2
Differentiating LV-CMs 1
Differentiating LV-CMs 3
Maturing LV-CMs
Trabecular LV-CMs
Immature arterial CMs
Stress-responsive CMs
Atrial CMs neuroendocrine
Immature MYH7B+ LV-CMs
Immune cells
Mural cells
Cycling arterial cardiomyocytes
Cycling LV-CMs
Fibroblasts
Transitional LV-CMs
Endothelial cells
Proliferating LV-CMs
Atrial CMs
Neuronal cells
Differentiating CMs, post-proliferation
S-phase primed LV-CMs
Terminally differentiated LV-CMs
Differentiating LV-CMs 2
Differentiating LV-CMs 1
Differentiating LV-CMs 3
Maturing LV-CMs
Trabecular LV-CMs
Immature arterial CMs
Stress-responsive CMs
Atrial CMs neuroendocrine
Immature MYH7B+ LV-CMs
Immune cells
Mural cells
Cycling arterial cardiomyocytes
Cycling LV-CMs
Fibroblasts
Transitional LV-CMs
Endothelial cells
Proliferating LV-CMs
Atrial CMs
Neuronal cells
S100A4 LBH RRAD MCM2 MCM5 MCM4 XIRP2 CASQ1 ACTA1 RHOQ NDUFB4 GJA1 FGF11 DIPK1C NXPH4 BRINP2 DHRS7C WNT9A IRX3 P2RX1 IRX5 SLC35D3 CSMD1 IRX1 SLC5A9 ATP1B4 RSPO3 OTUD1 ATF3 FOSB DAPK2 CPNE5 CAPN3 MROH7 MYH7B ACOT11 CSF1R MPEG1 ITGB2 MYH11 ACTG2 GREM2 SLC5A12 VWDE GREB1L CDC6 ZNF367 FAM111B DLK1 TCF21 PDGFRA AVIL PTTG1 TRH CDH5 MMRN2 VWF CENPA PIF1 CCNA1 MYBPHL CLDN18 MCEMP1 L1CAM FOXD3 GRIK3
Fraction of cells in group (%)
20 40 60 80 100
Mean expression in group
0.0 0.5 1.0
c
total_counts
20000
15000
10000
5000
d
pct_counts_MT
15.0
12.5
10.0
7.5
5.0
2.5
0.0
e
snRNA-seq Embryo Heart donors
sample_1
sample_2
sample_3
sample_4
sample_5

**Supplementary Figure 4 Single-nucleus RNA-seq dataset and quality control human embryo a,** Uniform Manifold Approximation and Projection (UMAP) of all nuclei profiled by single-nucleus RNA sequencing (snRNA-seq) from human embryo cardiac tissue. **b,** Dot-plot showing expression of canonical marker genes used to annotate rat cardiac cell populations. Dot size represents the fraction of nuclei expressing each gene within a given cell type, while color intensity indicates scaled mean expression. **c,** Quality control metrics projected onto the UMAP embedding. Total UMI counts per nucleus (*total_counts*), illustrating sequencing depth across rat cardiac cell populations. **d,** Percentage of mitochondrial transcripts (*pct_counts_mt*), indicating RNA quality and low levels of mitochondrial contamination across clusters. **e,** UMAP colored by snRNA-seq sample of the embryo human hearts.
